# Feature-based encoding of face identity by single neurons in the human amygdala and hippocampus

**DOI:** 10.1101/2020.09.01.278283

**Authors:** Runnan Cao, Jinge Wang, Chujun Lin, Emanuela De Falco, Alina Peter, Hernan G. Rey, James DiCarlo, Alexander Todorov, Ueli Rutishauser, Xin Li, Nicholas J. Brandmeir, Shuo Wang

**Affiliations:** Department of Chemical and Biomedical Engineering, West Virginia University, Morgantown, WV 26506, USA; Lane Department of Computer Science and Electrical Engineering, West Virginia University, Morgantown, WV 26506, USA; Department of Psychological and Brain Sciences, Dartmouth College, Hanover, NH 03755, USA; Laboratory of Cognitive Neuroscience, École Polytechnique Fédérale de Lausanne, Route Cantonale, Lausanne, 1015, Switzerland; Department of Brain and Cognitive Sciences, Massachusetts Institute of Technology, Cambridge, MA 02139, USA; Department of Neurosurgery, Baylor College of Medicine, Houston, TX 77030, USA; Booth School of Business, University of Chicago, Chicago, IL 60637; Departments of Neurosurgery and Neurology, Cedars-Sinai Medical Center, Los Angeles, CA 90048, USA; Department of Neurosurgery, West Virginia University, Morgantown, WV 26506, USA; Rockefeller Neurosciences Institute, West Virginia University, Morgantown, WV 26506, USA; Department of Radiology, Washington University in St. Louis, St. Louis, MO 63110, USA

**Keywords:** Human single-neuron recordings, Amygdala, Hippocampus, Face, Deep neural network, Identity neuron, Feature coding

## Abstract

Neurons in the human amygdala and hippocampus that are selective for the identity of specific people are classically thought to encode a person’s identity invariant to visual features (e.g., skin tone, eye shape). However, it remains largely unknown how visual information from higher visual cortical areas is translated into such a semantic representation of an individual person. Here, we show that some amygdala and hippocampal neurons are selective to multiple different unrelated face identities based on shared visual features. The encoded identities form clusters in the representation of a deep neural network trained to recognize faces. Contrary to prevailing views, these neurons thus represent an individual’s face with a visual feature-based code rather than one based on association with known concepts. Feature neurons encoded faces regardless of their identity, race, gender, familiarity, or pixel-level visual features; and the region of feature space to which feature neurons are tuned predicted their response to new face stimuli. Our results reveal a new class of neurons that bridge the perception-driven representation of facial features in the higher visual cortex with mnemonic semantic representations in the MTL, which may form the basis for declarative memory.

## Main Text

How the human brain encodes and stores in memory different face identities is one of the most fundamental and intriguing questions in neuroscience. There are two extreme hypotheses. The *feature-based model* posits that face representations are encoded over a broad and distributed population of neurons ^1–4^. Under this model, recognizing a particular individual requires access to many neurons, with each neuron responding to many different faces that share specific visual features such as shape and skin tone ^5,6^. Evidence for a specific type of feature-based coding, axis-based feature coding, has been demonstrated in the non-human primate inferotemporal (IT) cortex ^7–10^ and in humans using fMRI ^11–13^. That is, neurons/voxels parametrically correlate with facial features along specific axes in face space. On the other extreme, the *exemplar-based model* posits that explicit facial representations in the brain are formed by highly selective (sparse) but at the same time highly visually invariant neurons ^14–17^. Identity neurons that selectively respond to many different images of a specific person’s face embody exemplar-based coding. Such neurons exist in the human amygdala and hippocampus ^16,17^ and are thought to constitute the building blocks for episodic memories ^17^. The responses of identity neurons are clustered by high-level conceptual or semantic relatedness (e.g., Bill Clinton and Hillary Clinton) rather than by facial features ^18–20^.

Feature-based and exemplar-based models of face processing are not mutually exclusive, with the former thought to give rise to the latter. However, the neural computations bridging these two mechanisms remain little understood. Here, we identify a key missing link between the representation of specific facial features *feature-based coding*) and the representation of specific people (*exemplar-based coding*) in the amygdala and hippocampus. We show that a subset of neurons in the human amygdala and hippocampus carry a novel *region-based feature code* for face identity that forms an intermediate code linking between *feature-based coding* and *exemplar-based coding*.

## Results

### Identity neurons

We recorded from 2082 neurons in the amygdala and hippocampus (together referred to as Medial Temporal lobe [MTL]) of 12 neurosurgical patients (4 male; 38 sessions in total; **Extended Data Table 1; Extended Data Fig. 1**) while they performed a one-back task (**Extended Data Fig. 2a**; accuracy = 75.2%±20.0% [mean±SD across sessions]). Participants viewed 500 natural face pictures of 50 celebrities (**Extended Data Fig. 2b**; 10 different pictures per celebrity/identity). 1577 neurons had an overall firing rate greater than 0.15 Hz and we restricted our analysis to this subset of neurons, which included 753 neurons from the amygdala, 505 neurons form the anterior hippocampus, and 319 neurons from the posterior hippocampus (**Extended Data Table 1**). Below, we first delineate the response characteristics of neurons that encode face identities ^16,18,21^. Then we show a broader class of feature neurons and validate and generalize feature coding using new stimuli. Lastly, we compare feature coding models with further recordings from a monkey.

To select *identity neurons*, we first used a one-way ANOVA to identify neurons with a significantly unequal response to different identities (P < 0.05) in a window 250-1250 ms following stimulus onset. We next imposed an additional criterion to identify which identities a neuron was selectively responding to (selected identities): the neural response to such an identity was required to be at least 2 standard deviations (SD) above the mean of neural responses from all identities. We found 155 identity neurons (9.83%, binomial P = 1.67×10^-15^; **Fig. 1a, d; Extended Data Fig. 2c**; **Extended Data Table 1**), consistent with prior recordings from the human MTL ^16,18,21^. Of the 155 identity neurons, 53 neurons responded to a single identity (referred to here as *single-identity [S-ID] neurons* ^16,21^) and the remaining 102 neurons each responded to multiple identities (referred to here as *multiple-identity [M-ID] neurons* ^18,19^).

**Fig. 1.**
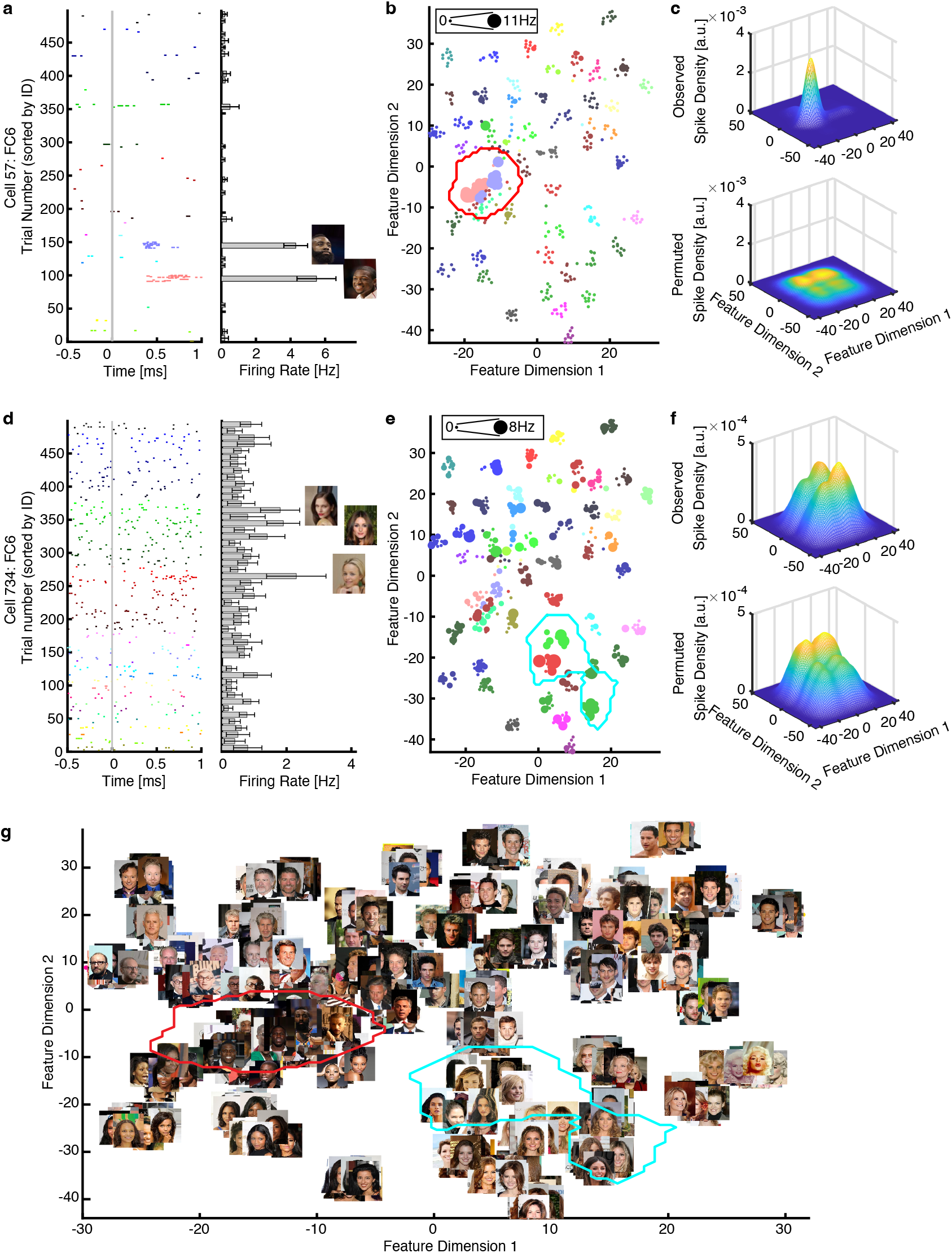
Feature-based neuronal coding of face identities. **(a-f)** Two example neurons that encoded visually similar identities (i.e., feature M-ID neurons). **(a, d)** Neuronal responses to 500 faces (50 identities). Trials are aligned to face stimulus onset (gray line) and are grouped by individual identity. Error bars denote ±SEM across faces. **(b, e)** Projection of the firing rate onto the FC6 feature space. Each color represents a different identity. The size of the dot indicates the firing rate. **(c, f)** Estimate of the spike density in the feature space. By comparing observed (upper) vs. permuted (lower) responses, we could identify a region where the observed neuronal response was significantly higher in the feature space. This region was defined as the tuning region of a neuron (delineated by the red/cyan outlines; also shown in **(g)**). **(g)** The face feature space constructed by t-distributed stochastic neighbor embedding (t-SNE) for the DNN layer FC6. All stimuli are shown in this space.

On average, M-ID neurons encoded 2.48±0.61 identities. We confirmed the results using an identity selective index (*d*’ between the most- and least-preferred identities; **Extended Data Fig. 2d**) and ordered responses from the most- to the least-preferred identities (**Extended Data Fig. 2e**). As expected, S-ID neurons had a sharp decrease of response from the most-preferred identity while M-ID neurons showed constantly steeper changes from the most- to the least-preferred identity compared to the non-identity neurons (**Extended Data Fig. 2e**). This was also captured by the larger decrease of firing rate between the most-preferred and the second most-preferred stimuli in S-ID neurons compared to M-ID neurons (**Extended Data Fig. 2f**; two-tailed unpaired *t*-test: *t*(153) = 4.64, P = 0.0007). We further confirmed the results using a depth of selectivity (DOS) index (**Extended Data Fig. 2g**) and single-trial population decoding (**Extended Data Fig. 2h, i**), which showed that it was possible to predict the identity of the face shown (see also **Supplementary Information** and **Extended Data Fig. 2j** for separate analysis for familiar vs. unfamiliar faces as judged by participants).

### Feature-based coding of face identities: feature identity neurons

It has been shown that some M-ID neurons encode *conceptually* related identities (e.g., Bill Clinton and Hillary Clinton) ^18–20^ in a way that makes the response of M-ID neurons invariant to visual features ^16,17,21^. However, it is unknown whether M-ID neurons can also encode *visually* (rather than conceptually) similar identities (i.e., identities sharing a similar visual appearance, e.g., a similar face and eye shape, skin or hair color). To answer this question, we extracted facial features from the pictures shown to the patients using a pre-trained deep neural network (DNN) VGG-16 trained to recognize faces (see **Extended Data Fig. 3a, b** for DNN architecture and visualization of DNN features). Facial features were coded by the weights and activation of thousands of DNN units (i.e., the DNN features), and we therefore used the DNN features to represent facial features. We further reduced the dimensionality of the DNN features and constructed a two-dimensional face feature space using t-distributed stochastic neighbor embedding (t-SNE) for each DNN layer (**Fig. 1b, e, g** and **Extended Data Fig. 4**; note that quantifications are in this t-SNE space but we replicated our results in the full dimensional space of the DNN; also note that the pairwise distance between faces in the full dimensional space is preserved in the t-SNE space as shown in **Extended Data Fig. 3d**). The dimensions (or axes) of the face feature space represented the major variations in faces that led to successful recognition of the identities by the DNN. Note that this face feature space was derived solely from the input pictures without knowledge of the patients’ neural responses and/or tuning of neurons (the aspect of a stimulus to which a neuron responds).

The feature space demonstrated an organized structure. For example, faces of the same identity were clustered in the first fully connected (FC) layer FC6 (which is towards the top of the DNN hierarchy and demonstrates clustering of identities; **Extended Data Fig. 3** and **Extended Data Fig. 4**), where Feature Dimension 2 represented a gender dichotomy, and darker skinned faces were clustered at the bottom left corner of the feature space (**Fig. 1g**). Note that the DNN had no access to semantic information about the faces (e.g., gender, ethnicity, social traits), and therefore, the representation of each face in the feature space was entirely driven by its visual appearance. Thus, faces were distributed in the DNN-derived feature space purely based on their visual appearance, regardless of any semantic information or conceptual association with each other.

We next projected the neuronal responses of a given neuron to each face onto this DNN-derived face feature space (i.e., multiplying the firing rate of each face to its corresponding location in the feature space to derive a response-weighted 2D feature map; **Fig. 1b, e**). Strikingly, this revealed that some M-ID neurons were selective to different identities that were clustered together in the face feature space (**Fig. 1b, e;** see **Extended Data Fig. 5** for more examples). This suggests that these M-ID neurons responded to face identities that were visually similar (e.g., similar in face shape or skin tone). To formally quantify the tuning of M-ID neurons, we estimated a continuous spike density map in the VGG-derived 2D feature space from our sparse sampling (**Fig. 1c, f** upper) and used a permutation test (1000 runs; **Fig. 1c, f** lower) to identify the region that had a significantly higher spike density above chance (red/cyan outlines in **Fig. 1b, e, g**; significant pixels were selected with permutation P < 0.01 and cluster size thresholds; see **Methods** and **Extended Data Fig. 6** for illustration of the selection procedure). This region shows the part of the feature space to which a neuron was tuned; and the significant neurons demonstrated *region-based feature coding* because they coded a certain region in the feature space. At the population level, we found that for 42/102 M-ID neurons (41.2%), all their selected identities were clustered in the feature space (referred to here as *feature M-ID neurons;* **Extended Data Fig. 2c**; the other M-ID neurons encoded identities distributed in the feature space and are referred to as *non-feature M-ID neurons*). Therefore, feature M-ID neurons encoded identities sharing similar facial features and thus looking visually similar.

In the DNN, the level of feature abstraction, and thus clustering of identities, increases from earlier layers to later layers (**Extended Data Fig. 3** and **Extended Data Fig. 4**). We therefore expect feature M-ID neurons best reflect the face features represented in later DNN layers. Indeed, we observed feature M-ID neurons in DNN layers Pool5 (25 neurons), FC6 (27 neurons), FC7 (30 neurons), and FC8 (30 neurons; some neurons appeared in multiple layers given the distribution of identities across layers; **Extended Data Fig. 4**; see **Extended Data Fig. 7a, d** for a breakdown of amygdala and hippocampal neurons). The tuning region of an individual feature M-ID neuron covered approximately 1.82-9.82% of the 2D feature space, with a similar mean coverage across layers (**Fig. 2a**; note that when we calculated the tuning region, we adjusted the kernel size to be proportional to the feature dimensions such that the percentage of space coverage was independent of the actual size of the feature space). In contrast, the response of an individual S-ID or non-feature M-ID neuron covered a significantly smaller region in the feature space (**Fig. 2a**; two-tailed unpaired *t*-test: P < 0.001 for all comparisons). This result was as expected because the identities (and thus the tuning regions) encoded by non-feature M-ID neurons were not contiguous with each other and were further apart (**Fig. 2b**). As a whole, the neuronal population that we sampled covered approximately 46-52% of the feature space (**Fig. 2c**; some areas were encoded by multiple neurons), and the tuning regions of the non-feature M-ID neurons were more distributed compared with feature M-ID neurons (**Fig. 2c**). The distribution of pairwise distance between faces within each neuron’s tuning region(s) further showed that feature M-ID neurons had a single large tuning region whereas non-feature M-ID neurons had smaller tuning regions that were more widely distributed across the feature space (**Fig. 2d**).

**Fig. 2.**
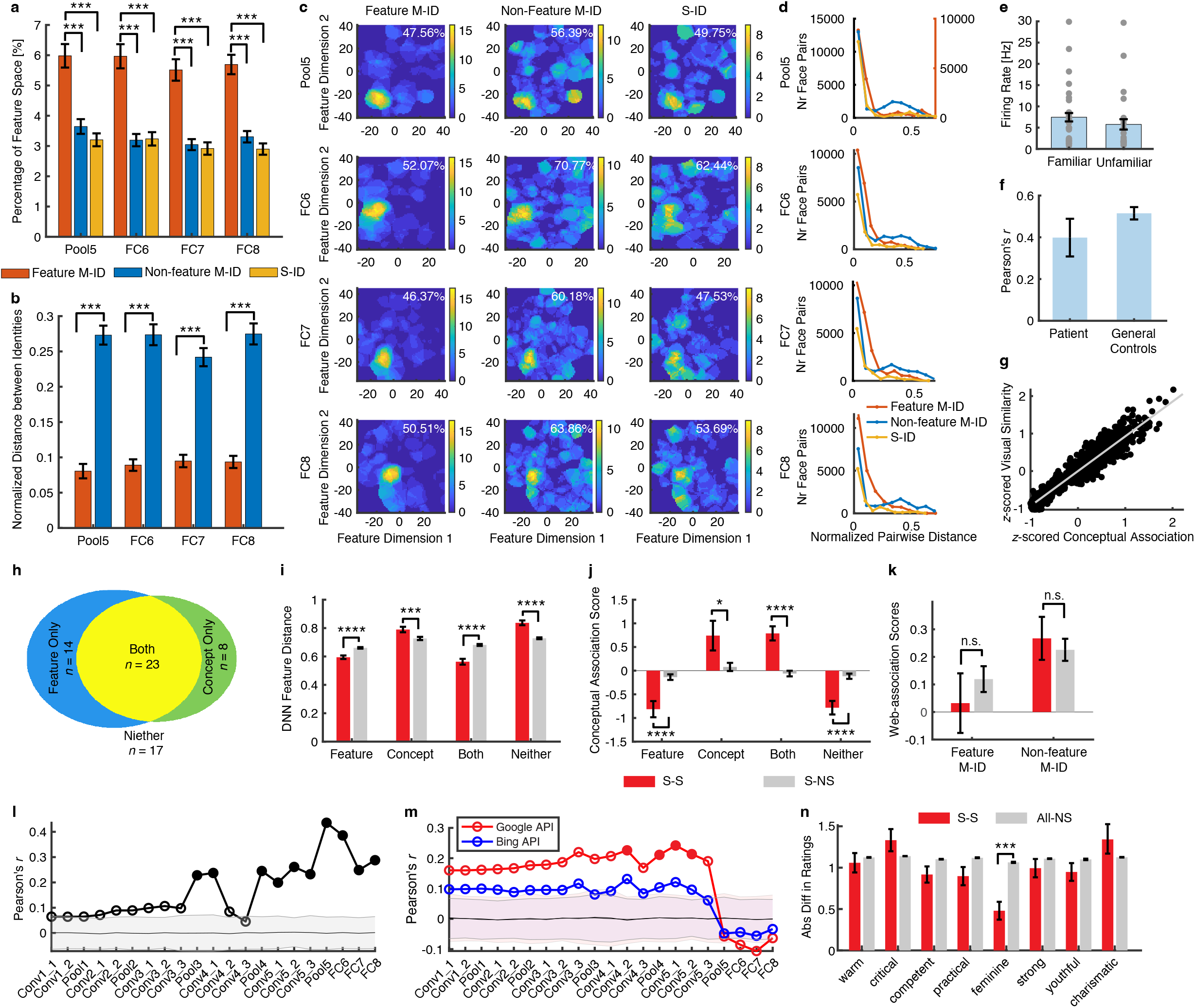
Summary of feature tuning for identity neurons and comparison between visual similarity and conceptual association. **(a)** Percentage of feature space covered by tuning regions of identity neurons. Note that here we did not apply the threshold for minimum cluster size for S-ID and nonfeature M-ID neurons in order to compare across different categories of identity neurons. **(b)** Normalized distance between M-ID neurons’ selected identities in the feature space. To be comparable for different layers, Euclidean distance was normalized by the maximum distance (i.e., diagonal line) of the feature space per layer. Error bars denote ±SEM across neurons. Asterisks indicate a significant difference between feature M-ID neurons and non-feature M-ID neurons using two-tailed unpaired *t*-test. *: P < 0.05, **: P < 0.01, and ***: P < 0.001. **(c)** The aggregated tuning regions of the neuronal population. Color bars show the counts of overlap between individual tuning regions. Numbers in the density map show the percentage of feature space covered by the tuning regions of the total observed neuronal population. **(d)** Distribution of pairwise distance between faces in each neuron’s tuning region(s). Euclidean distance was normalized by the maximum distance of the feature space. S-ID: *n* = 53 for all layers. Feature MID: *n* = 25 for Pool5, *n* = 27 for FC6, *n* = 30 for FC7, and *n* = 30 for FC8. Non-feature M-ID: *n* = 77 for Pool5, *n* = 75 for FC6, *n* = 72 for FC7, and *n* = 72 for FC8. **(e)** Feature M-ID neurons did not differentiate familiar vs. unfamiliar selected identities (two-tailed paired *t*-test, P > 0.05). Each dot represents a neuron. Error bars denote ±SEM across neurons. **(f)** Correlation between conceptual association ratings and visual similarity ratings. Error bars denote ±SEM across participants (*n* = 5 for patients and *n* = 40 for general controls). **(g)** Correlation between conceptual association ratings and visual similarity ratings across pairs of face identities. Ratings from the general controls were averaged across participants to derive consensus ratings. Each dot represents a pair of face identities (*n* = 1225) and the gray line denotes the linear fit. **(h)** Separate populations of M-ID neurons encoding visual features (i.e., visual similarity) and concepts (i.e., conceptual association). By selecting conceptual association neurons from M-ID neurons (i.e., comparing conceptual association ratings between encoded vs. unselected identities) from patients who provided ratings (*n* = 62), we found a subset of neurons that only encoded visual features (*n* = 14; note that to be comparable with the selection of conceptual association neurons, we here selected visual feature neurons by comparing feature distance between encoded vs. unselected identities, but the selected visual feature neurons highly overlapped with the feature M-ID neurons as we described above), a subset of neurons that only encoded conceptual associations (*n* = 8), a subset of neurons that encoded both visual features and conceptual associations (*n* = 23), and a subset of neurons that encoded neither visual features nor conceptual associations (*n* = 17). **(i)** DNN feature distance for each subpopulation of M-ID neurons. **(j)** Conceptual association ratings for each subpopulation of M-ID neurons. S-S: pairs of identities that a neuron was selective to. S-NS: pairs of identities where a neuron was selective to one of them but not selective (NS) to the other. Error bars denote ±SEM across neurons. Asterisks indicate a significant difference using two-tailed two-sample *t*-test: *: P < 0.05, ***: P < 0.001, and ****: P < 0.0001. **(k)** Web-association score for MID neurons. For each neuron, we calculated a mean association score between the pairs of identities that the neuron was selective to (S-S), and between the pairs of identities where the neuron was selective to one of them but not selective (NS) to the other (S-NS). Error bars denote ±SEM across neurons. Left: feature M-ID neurons (*n* = 38; note that here we only included neurons encoding more than one familiar identities). Right: non-feature M-ID neurons (*n* = 54). Neither feature MID neurons nor non-feature M-ID neurons had a significantly greater web-association score for the pairs of encoded identities. n.s.: not significant. **(l)** Correlation between patients’ visual similarity ratings and DNN feature similarity (i.e., the negative of the DNN feature distance) for each DNN layer. **(m)** Correlation between DNN feature similarity and web-association scores. Pearson correlation was performed across pairs of identities (*n* = 1225). Solid circles represent a significant correlation (permutation test: P < 0.05; Bonferroni correction for multiple comparisons across DNN layers) and open circles represent a non-significant correlation. Shaded area denotes ±SD across permutation runs. **(n)** The absolute difference in social trait judgments between identity pairs. S-S: pairs of identities that a neuron was selective to. All-NS: all other pairs (i.e., all excluding S-S pairs). Error bars denote ±SEM across neurons.

### Factors that may influence feature-based identity coding

We next investigated the factors that may influence feature-based coding in identity neurons. First, previous research primarily used well-known or familiar faces to study identity neurons ^16,18,21^. It is unknown whether feature-based coding also depends on familiarity. We found that feature MID neurons encoded both familiar (i.e., patients recognized the celebrity’s face) and unfamiliar faces (only 41.3%±33.4% [mean±SD across patients] of all selected identities were familiar; feature M-ID neurons did not differentiate familiar vs. unfamiliar selected identities; **Fig. 2e**; twotailed unpaired *t*-test: *t*(52) = 0.87, P = 0.39). This data suggests that face familiarity did not play an essential role for feature-based coding in the amygdala and hippocampus (we revisit this point later with datasets consisting of all unfamiliar faces and all model synthetic faces).

Second, we found that feature M-ID neurons from the same patient encoded different parts of the feature space, covering different identities (e.g., **Fig. 1a-c** vs. **Extended Data Fig. 5a**), whereas feature M-ID neurons from different patients could encode a similar region of the feature space, covering the same identities (e.g., **Fig. 1a-c** vs. **Extended Data Fig. 5c**). Furthermore, feature M-ID neurons were distributed across areas of the MTL (20 in the amygdala and 22 in the hippocampus) and across patients.

Third, S-ID neurons and M-ID neurons had a similar spike sorting isolation distance (**Extended Data Fig. 1h**; *t*(53) = 0.99, P = 0.32), suggesting that M-ID neurons were not more likely to be multi-units consisting of several neurons.

Fourth, we found that the response of feature M-ID neurons could not be explained by cross-race, cross-gender, or cross-age effects (i.e., a concept of other races/gender/age resulted from the tendency to have more accurate recognition/representation for same-race/gender/age faces could not explain results; **Supplementary Information**).

Fifth, we found that low-level (i.e., pixel-level) features such as saliency, luminance, contrast, and wavelength could not explain feature-based identity coding (**Supplementary Information**; **Extended Data Fig. 8a-d**). We also found that pixels critical for identity recognition could not explain the response of feature M-ID neurons (**Supplementary Information; Extended Data Fig. 8e-g**). To further understand what drove the neural response of feature M-ID neurons, we sorted individual images by the amount of activity they evoked regardless of identity. We presented the top, middle, and bottom 5 images for 4 example feature M-ID neurons (**Extended Data Fig. 8h-k**). Qualitatively, we did not observe a systematic change as a function of firing rate, indicating that the response of feature M-ID neurons was driven by more abstract features.

Sixth, non-feature M-ID neurons encoded identities that were distributed in the feature space, but this was not likely due to our feature spaces: they did not turn into feature neurons using other feature spaces (**Supplementary Information**; **Extended Data Fig. 9a-d**). We also confirmed that the selected identities were judged as more similar compared to the unselected identities in feature M-ID neurons, and that the identities encoded by the feature M-ID neurons were judged as more similar compared to those encoded by the non-feature M-ID neurons (**Supplementary Information; Extended Data Fig. 9e, f**).

Lastly, to further study the impact of face selectivity on our results and the specificity of our results to faces, we conducted two additional analyses (**Supplementary Information**). (1) We repeated our experiments with object stimuli and our results with 603 neurons from the amygdala and hippocampus demonstrated region-based feature coding in the object space (**Extended Data Fig. 10a, b**). (2) We found region-based coding when we projected simulated neural responses from an artificial neural network to our feature space (**Extended Data Fig. 10c-e**). Together, our results suggest that region-based feature coding could be a general mechanism to encode visual categories.

### Visual similarity vs. conceptual association

We conducted the following analyses to assess the relationship between visual similarity of the faces and their conceptual associations (i.e., association between identities through concepts).

First, 5 patients we recorded from rated how related each pair of identities was using a well-established paradigm to probe conceptual associations ^18,19^ (**Extended Data Fig. 11a**). They subsequently rated how visually similar each pair of identities was (**Extended Data Fig. 11b**). We found that conceptual association was highly correlated with visual similarity (Pearson’s correlation: P < 0.001 for all patients; two-tailed paired *t*-test of *r* against 0 for the group: *r* = 0.40±0.18 [mean ± SD], *t*(4) = 4.93, P = 0.008; **Fig. 2f**), suggesting that conceptual associations could be explained by a more objective measure of visual similarity (note that conceptual associations were always rated first to prevent patients from using visual similarity as a strategy to judge conceptual associations). This was also the case with patients’ familiar faces, suggesting that visual similarity could explain conceptual associations. To further confirm our results, we repeated our analyses with data from 40 control participants from the general population. Again, we found that participants’ conceptual association rating was highly correlated with the rating of visual similarity (*r* = 0.52±0.19, *t*(39) = 17.6, P < 10^-19^; **Fig. 2f**; correlation of mean rating for each face pair: *r*(1225) = 0.96, P < 10^-20^; **Fig. 2g**). However, we found separate populations of neurons encoding conceptual association and visual similarity (**Fig. 2h**). We confirmed that neurons encoding visual similarity had a smaller feature distance (**Fig. 2i**; two-tailed paired *t*-test: *t*(13) = 7.18, P = 7.15×10^-6^) and neurons encoding conceptual associations had a greater conceptual association ratings (**Fig. 2j**; *t*(7) = 2.85, P = 0.025) for the pairs of encoded/selected identities compared to the other pairs. Interestingly, we found that neurons encoding visual similarity had even lower conceptual associations for the pairs of encoded identities (**Fig. 2j**; *t*(13) = 5.43, P = 1.15×10^-4^) whereas neurons encoding conceptual associations had even larger feature distance (**Fig. 2i**; *t*(7) = 5.14, P = 0.0013). We obtained a consistent result in neurons encoding both visual similarity and conceptual associations (**Fig. 2i, j**) and in neurons encoding neither visual similarity nor conceptual associations (**Fig. 2i, j**). Furthermore, we replicated our results using the average (i.e., consensus) ratings acquired from the general controls (**Extended Data Fig. 11c-e**).

Second, we used a web-association metric ^18^ in which the names of celebrities were paired in internet searches to determine the degree to which they were associated in the search results (**Extended Data Fig. 12a**; see **Methods** for details). We restricted our analysis to the identities each patient rated as familiar, but similar results were found when we included all identities (both familiar and unfamiliar) in the analysis. We found that the web-association scores between pairs of visually similar identities were not significantly greater than between the other pairs (**Fig. 2k** left; two-tailed paired *t*-test: *t*(37) = 0.80, P = 0.43). This was also the case for non-feature M-ID neurons (**Fig. 2k** right; *t*(53) = 0.76, P = 0.45). Notably, we confirmed that the web-association scores were correlated with conceptual association ratings from both patients (**Extended Data Fig. 11f**; P < 0.05 for all patients; *t*(4) = 5.53, P = 0.0052) and general controls (**Extended Data Fig. 11f**; *t*(39) = 14.8, P < 10^-13^; **Extended Data Fig. 11g**; mean rating: *r*(1255) = 0.16, P = 4.5×10^-8^). We derived similar results using the search engine Google and the search engine Bing. Therefore, encoding of visually similar identities was not likely explained by conceptual associations measured by the web-association scores.

Third, we confirmed that visual similarity ratings from both patients and general controls were correlated with the DNN feature similarity (i.e., the negative of the DNN feature distance; **Fig. 2l** and **Extended Data Fig. 11h**), especially in the later layers. We also found that the DNN feature similarity (the negative of the full feature distance) was largely uncorrelated with the web-association scores (**Fig. 2m**; especially in the layers where we analyzed feature M-ID neurons) and the pairwise distance in the t-SNE feature space was not correlated with the web-association scores (P > 0.05 for all layers; **Extended Data Fig. 12b**), suggesting that the organization of the feature space could not be explained by conceptual associations measured by the web-association scores.

Fourth, six of our recorded patients provided social trait judgments of the stimuli using a comprehensive set of social traits ^22^ (see **Methods**). We found that the pairs of identities encoded by the feature M-ID neurons were not more similar in social trait judgments than the other pairs (**Fig. 2n**; all Ps > 0.05 except for *feminine* because the feature space was organized by gender [**Fig. 1g**], but feature M-ID neurons did not encode gender *per se* as shown above). Furthermore, we found that the absolute difference in social trait judgments between identity pairs was largely uncorrelated with visual similarity (**Extended Data Fig. 11i**) and conceptual association (**Extended Data Fig. 11j**), suggesting that our results could not be simply explained by the concept of social traits (i.e., relatedness in social traits). Notably, we confirmed that our patients’ social judgment ratings were consistent with the consensus ratings derived from a large sample (*n* = 500) of participants from the general population (**Extended Data Fig. 11k**).

Together, our results suggest that visual similarity can be dissociated from conceptual associations and our results cannot be simply accounted for by conceptual associations. We also show that both neurons encoding visual features and neurons encoding conceptual associations are present in the MTL.

### Results are robust to an independent dataset and different feature metrics

To consolidate our findings, we further analyzed data from a publicly available dataset that contains well-characterized identity neurons recorded using famous and/or familiar faces ^19^. In line with our present findings, the identity neurons from this dataset also indicated that feature-based coding is present in the MTL (**Supplementary Information; Extended Data Fig. 13** and **Extended Data Fig. 14**). In sum, we not only replicated our feature-based coding using an independent dataset with well-characterized identity neurons, but also demonstrated that feature-based coding could not simply be attributed to a cross-race or cross-gender effect. In addition, our results showed that feature-based coding was complementary to the coding of conceptual associations: we found different subsets of neurons that encoded visual facial features only, conceptual associations only, and both visual features and conceptual associations.

Lastly, similar results were derived if we constructed a three-dimensional feature space or used different perplexity parameters for t-SNE (balance between local and global aspects of the data) or kernel/cluster size parameters to detect a tuning region (balance between sensitivity and specificity of detecting a tuning region). Similar results were also derived if we constructed the feature space using other common methods, such as uniform manifold approximation and projection (UMAP; **Extended Data Fig. 15**) or principal component analysis (PCA). We could also replicate our findings using full DNN features, where the Euclidian distance between encoded identities was significantly smaller than that of non-encoded identities (**Extended Data Fig. 2k**). This suggests that our findings were robust to the construction of the feature space.

### A broader category of feature neurons

Because different faces of the same identity were not clustered in the feature space from earlier lower-level layers of the DNN (**Extended Data Fig. 3c** and **Extended Data Fig. 4**), we next asked whether there are neurons that code for similar visual features independent of face identity. Using the same method to select identity neurons that responded to visually similar faces, we identified a broader category of *feature neurons* that were tuned to a certain region of the feature space from each DNN layer (see **Fig. 3a, b** and **Extended Data Fig. 16** for examples and **Fig. 3c** for a summary; see **Extended Data Fig. 7b, e** for a breakdown of amygdala and hippocampal neurons), regardless whether the neuron was an identity neuron.

**Fig. 3.**
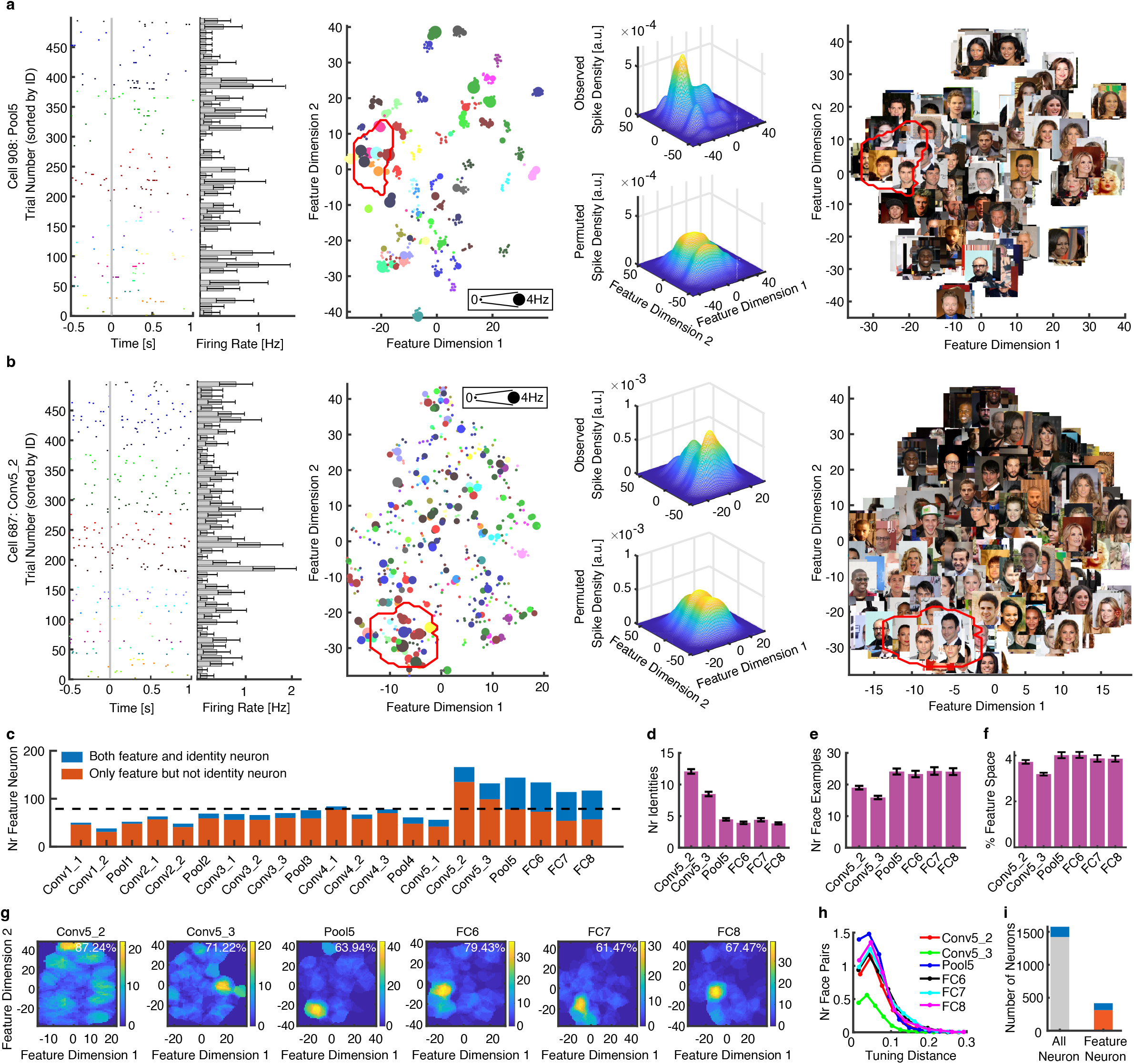
Characterization of feature neurons. **(a, b)** Two example feature neurons that encoded visually similar faces. Legend conventions as in **Fig. 1**. **(c)** The number of feature neurons identified from each DNN layer. Blue: feature neurons that were also identity neurons. **(d)** The number of identities encoded by feature neurons. **(e)** The number of faces encoded by feature neurons (i.e., the number of faces that fell within the tuning region of a feature neuron). Error bars denote ±SEM across neurons. **(f-h)** Population summary of feature tuning. Legend conventions as in **Fig. 2**. **(i)** The number of identity neurons in the whole population (left) and among feature neurons (right). Blue: the number of identity neurons (*n* = 155 for whole population and *n* = 104 for feature neurons). Red: the number of non-identity feature neurons (*n* = 312). Gray: the number of non-identity neurons (*n* = 1422).

First, we found that feature neurons mostly appeared in later DNN layers where faces started to become clustered by identities. Therefore, feature neurons from the MTL primarily encode higher-level visual information related to identification rather than lower-level image characteristics. Six of the later DNN layers (Conv5_2, Conv5_3, Pool5, FC6, FC7, and FC8) contained an above-chance number of feature neurons at the population level and we restricted our analysis to these feature neurons. The number of identities (**Fig. 3d**) and faces (**Fig. 3e**) covered by the tuning region of feature neurons indicated the size of the “receptive field” (in feature space) of these feature neurons. The tuning region of each feature neuron covered approximately 2.5-11% of the feature space (**Fig. 3f**) and the total observed neuronal population covered approximately 61-87% of the feature space (**Fig. 3g**). With increasing levels of abstraction, tuning regions in later layers Pool5, FC6, FC7, and FC8 contained fewer identities (**Fig. 3d**; two-tailed unpaired *t*-test: P < 0.001 for all comparisons) but more faces (**Fig. 3e**; P < 0.001) compared to the preceding convolutional layers. Faces were also more widely distributed in the FC layers than the preceding layers (**Fig. 3h**; two-tailed unpaired *t*-test on feature distance: P < 0.0001).

Second, although an appreciable proportion of feature neurons were identity neurons, some feature neurons were not identity neurons (i.e., neither S-ID nor M-ID neurons; red bars in **Fig. 3c**; in particular in convolutional layers; see **Fig. 3a** for an example) because they covered a region in the face space containing faces from different identities. Therefore, identity selectivity was not necessary for feature-based coding. In other words, feature neurons can respond to faces that were adjacent in the feature space but were not from the same identity.

Third, we investigated whether feature neurons were more likely to be identity neurons (i.e., either S-ID or M-ID neurons). Indeed, we found that feature neurons had a higher proportion (104/417; 24.94%) of identity neurons compared to the entire neuron population (155/1577; 9.83%; χ^2^-test: P = 3.33×10^-16^; **Fig. 3i**; note that here feature neurons included those from layer Conv5_2 and Conv5_3 even though identity neurons could in principle only emerge in layers with clustering of faces; see **Extended Data Fig. 7c, f** for a breakdown of amygdala and hippocampal neurons), suggesting that region-based feature tuning is a key component in identity selectivity.

Lastly, we tested the specificity of the feature space and stimuli in identifying feature neurons (**Supplementary Information**). Although to some extent region-based feature coding could still be observed in feature spaces constructed using DNNs trained for object recognition (e.g., AlexNet [**Extended Data Fig. 17a, b**] and ResNet [**Extended Data Fig. 17e-h**]), there were fewer feature neurons (**Supplementary Information**), suggesting that MTL neurons were sensitive to the organization of the feature space and thus the organization of the feature space played a critical role in identifying feature neurons. In addition, by projecting non-face stimuli onto the AlexNet (**Extended Data Fig. 17c, d**), ResNet (**Extended Data Fig. 17i-l**), or the original VGG-Face (**Extended Data Fig. 17m-p**) feature spaces, we found that faces had unique visual features and feature neurons encoded higher-level visual features related to faces rather than lower-level visual features that might be in common between face and non-face stimuli.

### Validation of region-based feature coding by two additional experiments

We conducted two additional experiments to validate region-based feature tuning using different stimuli (especially out-of-set stimuli for generalization) and explored whether such feature coding could be generalized to unfamiliar faces (note that patients should have little conceptual knowledge about these unfamiliar faces, in particular the FaceGen model faces). In the first additional experiment, we recorded from 837 neurons in the same 10 patients (27 sessions; firing rate > 0.15 Hz; accuracy = 75.7%±23.0% [mean±SD across sessions]) using face stimuli from the FBI Twins dataset (**Fig. 4a**), which were all novel to our patients. We applied the same DNN to extract face features and construct face feature spaces. We again found region-based feature coding by single neurons in this experiment (see **Fig. 4a** and **Extended Data Fig. 18a, b** for examples and **Fig. 4b-e** for group results). This suggests that feature coding by neurons in the MTL did not depend on the faces being familiar to the participants. Feature coding was evident even if we restricted our analyses to the very first exposure of the faces, when they were novel.

**Fig. 4.**
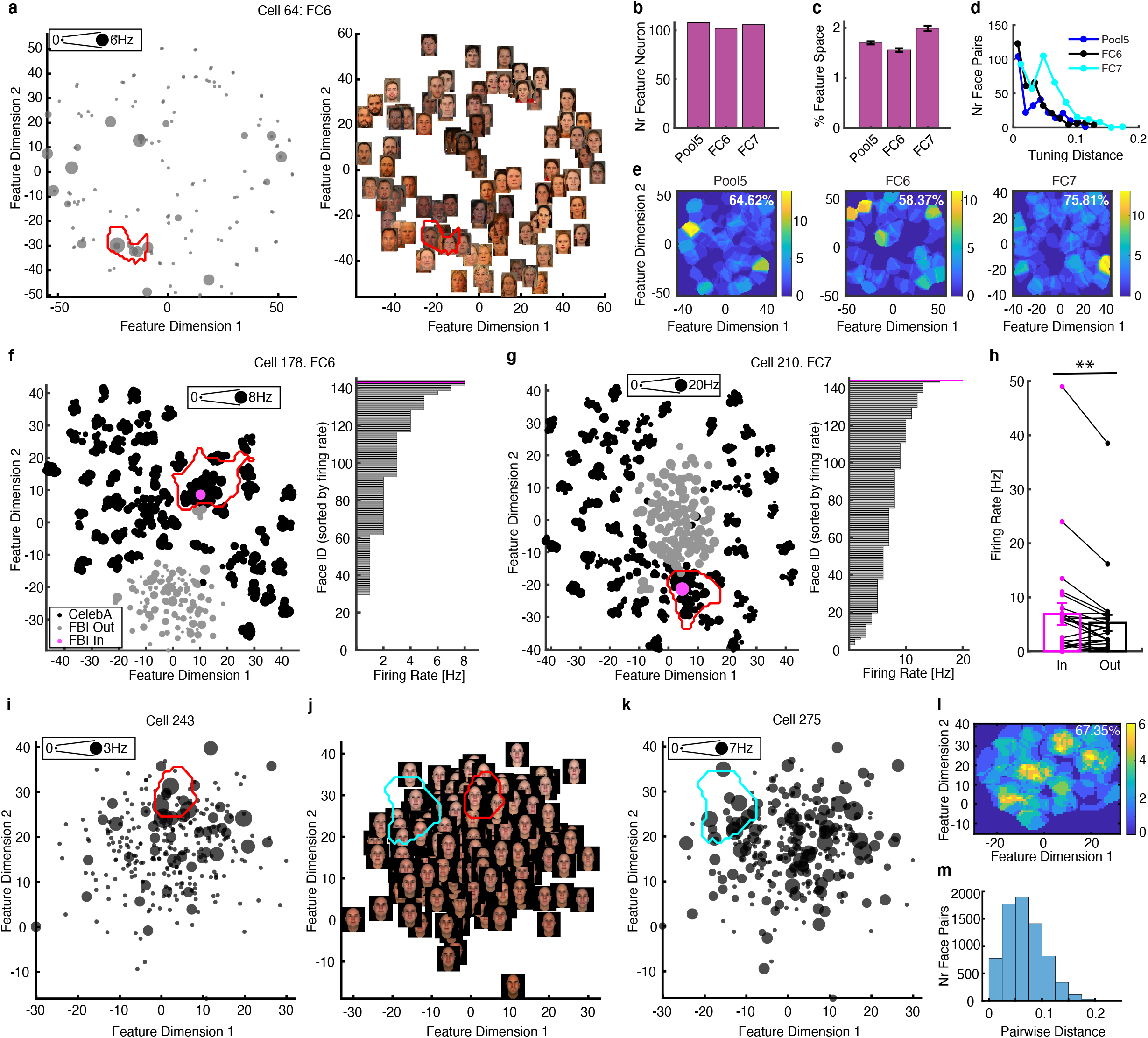
Validation and generalization of feature tuning with unfamiliar and model faces. **(a-h)** Results from the FBI Twins dataset. **(i-m)** Results from the FaceGen dataset. **(a)** An example neuron demonstrating region-based feature coding. In FBI face spaces, similar faces were also clustered and faces from different genders were organized in different areas of the feature space. The size of the dot indicates the firing rate. The red outline delineates the tuning region of the neuron in the feature space. **(b)** The number of identified feature neurons using FBI stimuli. Only DNN layers with an above-chance number of feature neurons (based on our simulations) are shown. **(c-e)** Population summary of feature tuning. Legend conventions as in **Fig. 2**. **(f, g)** Example CelebA feature neurons showing elevated responses for FBI stimuli falling in their tuning regions. Feature spaces were constructed for combined CelebA and FBI stimuli. The size of the dot indicates the firing rate. The red outline delineates the tuning region of the neuron (identified by the CelebA stimuli). Black: faces from the CelebA stimuli. Gray: faces from the FBI stimuli. Magenta: FBI stimuli falling in the tuning region of the neuron. Note that in the face feature space combining the CelebA and FBI stimuli, the clustering based on gender and skin color was retained and similar to the feature spaces using the CelebA stimuli only. **(h)** Population results comparing neuronal response to FBI stimuli falling in vs. out of the tuning region (*n* = 26). Each dot represents a neuron. Error bars denote ±SEM across neurons. Asterisks indicate a significant difference between In vs. Out responses using one-tailed paired *t*-test (P < 0.01). **(i-k)** Two example neurons demonstrating region-based feature coding. Note that the feature space was constructed using parameters (i.e., features) used to synthesize the faces rather than DNN features. The dimensions of the feature space are the first shape and tone/texture principal components (PCs) used to generate the stimuli. Note that face shape varied along Feature Dimension 1 and skin color varied along Feature Dimension 2. **(l, m)** Population summary of feature tuning (*n* = 61). Legend conventions as in **Fig. 2**.

Notably, we also recorded the response of a subset of the same 699 neurons using the CelebA stimuli and we were thus able to directly investigate the generalizability of feature tuning between these two tasks. In the common feature space for the CelebA and FBI stimuli, the tuning region of 38 CelebA feature neurons overlapped with identities from the FBI stimuli (**Fig. 4f, g** and **Extended Data Fig. 18c-e**). We found that FBI stimuli in the CelebA feature neurons’ tuning regions elicited a significantly greater response compared to the other FBI stimuli that were not inside the CelebA feature neurons’ tuning regions (paired *t*-test: *t*(25) = 2.61, P = 0.00076; see **Fig. 4f, g** for examples and **Fig. 4h** for group results). This shows that region-based feature tuning generalized between different image sets as well as to novel stimuli never seen by the participant before. Because the faces from the two datasets were in different styles, the distributions of the two datasets were separated to some extent. We repeated our analysis while restricting our comparison to images of the FBI dataset that showed overlap with the CelebA stimuli in the feature space (we applied a mask that defined the vicinity of the CelebA stimuli), so that the “Out” condition only contained faces that were inside the subregion occupied by the CelebA stimuli (**Extended Data Fig. 18f-h**). We confirmed our results and still found a greater response for FBI stimuli that were in the CelebA feature neurons’ tuning regions (**Extended Data Fig. 18i**; one-tailed paired *t*-test: *t*(26) = 2.69, P = 0.0062).

In the second additional experiment, we recorded from a separate population of 658 neurons (25 sessions from 6 patients; firing rate > 0.15 Hz) while patients performed a trustworthiness or dominance judgment task using the FaceGen model faces (**Fig. 4i-m**) ^23^, which contained only feature information but no real identity information. Behaviorally, the ratings from our patients were consistent with the consensus ratings ^23^ (Pearson correlation: *r* = 0.23±0.14 [mean±SD across sessions] for trustworthiness and *r* = 0.39±0.18 for dominance; two-tailed *t*-test against 0: both Ps < 0.001). Again, we found region-based feature coding (61 neurons; above-chance compared to our simulations; each neuron covered 2.50%±0.66% [mean±SD] of the feature space; see **Fig. 4i-k** for examples and **Fig. 4l, m** for group results). It is worth noting that FaceGen model faces did not convey information about concepts of the face identities, therefore, region-based feature coding could not be explained by conceptual associations between face identities.

Together, these additional experiments showed that region-based feature tuning could be generalized to new and unfamiliar face stimuli and was independent of conceptual associations.

### Axis-based vs. region-based feature coding

The region-based feature coding we found is different from the face feature coding shown in the IT cortex of non-human primates where neurons parametrically correlate with facial features along specific axes in face space (which corresponds to a linear combination of DNN features) ^7-10,24^. Rather than such axis code, MTL neurons encoded a certain range of feature values in the face space. We previously found that some human amygdala neurons encode a linear change in facial emotions ^25^. Therefore, we wondered whether some MTL neurons encode a linear combination of facial features as shown in the primate IT cortex ^7,9^. To answer this question, we used established approaches (see **Methods**) and identified a small subset of neurons that encoded a linear combination of DNN features (**Fig. 5a** and **Extended Data Fig. 19c, d**; similar results were derived using other models; **Extended Data Fig. 19a**), although this response was primarily driven by region-based feature coding because an elevated response in one part of the feature space could drive the regression (see **Extended Data Fig. 19e, f** for examples; see **Fig. 5c-f** for a comparison). Therefore, feature-based coding in the human MTL is primarily region-based rather than axisbased.

**Fig. 5.**
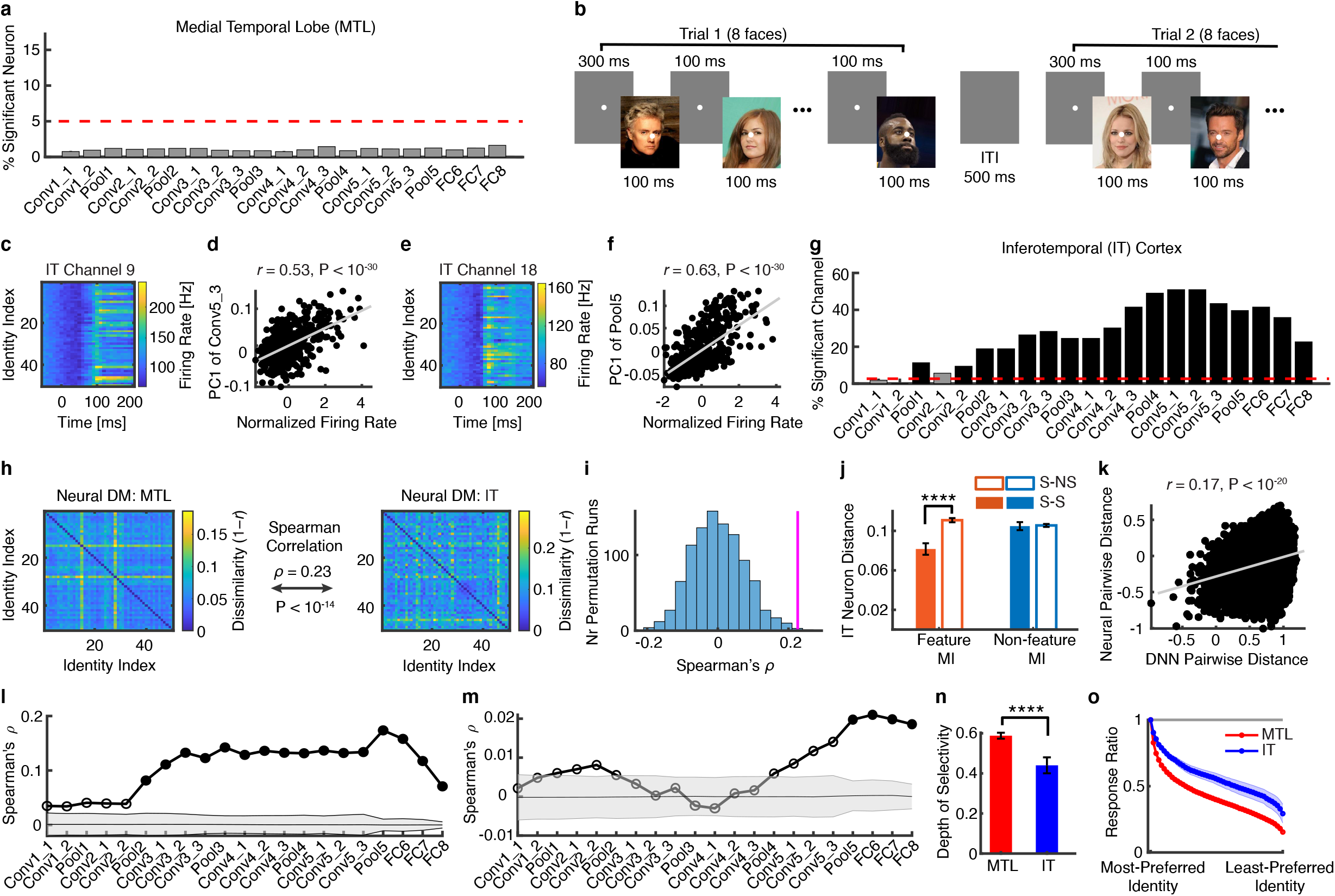
Comparison of coding in the inferotemporal (IT) cortex and medial temporal lobe (MTL). **(a)** A small proportion of MTL neurons demonstrated axis-based coding of facial features. Selection of neurons was performed using partial least squares (PLS) regression with deep neural network (DNN) feature maps. Dashed line denotes the chance level. **(b)** Task used to acquire neural responses from a monkey. In each trial, 8 faces (CelebA stimuli) were presented for 100 ms each, followed by a fixed inter-stimulus-interval (ISI) of 100 ms. There was a central fixation point of 300 ms at the beginning of each trial and there was an inter-trial-interval (ITI) of at least 500 ms following each trial. The central fixation point persisted through the trial. **(c-f)** Two example monkey IT multi-unit activity (MUA) channels showing axis-based feature coding. **(c, e)** MUA to 50 identities, shown in 10 ms time bins. Time 0 denotes the stimulus onset. Firing rate was normalized to the average of the gray images (i.e., control stimuli). **(d, f)** Correlation between the firing rate and the first principal component (PC1) of the feature map. Each dot represents a face pair, and the gray line denotes the linear fit. Note that both channels had a significant relationship with the feature map (PLS regression, permutation P < 0.001), and we show the correlation with PC1 for illustration purposes. **(g)** The proportion of IT MUA channels demonstrating axis-based feature coding. Selection of channels was performed using PLS regression with DNN feature maps. **(h)** Correlation between dissimilarity matrices (DMs). The MTL DM (left matrix) was correlated with the IT DM (right matrix). Color coding shows dissimilarity values (1-*r*). **(i)** Observed vs. permuted correlation coefficient between DMs. The correspondence between DMs was assessed using permutation tests with 1000 runs. The magenta line indicates the observed correlation coefficient between DMs. The null distribution of correlation coefficients (shown in gray histogram) was calculated by permutation tests of shuffling the face identities. **(j)** Neural distance (1-Pearson’s *r*) calculated using all IT channels. For each MTL neuron, we calculated the IT neural distance between the pairs of stimuli that the MTL neuron was selective to (S-S; solid bars), and between the pairs of stimuli where the MTL neuron was selective to one of them but not selective (NS) to the other (S-NS; open bars). Error bars denote ±SEM across neurons. Left: feature M-ID neurons (*n* = 42; red). Right: non-feature M-ID neurons (*n* = 60; blue). Only feature M-ID neurons but not non-feature M-ID neurons had a shorter IT neural distance. Asterisks indicate a significant difference using two-tailed paired *t*-test. ****: P < 0.0001. **(k)** An example IT channel showing correlation between MUA pairwise distance and DNN layer Pool4 feature pairwise distance (see **Methods**). Each dot represents a face pair, and the gray line denotes the linear fit. **(l)** Correlation between pairwise distance in the IT neuronal face space and pairwise distance in the DNN face space (see **Methods**). **(m)** Correlation between pairwise distance in the MTL neuronal face space and pairwise distance in the DNN face space. Solid circles represent a significant correlation (permutation test: P < 0.05, Bonferroni correction across layers) and open circles represent a non-significant correlation. **(n)** Depth of selectivity (DOS) index (see **Methods**). MTL identity neurons had a significantly higher DOS index than IT identity channels. For IT channels, mean response to the control stimulus (gray blank screen) was subtracted from the response to each face image. Error bars denote ±SEM across neurons/channels. Asterisks indicate a significant difference using two-tailed unpaired *t*-test. ****: P < 0.0001. **(o)** Ordered average responses from the most-to the least-preferred identity. Responses were normalized by the response to the most-preferred identity. Shaded areas denote ±SEM across neurons/channels. The top bar indicates a significant difference between MTL and IT identity channels (two-tailed unpaired *t*-test, P < 0.05, corrected by FDR for Q < 0.05). MTL neurons showed a steeper change from the most- to the least-preferred identity compared to IT channels.

Do neurons exhibit axis-based coding elsewhere in the brain? To confirm that IT neurons form an axis code for our stimuli, we conducted recordings from the IT cortex of a monkey using the same stimuli as in our main experiment (see **Methods**; 53 multi-unit activity [MUA] channels; **Fig. 5b**). First, we found that in contrast to MTL neurons (**Fig. 5a;** see **Extended Data Fig. 19g, h** for a comparison), many IT MUA channels demonstrated axis-based coding of face features (see **Fig. 5c-f** for examples and **Fig. 5g** for group summary; see **Extended Data Fig. 19b, h** for results derived using other models). Second, to assess the validity of comparing coding between the human MTL and macaque IT cortex, we used representational similarity analysis (RSA) ^26^ and found that the neuronal populations in the MTL and IT cortex had a similar representational structure (**Fig. 5h, i**; permutation P = 0.001). Furthermore, the identities encoded by MTL feature M-ID neurons also had a more similar neural encoding by IT neurons relative to the pairs not encoded by the same feature M-ID neuron (**Fig. 5j**; feature M-ID neurons: paired two-tailed *t*-test: *t*(41) = 4.61, P = 3.86×10^-5^; non-feature M-ID neurons: *t*(59) = 0.22, P = 0.82). Third, using a pairwise distance metric (see **Methods**) ^27^, we found that IT neurons primarily corresponded to the intermediate DNN layers (**Fig. 5k, l**), whereas MTL neurons primarily corresponded to the top/output DNN layers (**Fig. 5m**), consistent with their different processing stages along the ventral processing pathway. Lastly, we found that identity selectivity was significantly stronger in MTL neurons compared to IT neurons (**Fig. 5n, o**; DOS: two-tailed unpaired *t*-test: *t*(201) = 12.8, P = 7.70×10^-28^), supporting the abstraction towards coding specific people in the MTL. As a control, we derived similar results when we used only the first presentation of the stimuli in the monkey IT data to align with the single presentation in human recordings (**Extended Data Fig. 20a-e**). We also derived similar results using MUA rather than single-unit activity in the human MTL data to be more similar to the monkey recordings (**Extended Data Fig. 20f-m**).

Together, we show that in contrast to MTL neurons, IT neurons encoded the axes of the feature space and thus demonstrated axis-based coding for our stimulus set. Our results are in line with the flow of information along the ventral visual processing stream.

## Discussion

Our results reveal that the response of identity neurons in the human amygdala and hippocampus can encode identities that are related visually (i.e., faces sharing similar features) rather than conceptually (e.g., Bill and Hillary Clinton). We further show a broader category of feature neurons exhibiting region-based feature coding, a novel type of cell in the human MTL. The response of feature neurons does not depend on identity selectivity nor face familiarity, gender, race, or low-level image features, and their tuning regions can be validated using new face stimuli. Lastly, we show that in contrast to the macaque IT cortex, feature-based coding in the human MTL is primarily region-based rather than axis-based.

The amygdala and hippocampus are downstream of the face-selective regions in the higher visual cortex, where feature-based coding for faces is first evident ^11–13^. Despite being downstream from face-selective areas, in the human MTL no feature-based encoding of faces has been found. Instead, only exemplar-based coding has been demonstrated so far ^16,17^. This indicates that the format of the neural representation is fundamentally differently in the MTL compared to the higher visual cortex. It remains unknown how this transformation from a perception-driven to the memory-based semantic representation in the MTL is achieved. The novel type of cells we describe here encode an intermediate type of representation between these two formats. These previously unknown neurons encode a region in the high-level feature space such that they become selective to all the identities that fall into this region, thereby providing a bridge between these two coding mechanisms. We hypothesize that this form of representation serves as a basis for semantic representations in the MTL, which in turn are the basis for declarative memory ^28^. Therefore, our findings bridge the two extreme hypotheses by revealing an intermediate region-based feature code in the MTL.

To contrast our findings with prior work in macaques, we have established that for our stimuli macaque IT neurons exhibit an axis-based code as expected ^7-10,24^ (note that feature-based coding at the single cell level has only been shown in the macaque IT cortex, so we did this work with macaques to link to this literature). While macaque IT and human MTL representations were similar as assessed by representational similarity, responses in these two brain areas were best explained by different processing stages in our DNN and identity coding was significantly more prominent in the MTL, supporting our conclusion that region-based encoding is a feature of the human MTL but not macaque IT cortex (it remains an open question whether region-based encoding exists in the human IT cortex).

Although it may not be possible to entirely discount any sort of conceptual association in explaining our results, our various control analyses have suggested that visual similarity is dissociable from conceptual association and our results could not be simply attributed to conceptual associations of identities. First, feature neurons encode unfamiliar identities (shown by both CelebA and FBI stimuli; all identities from the FBI stimuli were unfamiliar to the patients), for which patients cannot have formed much conceptual knowledge; and feature coding could not be explained by simple conceptual knowledge of race, gender, or age (i.e., cross-race, cross-gender, or cross-age effects could not explain our results), nor social trait judgments. Feature-based coding of identities was even observed with synthetic model faces, of which patients had little conceptual knowledge. Second, when patients had a considerable knowledge of the identities (**Extended Data Fig. 13** and **Extended Data Fig. 14**), we observed a double dissociation of coding visually similar identities and coding of conceptually associated identities, consistent with our results using the CelebA stimuli (**Fig. 2**). Furthermore, conceptual association measured using web-association scores could neither explain feature-based coding or correlate with DNN features. On the other hand, the novel mechanism of coding visual features in the human MTL is not mutually exclusive but may complement the well-known coding of conceptual associations ^18,19^ (given the high correlation between conceptual association and visual similarity ratings [**Fig. 2f, g**], many prior findings in conceptual associations may in fact be explained by the objective measure of visual similarity). Indeed, we found separate subsets of neurons that only encoded visual features or conceptual associations, but also neurons that encoded both (**Fig. 2h**); and we found feature-based coding when we analyzed an independent dataset with well-characterized identity neurons ^19^ (**Extended Data Fig. 13** and **Extended Data Fig. 14**), an under-appreciated role of MTL neurons in these prior studies ^18,19^. Therefore, MTL neurons embody two forms of coding of face identities that may well complement each other. Notably, concept neurons and encoding of conceptual associations have also been found to be prominent in non-face stimuli ^17-19^; and thus a future study will be needed to investigate feature-based coding in a broader object space (e.g., ^10^). Our data has found compelling evidence that region-based feature-coding can be extended to non-face object stimuli (**Extended Data Fig. 10**).

Neurons in the human MTL have been shown to demonstrate prominent categorical responses to visual objects (i.e., visual selectivity) ^29^ and facial expressions of emotions (i.e., emotion selectivity) ^30,31^. Region-based feature coding may also provide an account for visual and emotion selectivity: objects or emotions falling within the coding region of a neuron may elicit an elevated response. A future direction will be to construct the feature space for objects in general (e.g., using the convolutional neural network AlexNet) and investigate region-based feature coding in this feature space. A face feature space in the MTL also supports the hypothesis that cognitive maps in the hippocampus ^32^ may generalize to all relevant dimensions in life experiences ^33^.

Previous research on identity neurons primarily used familiar faces ^16,18,21^. In the present study, we found that region-based feature coding of face identity was independent of face familiarity, similar to feature coding by primate IT cortex neurons that even encode computer generated faces ^7,9^. It is also worth noting that in contrast to the traditional axis-based face spaces where axes of the space and coordinates of faces are fixed ^7,34^, the feature space constructed by t-SNE in the present study varies as a function of the set of input stimuli because it models the similarity between all input stimuli (but note that our results were robust to the construction of the feature space and could be replicated using Euclidian distance of full DNN features). Therefore, our observed feature neurons in the human MTL may demonstrate a form of similarity-based or manifold-based coding (i.e., finding meaningful low-dimensional structures hidden in the high-dimensional observations using nonlinear dimensionality reduction) ^35,36^, which may in turn contribute to the MTL’s critical role in face recognition, classification, and memory.

Rapid advances in computer vision and development of DNNs have provided an unprecedented opportunity to help us understand the functional architecture of the brain ^8,24,27,37^. Our present study reiterates the advantages of using DNNs to study neural encoding for face identity: by extracting facial features from complex natural face images using DNNs and projecting them onto the feature space constructed by DNN feature reduction, we revealed a novel face code in the human MTL whereby neurons encode visually similar identities.

## Acknowledgements

We thank all patients for their participation, staff from WVU Ruby Memorial Hospital for support with patient testing, Minglei Yin and Stefan Uddenberg for help on analysis, Jeremy Dawson for contributing the FBI Twins dataset, and Ralph Adolphs, Carlos Ponce, Le Chang, Doris Tsao, Liang She, and Paula Webster for discussion and valuable comments. This research was supported by Air Force Young Investigator Program Award (FA9550-21-1-0088), NSF Grants (BCS-1945230, IIS-2114644), and Dana Foundation Clinical Neuroscience Award. The funders had no role in study design, data collection and analysis, decision to publish, or preparation of the manuscript.

## Author Contributions

R.C., A.T., X.L., and S.W. designed research. R.C., A.P., and S.W. performed experiments. N.J.B. performed surgery. R.C., J.W., C.L., E.D.F., A.P., H.G.R., X.L. and S.W. analyzed data. R.C, J.D., A.T., U.R., X.L., and S.W. wrote the paper. All authors discussed the results and contributed toward the manuscript.

