## Supplementary Materials for "Feature-based encoding of face identity by single neurons in the human amygdala and hippocampus"

#### Supplementary Information

#### Supplementary Results

*Identity coding with familiar vs. unfamiliar faces*

We analyzed identity selectivity separately for familiar faces and unfamiliar faces. Using the same selection criteria but given reduced statistical power, we identified 102 identity-selective neurons with familiar faces (6.47%; binomial $P = 0.0004$; 83 S-ID neurons and 19 M-ID neurons) and 85 identity-selective neurons with unfamiliar faces (5.39%; binomial $P = 0.22$; 43 S-ID neurons and 42 M-ID neurons). Notably, neurons selective to familiar identities demonstrated a significantly better decoding performance than neurons selective to unfamiliar identities (**Extended Data Fig. 2j**), consistent with previous results showing that MTL neurons respond more strongly to familiar faces than unfamiliar faces [1].

*Control for cross-race, cross-gender, or cross-age effects*

Could the response of feature M-ID neurons be attributed to cross-race, cross-gender, or cross-age effects? We found that feature M-ID neurons encoded both Caucasian faces and African-American faces (**Fig. 1** and **Extended Data Fig. 5**). All our patients are Caucasian and we found that feature M-ID neurons encoded both identities that were of the same race as the patient (e.g., **Fig. 1d-f**) as well as identities that were of a different race (e.g., **Fig. 1a-c)**. It is worth noting that for each race, feature M-ID neurons only encoded a subset of identities of that race (**Fig. 1** and **Extended Data Fig. 5**), suggesting that feature M-ID neurons did not simply encode the concept of one race or the concept of the same race or a different race. Additionally, because feature M-ID neurons encoded faces that were familiar to the patient as well as faces that were unfamiliar, the responses of feature M-ID neurons were not likely due to face unfamiliarity (i.e., the possibility that patients could not distinguish the identities or faces and thus grouped all unfamiliar faces as one category; for example, the patient was familiar with and able to recognize both identities shown in **Fig. 1a-c**). Furthermore, feature M-ID neurons encoded both male (e.g., **Fig. 1a-c**) and female (e.g., **Fig. 1d-f**) identities and they encoded identities that were of the same gender as well as the opposite gender as the patients (see also **Extended Data Fig. 5**). Lastly, by grouping the identities in the stimuli and patients by age group (e.g., young, middle-aged, elderly), we found that feature M-ID neurons encoded identities that were of a similar age as the patient as well as identities that were of a different age. Furthermore, using patients' own judgments of youthfulness of faces (see **Methods**), we found that the encoded identities did not differ significantly in youthfulness compared to the non-encoded identities (two-tailed paired $t$-test: $t(35) = 1.56$, P = 0.13). Taken together, this data suggests that the response of feature M-ID neurons could not be explained by cross-race, cross-gender, or cross-age effects.

*Control analysis for low-level visual features*

Could the response of feature M-ID neurons be attributed to low-level visual features such as saliency [2], luminance (i.e., luminance after RGB values converted to the CIE 1976 $L^*$, $u^*$, $v^*$ color space), contrast (RMS contrast, i.e., the standard deviation of the pixel intensities), and wavelength (mean hue value after RGB converted to hue, saturation, and value [HSV]; see https://osf.io/auzjy/ for detailed definition and implementations)? We extracted these low-level visual features and found that the selected identities (i.e., identities within the tuning region) did not differ in these low-level visual features compared to unselected identities (i.e., identities outside the tuning region; **Extended Data Fig. 8a-d**; two-tailed two-sample $t$-test: all Ps > 0.05). Furthermore, to reveal the critical pixels from the stimuli that contributed to identity recognition and provide an explanation of our model's output in the domain of its input, we applied layer-wise relevance propagation (LRP) to our DNN. LRP can use the network weights created by the forward-pass to propagate the output back through the network up until the original input image. The explanation given by LRP is a heat map of which pixels in the original image contribute to the final output (red pixels *positively* contributed to the classification whereas blue pixels *negatively* contributed to the classification). By comparing the LRP heat maps between selected (**Extended Data Fig. 8e**) vs. non-selected (**Extended Data Fig. 8f**) identities, we found that the selected identities used similar pixels in the input images for identity recognition compared to unselected identities (**Extended Data Fig. 8g**), suggesting that the response of feature M-ID neurons could not be explained by critical pixels for identity recognition.

*Control analysis for non-feature M-ID neurons*

Non-feature M-ID neurons encoded identities that were distributed in the feature space. Was this due to the feature space that we used? To answer this question, we constructed other feature spaces using the AlexNet [3] and ResNet [4]. We found that identities encoded by the non-feature M-ID neurons still did not cluster in these feature spaces (see **Extended Data Fig. 9a, b** for examples) and the percentage of non-feature M-ID neurons that became feature M-ID neurons in these feature spaces was at chance (**Extended Data Fig. 9c, d**; $\chi^2$-test against the overall percentage of feature M-ID neurons: all Ps > 0.05). In addition, we acquired visual similarity ratings between face identities from general controls (**Fig. 2**) and employed a Siamese neural network (i.e., a convolutional neural network that assigns a similarity score between input images) [5] to estimate the visual similarity between face identities. We confirmed that identities encoded by non-feature M-ID neurons were less visually similar than those encoded by feature M-ID neurons (**Extended Data Fig. 9e, f**; two-tailed two-sample $t$-test: human rating: $t(100) = 3.97$, P = 0.00013; Siamese score: $t(100) = 1.96$, P = 0.052). We also confirmed that the selected identities were rated as more similar compared to the unselected identities in feature M-ID neurons (**Extended Data Fig. 9e, f**; two-tailed paired $t$-test: human rating: $t(41) = 4.97$, P = 0.000012; Siamese score: $t(41) = 4.42$, P = 0.00007) but not non-feature M-ID neurons (**Extended Data Fig. 9e, f**; human rating: $t(59) = 1.92$, P = 0.06; Siamese score: $t(59) = 0.39$, P = 0.69).

*The role of face selectivity in region-based feature coding*

We conducted the following analyses to explore the role of face selectivity in region-based feature coding.

First, we used 500 images from the ImageNet with features extracted using the ResNet. The 500 images came from the following 50 categories: arachnid, bark, battery, beverage, board, bread, brier, building, car, cat, collection, crustacean, dainty, dog, electrical device, electronic device, equipment, fare, fern, fish, flower, frog, fruit, fungus, furniture, game bird, gymnast, herb, hole, insect, light, man clothing, moped, musical instrument, needlework, nest, plate, reptile, ridge, rock, rodent, star, sugar maple, support, tool, utensil, vegetable, vessel, weapon, young mammal. The ResNet could recognize the object category with an accuracy of 82.6%. We recorded 603 neurons from the MTL (311 neurons from the amygdala, 138 neurons from the anterior hippocampus, and 154 neurons from the posterior hippocampus) from 5 patients (12 sessions in total). Patients performed the same one-back task (accuracy = 69.83%±25.43%; mean±SD across sessions). Using the same selection procedure of feature neurons, we found that 48 neurons demonstrated region-based feature coding (7.96%; binomial P = 0.0007; **Extended Data Fig. 10a, b**). Therefore, region-based feature coding was not specific to faces but could be generalized to non-face objects.

Second, we employed a simulation with artificial neurons from the AlexNet [3] to explore the role of face selectivity in region-based feature coding. We input our CelebA stimuli and the same 500 objects as above to the AlexNet and acquired activation of each AlexNet unit from layer FC6 (after ReLu; $n$ = 4096 units in total, but note that only units that had an average response greater than 0 before ReLu were included for further analysis, leading to a total of $n$ = 3650 units). We then contrasted the activation of CelebA faces to the activation of ImageNet objects to select feaze-selective units (right-tailed two-sample $t$-test, P < 0.05, FDR corrected for multiple comparisons). We identified 929 *face-selective units* (**Extended Data Fig. 10c, d**) that had a significantly higher response to faces than objects, and the remaining 2721 units were defined as *non-face-selective units* (**Extended Data Fig. 10g, h**). We subsequently input the response of AlexNet face-selective units and AlexNet non-face-selective units to our original feature space (constructed by the VGG-Face layer FC6) and employed the same selection procedure of feature neurons. We found that both face-selective units (87.8%; **Extended Data Fig. 10e, f**) and non-face-selective units (31.6%; **Extended Data Fig. 10i, j**) from the AlexNet demonstrated region-based feature coding, although face-selective units had a significantly higher proportion of neurons demonstrating region-based feature coding ($\chi^2$-test: P < $10^{-20}$). It is worth noting that the AlexNet was pre-trained for object recognition whereas the VGG-Face was pre-trained for identity recognition. Therefore, the response of AlexNet units to CelebA stimuli may not reflect the same face processing as in the human brain.

Together, our results suggest that region-based feature coding could be a general mechanism to encode visual categories and it may not be specific to faces.

*Replication of feature-based coding using an independent dataset*

We further analyzed feature-based coding using data from a publicly available dataset (https://doi.org/10.25392/leicester.data.8796335.v1) [6]. There was a total of 37 neurons responding to more than one identity (i.e., multiple identity [M-ID] neurons), based on the responses to pictures from famous faces, family members, non-face objects, animals, and landmarks. We conducted two analyses. First, by restricting our analysis to identities that could be associated to individual faces (removing, for example, landmarks, animals, and music bands) while removing the concepts for which we could not compute an association score, we had a subset of 16 neurons from 5 patients, and we focused on this subpopulation of M-ID neurons for further analysis. We calculated the deep neural network (DNN) features for each picture and used the coordinates in the DNN full feature space to compare feature distance between selective-selective (S-S) pictures and selective-non-selective (S-NS) pictures. We found that 8 out of 16 M-ID neurons encoded identities that were close in the layer FC6 or FC7 feature space, indicating region-based feature-based coding (see **Extended Data Fig. 13a, c** for examples and **Extended Data Fig. 13e, f** for a group summary; one-tailed one-sample Wilcoxon signed-rank test; note that to construct the t-distributed stochastic neighbor embedding [t-SNE] projection, we included the CelebA stimulus set and multiple pictures of each selected identity in order to stabilize the projection, although we did not plot the points for these additional images). We also compared the association scores between selective and non-selective concepts (S-S vs. S-NS), and found that 10 out of 16 M-ID neurons encoded identities with significantly larger association scores (**Extended Data Fig. 13a, b, g**). Notably, we found different subsets of neurons when analyzing the encoding of features and association (**Extended Data Fig. 13h**). Some of the neurons encoded conceptual associations only (e.g., Cell 1 encoded Prince William and his wife Kate Middleton; **Extended Data Fig. 13b**), the original conclusions from previous works [6,7]; some neurons encoded both conceptual association and visual similarity that we could not tease apart using this dataset (e.g., Cell 11 encoded "Rocky", "Chucky", and "The Exorcist", which were both conceptually associated and visually similar; **Extended Data Fig. 13a**); some neurons seemed to primarily encode visually similar identities that were not conceptually related (e.g., Cell 8 encoded Rambo/Sylvester Stallone and Daniel Craig; **Extended Data Fig. 13c**); and still some neurons encoded non-related identities (e.g., Cell 10 encoded Jim Carrey and John Terry; **Extended Data Fig. 13d**). Therefore, these results are in line with our feature-based coding, a novel mechanism that well complements the coding of conceptual associations (**Extended Data Fig. 13h**).

Using this well-established dataset, we further analyzed cross-race and cross-gender effects. Four out of five patients are Caucasian (the other is Black), and all 8 feature M-ID neurons encoded Caucasian identities, with one of these neurons from a Caucasian patient encoding an additional Black identity. Therefore, feature-based coding could not be explained by a cross-race effect alone. Furthermore, we found that feature M-ID neurons encoded identities that were of the same gender (12/19) as well as the opposite gender (7/19) as the patients. Three feature M-ID neurons only encoded identities of the same gender as the patient, one feature M-ID neuron only encoded identities of the opposite gender as the patient, and four feature M-ID neurons encoded both. Therefore, feature-based coding could not be explained by a cross-gender effect alone.

In the second analysis, we restricted our analysis to famous faces only and compared the encoded identities with the CelebA stimuli (all famous faces). We had a subset of 11 neurons and we focused on this subpopulation of M-ID neurons for further analysis. Based on the names of the identities, we downloaded 8 images for each identity from the internet and input these images into the feature space constructed by the CelebA stimuli (resulting in a new feature space that was slightly varied from the original one due to the addition of new stimuli, because t-SNE works by attempting to separate all categories; also note that for the same reason, adding different stimuli would result in a slightly different feature space). Again, we found that 6 out of 11 M-ID neurons encoded identities that were close in the layer FC6 or FC7 feature space, indicating region-based feature-based coding (see **Extended Data Fig. 14a, b, d** for examples and **Extended Data Fig. 14e, f** for a group summary; one-tailed one-sample Wilcoxon signed-rank test on the DNN full feature distance between S-S vs. S-NS identity pairs). Notably, some of these neurons encoded conceptual associations only (**Extended Data Fig. 14c, g**; e.g., Cell 1 encoded Prince William and his wife Kate Middleton); some neurons encoded both conceptual association and visual similarity (**Extended Data Fig. 14a, b**; e.g., Cell 2, 3, 4, 5) that we could not tease apart using this dataset (e.g., Prince Charles and Prince William are both conceptually associated and visually similar); and some neurons seemed to primarily encode visually similar identities that were not conceptually related (**Extended Data Fig. 14d**; e.g., Cell 8 encoded Rambo/Sylvester Stallone and Daniel Craig). Similar results were derived when we used Pearson correlation to calculate feature distances, and we also confirmed the cross-race and cross-gender effects. It is worth noting that in contrast to the first analysis (**Extended Data Fig. 13**), this second analysis was based on three assumptions given the sparse coding properties and the general visually invariant response of identity neurons: (1) The M-ID neurons from this dataset would not respond to our CelebA stimuli. (2) The M-ID neurons from this dataset would respond to the stimuli downloaded from the internet (note that these stimuli were not shown to the patients; in the original experiments in which the neurons were recorded, there was only one image shown to the patient but the image was repeatedly shown 30 times). (3) The M-ID neurons encoded the selected identities similarly (note that in the analysis we did not use the actual firing rate but treated each encoded identity equally). However, this second analysis further confirmed feature-based coding using a different set of pictures. Furthermore, for both analyses, we assumed that web-association scores were "universal associations" between identities [7], although we could not rule out some personal associations that were not captured by the association score.

Together, we replicated our findings with well-characterized identity neurons. We found that different subsets of neurons embodied different types of coding and the coding of visual features complemented the coding of conceptual associations very well. With this additional data, we further showed that feature-based coding could not simply be attributed to face unfamiliarity, cross-race effects, or cross-gender effects.

*Specificity of the feature space and stimuli in identifying feature neurons*

We conducted three control analyses to explore the specificity of the feature space and stimuli in identifying feature neurons.

First, we explored whether the results changed if neural responses were projected onto a different feature map. We used the AlexNet [3] (**Extended Data Fig. 17a, b**) and ResNet [4] (**Extended Data Fig. 17e-h**) that were both pre-trained for object recognition using the ImageNet stimuli [8] to construct feature spaces and projected the neural response to CelebA stimuli onto these feature spaces. Although the AlexNet and ResNet were not trained for face recognition nor fine-tuned for our stimuli, CelebA faces were still organized to some extent in these feature spaces (e.g., faces of the same gender were clustered, faces of a similar skin color were clustered, and faces of the same identity were clustered), suggesting that some processing of visual information was in common between different networks. However, the distribution of faces was not as organized as the VGG-Face feature spaces used to identify feature neurons in the present study. As a consequence, although we still observed region-based feature coding in these feature spaces, we found fewer feature neurons ($n$ = 93 for AlexNet FC6; $n$ = 46 for AlexNet FC7; $n$ = 30 for ResNet Res5b; $n$ = 97 for ResNet Res5c; $n$ = 153 for ResNet Pooling; $n$ = 60 for ResNet FC; cf. **Fig. 3c**). Notably, we observed more feature neurons in layers where the distribution of faces was more organized and resembling the VGG-Face feature spaces (e.g., ResNet Pooling; in contrast, the distribution of faces was less organized in AlexNet FC7). Therefore, our results suggested that MTL neurons were sensitive to the organization of the feature space and thus the organization of the feature space played an important role in identifying feature neurons. Interestingly, we found that using the AlexNet or ResNet, some object information was extracted from the face stimuli. For example, faces wearing hats were clustered in the ResNet Pooling layer (**Extended Data Fig. 17g**), suggesting that some faces were not organized by facial features such as skin color but object information in these feature spaces.

Second, we explored whether/what lower-level features could characterize the tuning regions of feature neurons by projecting non-face images from the ImageNet to the AlexNet (**Extended Data Fig. 17c, d**) or ResNet (**Extended Data Fig. 17i-l**) feature spaces. We found that although round objects (**Extended Data Fig. 17d**) and/or complex patterns (**Extended Data Fig. 17j**) were closer to faces in the feature spaces, face stimuli were essentially segregated from non-face stimuli (**Extended Data Fig. 17c, d, i-l**; note that as a consequence, we were not able to find non-face images in or close to the tuning regions of feature neurons), suggesting that faces had unique visual features. This also indicated that feature neurons encoded higher-level (more abstract) visual features related to the face category rather than lower-level visual features that might be in common between face and non-face stimuli.

Third, we projected non-face images from the ImageNet to the original VGG-Face feature spaces (**Extended Data Fig. 17m-p**). Again, we found that although some non-face stimuli such as animals and utensils were closer to faces, face stimuli were largely segregated from non-face stimuli (**Extended Data Fig. 17n, p**), confirming that faces had unique visual features and feature neurons encoded higher-level visual features related to faces.

#### Supplementary Discussion

*Possible caveats*

In our main experiment, we presented each image only once, which was designed to avoid differences due to differential levels of familiarity, as is common in the human MTL [9]. However, we were still able to perform a robust analysis using our DNN-based approach because our analysis was performed at the feature level and there were many repetitions of the same features across the images. Notably, feature-based coding was still evident using repeated images [6] (**Extended Data Fig. 13** and **Extended Data Fig. 14**), and our monkey data has also shown similar results using the first presentation (**Extended Data Fig. 20a-e**) vs. all presentations (**Fig. 5**) of images. In addition, it is worth noting that our DNN-based approach is able to analyze natural faces without normalizing for shape. At the same time, our FBI and FaceGen stimuli were normalized by shape and we validated region-based feature coding using these stimuli as well. Lastly, it is worth noting that different parts of the MTL are differently involved in recognition memory [10] and our present results are restricted to the amygdala and hippocampus of the MTL.

*Identity neurons*

Although the operational definition of S-ID and M-ID neurons has been well established in the prior literature [6,7] and our definition of S-ID and M-ID neurons was in line with these previous studies, given the sparse coding properties of MTL neurons [11], it has been estimated that each MTL neuron may encode up to a few dozen of the 10,000-30,000 things a person can recognize [11] (see for [12] a detailed analysis), thus primarily appearing as an M-ID neuron. It is thus worth noting that the distinction between the S-ID and M-ID neurons is operational and may depend on the stimuli used; and the distinction between the feature and non-feature M-ID neurons may depend on the feature space. Due to the limitations on the sampling of MTL neurons and on the sampling of the stimulus space, it is not feasible to know exactly how many stimuli a given MTL neuron will respond to and conversely how many MTL neurons are involved in the representation of a given identity, although our large sample size will facilitate a more accurate estimate of these numbers than has been available before. In addition, our *in silico* experiments with the same stimuli on the same DNN have also shed light on the identity coding properties [13]. Lastly, a future study with more recorded neurons will be needed to determine whether certain identities are encoded by more neurons than others. Interestingly, in our *in silico* experiments, some identities were encoded by more neurons than others [13].

We found that at the population level, the selected identities covered both dark-skinned and light-skinned faces. Therefore, our results could not be explained by the skin tone (luminance) of the face. Furthermore, we demonstrated that the response of feature M-ID neurons could not be explained by low-level / pixel-level visual features (**Extended Data Fig. 8**), consistent with our results that feature M-ID neurons encoded abstract features from the later layers of the DNN but barely encoded low-level features from the earlier layers of the DNN (**Fig. 2** and **Fig. 3**).

*Response latency*

MTL neurons show identity selectivity around ~250 ms, consistent with the large literature of single-unit studies in humans (see [14] for a review). Our results are also consistent with the response latency in the higher visual cortex [15-19]. For example, recent human single-neuron recordings in the higher visual cortex have shown that response to faces peaks around 200 ms [18,19], which is much longer than the response latency from monkeys but shorter than that from the human MTL. Furthermore, an intracranial electrocorticography (ECoG) study has shown that early fusiform face area (FFA) activity (50-75 ms) contains information regarding whether participants are viewing a face and activity between 200 and 500 ms contains expression-invariant information about face identities [16]. Therefore, it is likely that face identity signals arrive at the MTL at 250 ms. Together, the response latency in our data is consistent with the speed of semantic processing in humans.

At the same time, our data from the monkey IT cortex (**Fig. 5**) shows a latency of approximately 70 ms for processing facial features (e.g., axis-based coding), consistent with the previous studies [20,21]. It is worth noting that monkeys have a much shorter response latency than humans [22], which has been shown using the same stimuli and task [23]. It is also worth noting that human exemplar-based coding of faces as well as semantic/conceptual coding are considerably more complex than those in monkeys (humans need to keep track of and distinguish many more people; humans evolved to have larger social networks). Therefore, the relative latency differences between monkeys and humans already in the IT cortex may get further magnified in the MTL for exemplar and concept-level processing.

*Comparison with MTL lesion results*

Although MTL lesions do not cause prosopagnosia, MTL lesions have been shown to impair ability to form new memories of any kind (including for faces) [24,25], and the type of encoding we reveal in the present study is therefore likely necessary for forming new memories. The seminal study of patient H.M. shows that lesion of the MTL abolishes ability to form new memories (including for faces) as well as subsequent recognition of familiar faces (i.e., retrograde amnesia) [24,25]. Together with a plethora of literature showing the functional role of the MTL in recognition memory for faces [1,7,11,26-28], the MTL has been shown to be critically involved in face recognition, exemplar-based coding, as well as semantic representations of face identities. Based on this literature, it is thought that the MTL creates a concept cell assembly (i.e., encoding an abstract multimodal representation of a person) that is necessary to form associations between concepts and ultimately generate new memories. On the other hand, it is worth noting that the conclusions from lesion studies and the conclusions from recordings are different. Face identity representation in the MTL does not mean that the MTL therefore has to be necessary for recognition of familiar faces, because face identity representations could be used (necessarily or not) for other processing (e.g., encoding, learning, etc., as we pointed out above).

#### Online Methods

*Patients*

There were 38 sessions with 12 patients in total (**Extended Data Table 1**). All participants provided written informed consent using procedures approved by the Institutional Review Board of West Virginia University (WVU).

*Stimuli*

We used faces of celebrities from the CelebA dataset [29]. We selected 50 identities with 10 images for each identity, totaling 500 face images. To avoid the confound of cross-race or cross-gender effects, our selected stimuli included both genders (33 of the 50 identities were male) and multiple races (40 identities were Caucasian, 9 identities were African-American, and 1 identity was biracial; **Extended Data Fig. 2b**). We used the same stimuli for all patients.

All images had the same resolution, and the faces had a similar size and position in the images [29] (**Extended Data Fig. 2b**). The deep neural network (DNN; see below for details) could well recognize the faces (98.20±4.38% accuracy; mean±SD across identities); however, the DNN was not able to recognize faces based only on the background (3.84±11.12% accuracy; in contrast, the accuracy for cropped faces was 97.40±5.27%). We also confirmed that the DNN had a similar accuracy to recognize feature M-ID neurons' selected identities (97.65±4.37%) vs. unselected identities (98.48±4.42%; two-tailed two-sample *t*-test: $t(48) = 0.64$, $P = 0.53$). Furthermore, although African-American faces had a lower brightness (**Extended Data Fig. 2b**), within each race group there was no significant difference between selected identities (Caucasian: 112.46±40.88; African-American: 97.58±44.16) vs. unselected identities (Caucasian: 119.83±37.52; African-American: 101.37±39.02; Caucasian: $t(408) = 1.67$, $P = 0.10$; African-American: $t(88) = 0.35$, $P = 0.73$). Therefore, the response of feature M-ID neurons could not simply be explained by image resolution, brightness, or background. Lastly, we asked patients to indicate whether they were familiar with each identity in a follow-up survey. We confirmed with the patients that they were able to visually distinguish different identities and recognize those with which they were familiar. It is also worth noting that in the present study we focused on the coding of *identities* rather than *concepts*. Therefore, we did not expect the patients to be familiar with all identities and we did not inquire about their depth of knowledge regarding the identities, simply whether they recognized them (i.e., face familiarity).

We further used two validation datasets. First, we used a newly-collected FBI Twins Dataset that included pairs of colored photos with the following relationships: identical twins (IT), mirror twins (MT), fraternal twins (FT), mother-child (MC), father-child (FC), and spouses (SP). Therefore, this dataset contained faces with various levels of similarity, and all faces from this dataset were unfamiliar to the patients. The photographing conditions were well controlled to ensure similar background and lighting, and all photos are high resolution (3840 × 5760). There was one face per identity and a total of 144 faces.

Second, we used a FaceGen Dataset with model faces, which notably contained only feature information but no real identity information. We used the FaceGen Modeller program (http://facegen.com; version 3.1) to randomly generate 300 faces (see [30] for detailed procedures). FaceGen constructs face space models using information extracted from 3D laser scans of real faces. To create the face space model, the shape of a face was represented by the vertex positions of a polygonal model of fixed mesh topology. With the vertex positions, a principal component analysis (PCA) was used to extract the components that accounted for most of the variance in face shape. Each principal component (PC) thus represented a different holistic non-localized set of changes in all vertex positions. The first 50 shape PCs were used to construct faces that had a symmetric shape. Similarly, because skin texture is also important for face perception, 50 texture PCs based on PCA of the RGB values of the faces were also used to represent faces. The resulting 300 faces were randomly generated from the 50 shape and 50 skin texture components with the constraint that all faces were set to be Caucasian. It is worth noting that each PC is a feature dimension of the face space.

*Social trait judgment ratings of our stimuli*

To understand if social trait judgments could explain our results, we used a set of social traits that most comprehensively characterize social trait judgments [31], including warm, critical, competent, practical, feminine, strong, youthful, and charismatic. These social traits represent the four core psychological dimensions of comprehensive trait judgments of faces (warmth, competence, femininity, and youth; 2 traits per dimension), and they were well validated in the previous study [31]. Patients were asked to rate the faces on eight social traits using a 7-point Likert scale through an online rating task. We further acquired social trait ratings of the faces from 500 participants from the general population (age [M = 26.20 years, SD = 7.11], 180/500 females). Identical to the neurosurgical patients from the current study, online participants rated the faces on eight social traits using a 7-point Likert scale.

*Experimental procedure*

We used a 1-back task for CelebA and FBI stimuli. In each trial, a single face was presented at the center of the screen for a fixed duration of 1 s, with uniformly jittered inter-stimulus-interval (ISI) of 0.5-0.75 s (**Extended Data Fig. 2a**). Each image subtended a visual angle of approximately 10$^{o}$. Patients pressed a button if the present face image was *identical* to the immediately previous image. 9% of trials were one-back repetitions. Each face was shown once unless repeated in one-back trials; and we excluded responses from one-back trials to have an equal number of responses for each face. This task kept patients attending to the faces but avoided potential biases from focusing on a particular facial feature (e.g., compared to asking patients to judge a particular facial feature). The order of faces was randomized for each patient. This task procedure has been shown to be effective to study face representation in humans [17].

For FaceGen stimuli, patients performed two face judgment tasks. In each task, there was a judgment instruction, i.e., patients judged how trustworthy or how dominant a face was. We used a 1-4 scale: ‘1’: not trustworthy / dominant at all, ‘2’: somewhat trustworthy / dominant, ‘3’: trustworthy / dominant, and ‘4’: very trustworthy / dominant. Each image was presented for 1.5 s at the center of the screen. One patient performed an additional passive-viewing task. We combined data from all tasks for analysis.

Although judgment instructions might potentially impact the response of MTL neurons (but note that all responses of each neuron were under the same instruction), it provided an opportunity to investigate whether feature-based coding could be generalized to tasks with explicit judgment instructions rather than passive viewing only. Notably, we further used the consensus ratings of the stimuli from [30] to dissociate the coding of facial trustworthiness / dominance from feature-based coding of visual similarity. Although faces within a feature neuron's tuning region were perceptually similar and thus had similar facial trustworthiness / dominance, we found that other unselected faces could be similarly trustworthy / dominant in comparison to the selected faces. In other words, faces with similar facial trustworthiness / dominance were not all clustered within a feature neuron's tuning region but distributed in the feature space. Therefore, feature neurons did not encode facial trustworthiness or dominance *per se* and feature-based coding could be generalized to tasks with explicit judgment instructions.

Stimuli were presented using MATLAB with the Psychtoolbox 3 [32] (http://psychtoolbox.org) (screen resolution: 1600 × 1280).

*Feature extraction and construction of feature space*

We used the well-known deep neural network (DNN) implementation based on the VGG-16 convolutional neural network (CNN) architecture [33] to extract features for each face image (see **Extended Data Fig. 3a** for details). Fine-tuning of the FC8 layer was performed on the pre-trained VGG-Face deep model using all images of the 50 identities in the CelebA dataset (16-30 images for each identity). This was to confirm that the pre-trained model was able to discriminate the identities and ensure that the pre-trained model was suitable as a feature extractor. Specifically, we modified the output layer to 50 units for our model. Two-thirds of the stimuli were used as the training set and the remaining stimuli were used as the testing set. We used the Adam optimizer with an initial learning rate of $5\times10^{-4}$ and we had 10 epochs in total. A learning rate scheduler was applied after each epoch with the gamma value set to 0.9 to facilitate the convergence of the loss function. To update the weights during fine-tuning, we computed the cross-entropy loss on random batches of four face images (scaled to 224×224 pixels) for back propagation. We used 5-fold cross validation, which reached an accuracy of approximately 95%. Note that only the FC8 layer was fine-tuned and all the other layers were frozen. Features that differentiated identities (i.e., identity recognition) were extracted using this transferred model. The same network was also used in recent work [17] as the computational model for deep face feature extraction.

It is worth noting that our DNN was trained on both Caucasian and African-American identities and our DNN could recognize both races equally well (Caucasian: 98.05%±4.59%; African-American: 98.89%±3.33%; two-tailed two-sample *t*-test: $t(48) = 0.52$, $P = 0.61$). Importantly, we used the same parameters for all identities and the DNN had no knowledge about the race of the identities. Therefore, the organization of the feature space and clustering of identities with similar brightness or race was entirely derived from the input images and learning by the DNN.

We subsequently applied a t-distributed stochastic neighbor embedding (t-SNE) method to convert high-dimensional features into a two-dimensional feature space. t-SNE is a variation of stochastic neighbor embedding (SNE) [34], a commonly used method for multiple class high-dimensional data visualization [35]. We applied t-SNE for each layer, with the cost function parameter (Prep) of t-SNE, representing the perplexity of the conditional probability distribution induced by a Gaussian kernel, set individually for each layer. Because a sparse distribution of faces could lead to a larger tuning region, we adjusted the distribution of faces using the t-SNE perplexity parameter so that the faces were distributed approximately homogeneously (**Extended Data Fig. 4**). We implemented t-SNE in the MATLAB platform.

Notably, neither feature extraction nor construction of feature space utilized any information from neurons. Therefore, clustering of encoded identities in the feature space was not by construction.

*Electrophysiology*

We recorded using implanted depth electrodes in the amygdala and hippocampus from patients with pharmacologically intractable epilepsy. Target locations in the amygdala and hippocampus were determined by the neurosurgeon based solely on clinical need and verified using post-implantation CT. At each site, we recorded from eight 40 μm microwires inserted into a clinical electrode as described previously [9,36]. Efforts were always made to avoid passing the electrode through a sulcus, and its attendant sulcal blood vessels, and thus the location varied but was always well within the body of the targeted area. Microwires projected medially out at the end of the depth electrode and examination of the microwires after removal suggests a spread of about 20-30 degrees. The amygdala electrodes were likely sampling neurons in the mid-medial part of the amygdala and the most likely microwire location is the basomedial nucleus or possibly the deepest part of the basolateral nucleus. Bipolar wide-band recordings (0.1-9000 Hz), using one of the eight microwires as reference, were sampled at 32 kHz and stored continuously for off-line analysis with a Neuralynx system. The raw signal was filtered with a zero-phase lag 300-3000 Hz bandpass filter and spikes were sorted using a semi-automatic template matching algorithm as described previously [37]. Units were carefully isolated and recording and spike sorting quality were assessed quantitatively (**Extended Data Fig. 1**).

Consistent with our previous studies [38-42], only single units with an average firing rate of at least 0.15 Hz throughout the entire task were considered. Trials were aligned to stimulus onset. For CelebA and FBI stimuli, we used the mean firing rate in a time window 250 ms to 1250 ms after stimulus onset as the response to each face. For FaceGen stimuli, we used the mean firing rate in a time window 250 ms to 1750 ms after stimulus onset as the response to each face.

*Neural recordings from a monkey*

One male rhesus macaque (Macaca mulatta) was used in this study. All procedures conformed to local and U.S. National Institutes of Health guidelines, including the U.S. National Institutes of Health Guide for Care and Use of Laboratory Animals. All experiments were performed with the approval of the MIT Institutional Animal Care and Use Committee (IACUC).

The monkey passively viewed the original CelebA stimuli. In each trial, the monkey first viewed a white central fixation point (0.2 degrees of visual angle [DVA]) on a gray background for 300 ms to initiate a trial. Then, 8 faces were presented for 100 ms each, each followed by a blank (gray) screen for an inter-stimulus-interval (ISI) of 100 ms. The central fixation point persisted through the trial, and fluid reward was given if the monkey successfully fixated through the entire trial. The inter-trial-interval (ITI) of blank gray screen was at least 500 ms. We recorded 4155 trials in total, and we rejected 666 trials where the monkey broke the fixation (±2 DVA). For each round of presentation, we generated a random sequence for the 500 faces; and we used different sequences for different rounds of presentation. On average, each face was presented 55.7±1.49 (mean±SD) times. Note that we randomly inserted one gray image in each round of presentation as a control stimulus for baseline subtraction and normalization.

The monkey was chronically implanted with two Utah arrays (Blackrock Microsystems) in the anterior and central inferotemporal (IT) cortex (see [20,21] for details). Each array consisted of one 10-by-10 electrode grid with 96 active iridium oxide electrodes. Each electrode was 1.5 mm long with an inter-electrode distance of 400 $\mu$m. During each recording session, band-pass filtered (0.1 Hz to 7.5 kHz) neural activity was recorded continuously at a sampling rate of 20 kHz using Intan Recording Controller (Intan Technologies, LLC). We detected the multi-unit spikes after the raw data were zero-phase band-pass filtered between 300-6000 Hz (Matlab ellip function, fourth order with 0.1 decibel pass-band ripple and 40 dB stop-band attenuation), and we used multi-unit activity (MUA) for analyses. A multiunit spike event was defined as the threshold crossing when voltage (falling edge) deviated by more than three times the standard deviation of the raw voltage values. We estimated internal consistency for each channel using a standardized image set that was run before the recording session on the same day and we accepted 53 MUA channels (from two arrays) that showed sufficient internal consistency (> 0.6). Consistent with previous studies [20,21], we used the mean firing rate in a time window 70 ms to 180 ms after stimulus onset as the response to each face. We averaged the response from repeated presentations for each face.

*Selection of identity neurons*

To select identity neurons, we first used a one-way ANOVA to identify neurons with a significantly unequal response to different identities. We next imposed an additional criterion to identify the *selected identities*: the neural response of an identity was 2 standard deviations (SD) above the mean of neural responses from all identities (note that the mean neural response could be considered as a baseline for the epochs when images were shown, even if we did not subtract a baseline). These identified identities whose response stood out from the global mean were the encoded identities. We refer to the neurons that encoded a single identity as single-identity (S-ID) neurons and we refer to the neurons that encoded multiple identities as multiple-identity (M-ID) neurons.

We were able to select a similar set of neurons using the same criteria from previous studies [6,7]. Note that because identity neurons might change their firing rate for only a few stimuli, an overall response to stimulus onset might not be observed. Therefore, given such sparseness of firing of

MTL neurons, we did not impose face responsiveness (overall change of activity in response to stimulus onset compared to baseline) as a criterion for neuron selection.

*Selection of feature neurons*

To select feature neurons, we first estimated a continuous spike density map in the feature space by smoothing the discrete firing rate map using a 2D Gaussian kernel (kernel size = feature dimension range * 0.2, SD = 4). We then estimated statistical significance for each pixel by permutation testing: in each of the 1000 runs, we randomly shuffled the labels of faces. We calculated the p-value for each pixel by comparing the observed spike density value to those from the null distribution derived from permutation. We applied a mask to exclude pixels from the edges and corners of the spike density map where there were no faces because these regions were susceptible to false positives given our procedure. We lastly selected the region with significant pixels (permutation $P < 0.01$, cluster size > 2.5% of the pixels within the mask). If a neuron had a region with significant pixels, the neuron was defined as a "feature neuron" and demonstrated "region-based feature coding". We selected feature neurons for each individual DNN layer. Because the distribution of faces was more sparse for the FBI stimuli, we used a larger kernel (kernel size = feature value range * 0.3) and a lower threshold for cluster size (cluster size > 1.6% of the pixels within the mask). Note that when we constructed the FBI face space, we also included additional faces for each identity in different viewpoints (5 faces in total) to stabilize the t-SNE projection. Lastly, given the configuration of the FaceGen face space, we considered clusters whose size was greater than 1.8% of the total number of pixels of the face space. Qualitatively the same results were derived when we used different thresholds to define feature neurons.

*Identity selectivity index*

To assess each neuron's selectivity to different identities, we defined an identity selectivity index as the $d'$ between the most- and least-preferred identities:

$$IdentitySelectivityIndex = \frac{\mu_{best} - \mu_{least}}{\sqrt{\frac{1}{2}(\sigma_{best}^2 + \sigma_{least}^2)}}$$

where $\mu_{best}$ and $\mu_{worst}$ denote the mean firing rate for the most- and least-preferred identities, respectively, and $\sigma^2_{best}$ and $\sigma^2_{worst}$ denote the variance of firing rate for the most- and least-preferred identities, respectively. A similar index was used in previous studies to assess the level of selectivity to different faces [17]. It is worth noting that the identity selectivity index was not used to select identity neurons or estimate the number of neurons that were identity selective. Instead, the identity selectivity index was used to quantify the degree of identity selectivity for the identity and non-identity neurons that had already been selected.

*Response ratio*

Response ratio was calculated for each identity by first dividing by the response of the most preferred identity and then ranking the identities from the most preferred to the least preferred. The response ratio of the most preferred identity is thus 1. We compared response ratio for each ordered identity between S-ID/M-ID vs. non-identity neurons using two-tailed unpaired *t*-test (corrected for multiple comparisons using false discovery rate [FDR] [43]). A steeper change from the best to the worst identity indicates a stronger identity selectivity.

*Depth of selectivity (DOS) index*

To summarize the response of identity neurons, we quantified the depth of selectivity (DOS) for each neuron: $DOS = \frac{n-(\sum_{j=1}^{n} r_j)/r_{max}}{n-1}$, where $n$ is the number of identities ($n = 50$), $r_j$ is the mean firing rate to identity $j$, and $r_{max}$ is the maximal mean firing rate across all identities. DOS varies from 0 to 1, with 0 indicating an equal response to all identities and 1 exclusive response to one identity, but not to any of the other identities. Thus, a DOS value of 1 is equal to maximal sparseness of identity coding. The DOS index has been used in many prior studies investigating visual selectivity [23,42,44].

*Population decoding of face identities*

We pooled all recorded neurons into a large pseudo-population. Firing rates were *z*-scored individually for each neuron to give equal weight to each unit regardless of firing rate. We used a maximal correlation coefficient classifier (MCC) as implemented in the MATLAB neural decoding toolbox (NDT) [45]. The MCC estimates a mean template for each class *i* and assigns the class for test trial. We used 8-fold cross-validation, i.e., all trials were randomly partitioned into 8 equal sized subsamples, of which 7 subsamples were used as the training data and the remaining single subsample was retained as the validation data for assessing the accuracy of the model, and this process was repeated 8 times, with each of the 8 subsamples used exactly once as the validation data. We then repeated the cross-validation procedure 50 times for different random train/test splits. Statistical significance of the decoding performance for each group of neurons against chance was estimated by calculating the percentage of bootstrap runs (50 in total) that had an accuracy below chance (i.e., 2% when decoding all identities). Statistical significance for comparing between groups of neurons was estimated by calculating the percentage of bootstrap runs (50 in total) that one group of neurons had a greater accuracy than the other. Spikes were counted in bins of 500 ms size and advanced by a step size of 50 ms. The first bin started −500 ms relative to trial onset (bin center was thus 250 ms before trial onset), and we tested 31 consecutive bins (the last bin was thus from 1000 ms to 1500 ms after trial onset). For each bin, a different classifier was trained/tested. For both tests, we used FDR [43] to correct for multiple comparisons across time points. The same decoding approach was used in our prior studies [46,47] and has been shown to be very effective in the study of neural population activity.

*Web-association score*

We employed a web-based association metric to study the relationship between different identities. To estimate the degree of relationship between the 50 celebrity identities, we used an internet search engine (Google) and compared the number of hits to the joint searches with the number of hits to the individual searches. The rationale is that the name of associated concepts will often appear together in web pages. The web-association score for each identity pair was calculated as

$a_{ij} = log_2\left(\frac{hits(identity_i AND identity_j)}{hits(identity_i) \cdot hits(identity_j)}\right)$. Because we used celebrity faces, all identities were well searchable from the internet and could thus give a reasonable number of hits to calculate web-association values. We lastly normalized the web-association value using *z*-scoring. Similar results were derived using the search engine Bing. The web-association score has been shown to be effective to reveal "universal associations" between identities in a previous study [7].

*Regression analyses*

To identify neurons that encoded a linear combination of facial features, we employed both a partial least squares (PLS) regression with DNN feature maps and a linear regression with the two dimensions of the t-SNE feature space. The PLS method has been shown to be effective to study the neural response to DNN features [48,49]. For PLS, we used 4 components for each layer (explaining at least 80% of variance; we selected the number of components with a 10-fold cross validation to minimize the prediction error). For both approaches, we used a permutation test with 1000 runs to determine whether a neuron encoded a significant face model (i.e., the neuron encoded the dimensions of the face space). In each run, we randomly shuffled the face labels and used 50% of the faces as the training dataset. We used the training dataset to construct a model (i.e., deriving regression coefficients), predicted responses using this model for each face in the remaining 50% of faces (i.e., test dataset), and computed the Pearson correlation between the predicted and actual response in the test dataset. The distribution of correlation coefficients computed *with* shuffling (i.e., null distribution) was eventually compared to the one *without* shuffling (i.e., observed response). If the correlation coefficient of the observed response was greater than 95% of the correlation coefficients from the null distribution, this face model was considered *significant*. This procedure has been shown to be very effective to select units with significant face models [50]. The correlation coefficient could also indicate the model's predictability and thus be compared between different neurons (**Extended Data Fig. 19g, h**). Similar results were derived using face models from [50] and [30].

*Representational similarity analysis (RSA)*

Dissimilarity matrices (DMs) [51] are symmetric matrices of dissimilarity between all pairs of face images or face identities. In a DM, larger values represent larger dissimilarity of pairs, such that the smallest value possible is the similarity of a condition to itself (dissimilarity of 0). We used the Pearson correlation to calculate DMs (firing rates were normalized to the mean baseline of each neuron), and we used the Spearman correlation to calculate the correspondence between the DMs (Spearman correlation was used because it does not assume a linear relationship [52]; Fisher $z$-transformation was performed on Pearson's $r$ to ensure that sample distribution was approximately normal). We further used permutation tests with 1000 runs to assess the significance of the correspondence between the MTL DM and the IT DM.

*Pairwise distances in the face space*

We employed a pairwise distance metric [17] to compare neural coding of face identities between human/monkey neurons and DNN units. For each pair of identities, we used the dissimilarity value (1−Pearson's $r$) [51] as a distance metric. The human/monkey neuronal distance metric was calculated between firing rates of all recorded neurons and the DNN distance metric was calculated between feature weights of all DNN units. For human neurons, we used the mean firing rate in a time window 250 ms to 1250 ms after stimulus onset as the response to each face [14]. For monkey neurons, we used the mean firing rate in a time window 70 ms to 180 ms after stimulus onset as the response to each face [20,21]. We then correlated the human/monkey neuronal distance metric and the DNN distance metric. To determine statistical significance, we used a non-parametric permutation test with 1000 runs. In each run, we randomly shuffled the face labels and calculated the correlation between the human/monkey neuronal distance metric and the DNN distance metric. The distribution of correlation coefficients computed *with* shuffling (i.e., null distribution) was eventually compared to the one *without* shuffling (i.e., observed response). If the correlation coefficient of the observed response was greater than 95% of the correlation coefficients from the null distribution, it was considered *significant*. A significant correlation indicated that the DNN face space had some correspondence to the human/monkey neuronal face space [17]. We computed the correlation for each DNN layer so that we could determine the specific layer to which the neuronal population most closely corresponded.

*Data availability*

All data that support the findings of this study are publicly available on OSF (https://osf.io/36kzc/).

**Extended Data Table 1.** List of patients.

| ID | Age | Sex | Race | Epilepsy diagnosis | Number of Amygdala Neurons | | | | | Number of Hippocampus Neurons | | | | |
|---|---|---|---|---|---|---|---|---|---|---|---|---|---|---|
| | | | | | Total | Left | Right | S-ID | M-ID | Total | Left | Right | S-ID | M-ID |
| P6 | 33 | F | Caucasian | Left posterior neocortical extratemporal / parietal | 9 | 9 | 0 | 0 | 0 | 31 | 31 | 0 | 5 | 4 |
| | | | | | 10 | 10 | 0 | 1 | 0 | 29 | 29 | 0 | 2 | 3 |
| P7 | 28 | F | Caucasian | Right mesial temporal | 10 | 10 | 0 | 0 | 4 | 34 | 34 | 0 | 4 | 4 |
| | | | | | 6 | 6 | 0 | 1 | 2 | 21 | 19 | 2 | 1 | 3 |
| | | | | | 6 | 6 | 0 | 0 | 1 | 20 | 19 | 1 | 0 | 3 |
| | | | | | 2 | 2 | 0 | 0 | 1 | 31 | 26 | 5 | 3 | 1 |
| P9 | 42 | M | Caucasian | Left frontal | 28 | 28 | 0 | 0 | 2 | 7 | 7 | 0 | 0 | 0 |
| | | | | | 29 | 29 | 0 | 0 | 0 | 7 | 7 | 0 | 0 | 1 |
| | | | | | 25 | 25 | 0 | 0 | 4 | 6 | 6 | 0 | 0 | 0 |
| | | | | | 23 | 23 | 0 | 1 | 1 | 3 | 3 | 0 | 0 | 0 |
| P10 | 47 | F | Caucasian | Right mesial temporal and neocortical temporal | 25 | 0 | 25 | 1 | 1 | 7 | 7 | 0 | 0 | 0 |
| | | | | | 24 | 0 | 24 | 0 | 0 | 27 | 27 | 0 | 1 | 1 |
| P11 | 33 | F | Caucasian | Right mesial temporal and extratemporal | 16 | 0 | 16 | 0 | 0 | 0 | 0 | 0 | 0 | 0 |
| | | | | | 14 | 0 | 14 | 0 | 0 | 0 | 0 | 0 | 0 | 0 |
| | | | | | 9 | 0 | 9 | 0 | 0 | 15 | 0 | 15 | 0 | 0 |
| | | | | | 6 | 0 | 6 | 0 | 0 | 10 | 0 | 10 | 0 | 1 |
| P13 | 41 | M | Caucasian | Left hippocampal | 1 | 1 | 0 | 0 | 0 | 3 | 3 | 0 | 0 | 0 |
| P14 | 26 | M | Caucasian | Bilateral amygdylar/hippocampal | 19 | 17 | 2 | 1 | 1 | 7 | 4 | 3 | 0 | 1 |
| | | | | | 24 | 22 | 2 | 2 | 2 | 8 | 5 | 3 | 0 | 1 |
| | | | | | 20 | 17 | 3 | 0 | 1 | 35 | 35 | 0 | 0 | 2 |
| | | | | | 10 | 8 | 2 | 0 | 1 | 7 | 5 | 2 | 2 | 0 |
| P15 | 37 | F | Caucasian | Left amygdylar/hippocampal | 12 | 12 | 0 | 1 | 1 | 13 | 0 | 13 | 1 | 1 |
| | | | | | 12 | 12 | 0 | 0 | 1 | 6 | 0 | 6 | 0 | 0 |
| P16 | 29 | F | Caucasian | Right temporal neocortex | 18 | 10 | 8 | 1 | 0 | 34 | 24 | 10 | 3 | 2 |
| | | | | | 28 | 19 | 9 | 0 | 5 | 46 | 32 | 14 | 3 | 5 |
| | | | | | 23 | 15 | 8 | 0 | 0 | 34 | 23 | 11 | 2 | 3 |
| | | | | | 27 | 15 | 12 | 1 | 2 | 37 | 25 | 12 | 0 | 3 |
| | | | | | 34 | 23 | 11 | 1 | 2 | 43 | 27 | 16 | 2 | 3 |
| | | | | | 23 | 13 | 10 | 0 | 1 | 28 | 19 | 9 | 0 | 1 |

| | | | | | | | | | | | | | | |
|---|---|---|---|---|---|---|---|---|---|---|---|---|---|---|
| P18 | 53 | F | Caucasian | Left temporal | 13 | 2 | 11 | 0 | 1 | 59 | 30 | 29 | 1 | 2 |
| | | | | | 59 | 22 | 37 | 1 | 0 | 19 | 11 | 8 | 1 | 1 |
| | | | | | 40 | 18 | 22 | 1 | 2 | 14 | 3 | 11 | 0 | 1 |
| | | | | | 34 | 18 | 16 | 1 | 1 | 10 | 1 | 9 | 0 | 1 |
| P19 | 49 | M | Caucasian | Right mesial temporal onset | 1 | 0 | 1 | 0 | 0 | 18 | 6 | 12 | 0 | 0 |
| | | | | | 0 | 0 | 0 | 0 | 0 | 14 | 6 | 8 | 2 | 0 |
| P20 | 31 | F | Caucasian | Left temporal | 43 | 43 | 0 | 0 | 2 | 60 | 57 | 3 | 3 | 6 |
| | | | | | 36 | 36 | 0 | 0 | 3 | 53 | 51 | 2 | 1 | 3 |
| | | | | | 34 | 34 | 0 | 0 | 1 | 28 | 27 | 1 | 1 | 0 |
| Sum | | | | | 340 | 237 | 103 | 8 | 23 | 327 | 267 | 60 | 19 | 26 |

Each row of neurons represents a separate recording session using the CelebA stimuli. Each session was recorded on a separate day. Total: all neurons recorded from an area. Left: neurons that were recorded from the left side of an area and had a firing rate greater than 0.15 Hz. Right: neurons that were recorded from the right side of an area and had a firing rate greater than 0.15 Hz. These neurons were included for further analysis. S-ID: identity neurons that encoded a single identity. M-ID: identity neurons that encoded multiple identities.

#### Extended Data Figure Legends

**Extended Data Fig. 1.** Spike sorting and recording quality assessment. **(a)** Histogram of the number of units identified on each active wire (only wires with at least one unit identified are counted). The average yield per wire with at least one unit was 2.63±1.56 (mean±SD). **(b)** Histogram of mean firing rates. **(c)** Histogram of proportion of inter-spike intervals (ISIs) which are shorter than 3 ms. The large majority of clusters had less than 0.5% of such short ISIs. **(d)** Histogram of the signal-to-noise ratio (SNR) of the mean waveform peak of each unit. **(e)** Histogram of the SNR of the entire waveform of all units. **(f)** Pairwise distance between all possible pairs of units on all wires where more than 1 cluster was isolated. Distances are expressed in units of standard deviation (SD) after normalizing the data such that the distribution of waveforms around their mean is equal to 1. **(g)** Isolation distance of all units for which this metric was defined ($n$ = 667, median = 15.22). Isolation distance was calculated based on [53,54]. If a cluster contains $n_C$ cluster spikes, the isolation distance of the cluster is the $D^2$ value of the $n_C$ th closest noise spike. Isolation distance is therefore the radius of the smallest ellipsoid from the cluster center containing all of the cluster spikes and an equal number of noise spikes. As such, isolation distance estimates how distant the cluster spikes are from the other spikes recorded on the same electrode. Isolation distance is not defined for cases in which the number of cluster spikes is greater than the number of noise spikes. **(h)** Single-identity (S-ID) and multiple-identity (M-ID) neurons did not differ significantly in isolation distance ($t(53) = 0.99$, P = 0.32).

**Extended Data Fig. 2.** Identity neurons. **(a)** Task. We employed a one-back task, in which patients responded whenever an identical famous face was repeated. Each face was presented for 1 second, followed by a jittered inter-stimulus-interval (ISI) of 0.5 to 0.75 seconds. **(b)** Sample stimuli. Faces from 50 celebrity identities were used for neural recordings. The identities were diverse in race, gender, and age, with a variety of facial expressions. All images had the same resolution, and the faces had a similar size and position in the images. **(c)** Percentage of single-identity (S-ID; shown in yellow) and multiple-identity (M-ID) neurons in the entire neuronal population. Stacked bar shows M-ID neurons that encoded visually similar identities (i.e., feature M-ID neurons; red) or not (i.e., non-feature M-ID neurons; blue). **(d)** Identity selectivity index. Both S-ID neurons ($n$ = 53; shown in yellow) and M-ID neurons ($n$ = 102; combining both feature M-ID and non-feature

M-ID neurons; shown in magenta) had a significantly higher identity selectivity index than non-identity neurons ($n$ = 1422). Error bars denote ±SEM across neurons. Asterisks indicate a significant difference using two-tailed unpaired $t$-test. ****: P < 0.0001. **(e)** Ordered average responses from the most- to the least-preferred identity. Non-identity neurons are shown for comparison purposes. Responses were normalized by the response to the most-preferred identity. Shaded areas denote ±SEM across neurons. The top bars indicate significant differences between S-ID/M-ID and non-identity neurons (two-tailed unpaired $t$-test, P < 0.05, corrected by FDR for Q < 0.05). **(f)** Difference in response ratio between the most-preferred and second most-preferred identities. **(g)** Depth of selectivity (DOS) index. Identity neurons had a significantly higher DOS index than non-identity neurons. Error bars denote ±SEM across neurons. Asterisks indicate a significant difference using two-tailed unpaired $t$-test. ****: P < 0.0001. **(h-j)** Population decoding of face identity. Shaded area denotes ±SEM across bootstraps. The horizontal dotted lines indicate the chance level. **(h)** Decoding performance was primarily driven by identity neurons (red). As expected, the response of identity neurons was informative only for a small subset of identities. The top bars illustrate the time points with a significant above-chance decoding performance (bootstrap, P < 0.05, corrected by FDR for Q < 0.05). **(i)** M-ID neurons had a significantly better decoding performance than S-ID neurons because the encoding by M-ID neurons was less sparse. The top bar illustrates the time points with a significant difference between M-ID and S-ID neurons (bootstrap, P < 0.05, corrected by FDR for Q < 0.05). **(j)** Decoding performance was better for familiar than unfamiliar identities. The top bar illustrates the time points with a significant difference between familiar and unfamiliar identities (bootstrap, P < 0.05, corrected by FDR for Q < 0.05). **(k)** Comparison of DNN full feature distance in feature M-ID neurons between selective-selective (S-S) identity pairs (i.e., neurons were selective to both identities; shown in red) vs. selective-non-selective (S-NS) identity pairs (i.e., neurons were selective to only one of the identities; shown in gray) using one-tailed one-sample Wilcoxon signed rank test. *: P < 0.05, **: P < 0.01, and ***: P < 0.001. Error bars denote ±SEM across neurons. Feature distance was calculated between identities using the average DNN full feature of faces from each identity. Feature distance was then normalized by the maximum DNN full feature distance separately for each layer. We first averaged the feature distance of all S-S identity pairs and all S-NS identity pairs for each feature M-ID neuron, and then we compared feature distance between S-S vs. S-NS identity pairs across neurons. We confirmed that the feature distance was significantly shorter for

S-S identity pairs in later DNN layers where we identified feature M-ID neurons, suggesting that feature M-ID neurons encoded identities that were clustered in these layers.

**Extended Data Fig. 3.** The deep neural network (DNN) used in this study. **(a)** Structure of the DNN. The convolutional neural network (CNN) consisted of a feature extraction section (13 convolutional layers) and a classification section (3 fully connected [FC] layers). The feature extraction section was consistent with the typical architecture of a CNN. A 3×3 filter with 1-pixel padding and 1-pixel stride was applied to each convolutional layer, which was followed by a Batch Normalization (BatchNorm) and Rectified Linear Unit (ReLU) operation. Some of the convolutional layers were followed by five 2×2 max-pool operations with a stride of 2. There were 3 FC layers in each classification section: the first two had 4096 channels each, and the third performed an *n*-way classification. Each FC layer was followed by a ReLU and 50% dropout to avoid overfitting. A nonlinear Softmax operation was applied to the final output of VGG-16 network to make the classification prediction of 50 identities. **(b)** Visualization of DNN features. **(c)** Feature correlation between faces. We calculated a correlation matrix of features between each face, grouped by individual identity. It is worth noting that in earlier layers, DNN features from faces of the same identity were not highly correlated, suggesting that these faces were not grouped together but distributed in the feature space (see also **Extended Data Fig. 4**). In later layers, DNN features from faces of the same identity were highly correlated, suggesting that these faces were clustered in the feature space. **(d)** Pairwise distance between faces in the full dimensional space was correlated with that in the t-SNE space. Shown are Pearson correlation coefficients *r*. Asterisk indicates a significant correlation in that layer ($P < 0.05$, Bonferroni-corrected for all layers).

**Extended Data Fig. 4.** Projection of faces onto feature space derived from each deep neural network (DNN) layer. Each color represents a different identity (names shown in the legend). The level of feature abstraction increased from earlier layers to later layers. Faces from the same identity were not clustered in earlier layers but became more clustered towards the top of the DNN hierarchy (later layers).

**Extended Data Fig. 5.** More example feature neurons from the deep neural network (DNN) layers FC6 and FC8. Legend conventions as in **Fig. 1**. It is worth noting that feature neurons encoded both Caucasian and African-American faces. On the one hand, different feature M-ID neurons from the same patient could encode different identities and covered different parts of the feature space (**Fig. 1a-c** vs. **Extended Data Fig. 5a**). On the other hand, feature M-ID neurons from different patients could encode a similar region of the feature space and cover the same identities (**Fig. 1a-c** vs. **Extended Data Fig. 5c**).

**Extended Data Fig. 6.** Illustration of the selection procedure of feature neurons using the two examples shown in **Fig. 1**. **(a-g)** Cell 57. **(h-n)** Cell 734. **(a, h)** Projection of the neuronal firing rate onto the feature space. **(b, i)** Density maps for observed distributions. **(c, j)** Density maps for permuted distributions. **(d, k)** The difference maps between observed and permuted distributions. **(e, f, l, m)** Statistically significant pixels identified by comparing the observed distribution to the permuted distribution. **(e, l)** Permutation test for each pixel: $P < 0.01$. **(f, m)** Permutation test for each pixel: $P < 0.05$, corrected by false discovery rate (FDR) [43]. **(g, n)** Identified regions after thresholding for the minimum number of pixels within the cluster. A mask (shown in magenta) was first applied to exclude pixels from the edges and corners where there were no faces because the regions with a small number of faces (i.e., the samples were sparse) were susceptible to false positives. Cluster size must be greater than 2.5% of the total number of pixels of the face space within the mask because small clusters were likely to be false positive. Note that correction by FDR led to similar results.

**Extended Data Fig. 7.** Population summary of neurons broken down for the amygdala and hippocampus. **(a-c)** Amygdala neurons. **(d-f)** Hippocampus neurons. **(a, d)** Percentage of single-identity (S-ID) and multiple-identity (M-ID) neurons. Stacked bar shows M-ID neurons that encoded visually similar identities (i.e., demonstrating feature-based coding; red) or not (blue). **(b, e)** The number of feature neurons identified from each deep neural network (DNN) layer. Blue: feature neurons that were also identity neurons. **(c, f)** The number of identity neurons in the whole population (left) and among feature neurons (right). Blue: the number of identity neurons. Red: the number of non-identity feature neurons. Gray: the number of non-identity neurons.

**Extended Data Fig. 8.** Analysis of low-level visual features. **(a-d)** Comparison of low-level visual features between images inside (selected) vs. outside (unselected) the feature M-ID neurons' tuning regions. For each neuron, we calculated a mean visual feature for selected vs. unselected images. Error bars denote ±SEM across neurons. **(a)** Itti-Koch saliency [2]. **(b)** Luminance (i.e., luminance after RGB values converted to the CIE 1976 *L**, *u**, *v** color space). **(c)** Contrast (RMS contrast, i.e., the standard deviation of the pixel intensities). **(d)** Wavelength (mean hue value after RGB converted to hue, saturation, and value [HSV]). **(e-g)** Comparison of layer-wise relevance propagation (LRP) [55] heat maps. **(e, f)** Average LRP heat maps across images. The scale bar (color bar) is in arbitrary units. Red: pixels that *positively* contributed to the classification (i.e., identity recognition). Blue: pixels that *negatively* contributed to the classification. **(e)** Selected identities. **(f)** Unselected identities. **(g)** Group difference heat map (selected − unselected) shows pixels that selected images used more than unselected images. A statistical map showing areas that had a significant difference in heat maps between selected vs. unselected images was overlaid (purple; two-tailed two-sample *t*-test between individual heat maps at each pixel, $P < 0.05$ uncorrected). **(h-k)** Ranking of face images by evoked firing rate. Each panel represents a feature M-ID neuron. (Upper) Sorted firing rate. (Middle) Face images were sorted by evoked firing rate. Top, middle, and bottom 5 face images are shown. We randomly selected faces that had the same firing rate (note that most faces in **(h)** and **(i)** had a zero firing rate; and we picked the same bottom 5 faces for **(h)** and **(i)** for a comparison with their different top 5 faces). (Lower) The corresponding LRP heat maps.

**Extended Data Fig. 9.** Control analyses for non-feature M-ID neurons. **(a, b)** Example non-feature M-ID neurons whose response was projected onto different feature spaces (i.e., from different DNNs other than the VGG-16). Legend conventions as in **Fig. 1**. **(a)** Feature space constructed by the AlexNet layer FC6. **(b)** Feature space constructed by the ResNet layer Res5c. **(c)** Percentage of non-feature M-ID neurons that became feature M-ID neurons in the AlexNet feature spaces. **(d)**

Percentage of non-feature M-ID neurons that became feature M-ID neurons in the ResNet feature spaces. **(e)** Comparison of visual similarity between selected vs. unselected identities and between feature M-ID vs. non-feature M-ID neurons. **(e)** Human rating of visual similarity. **(f)** Siamese score of visual similarity [5]. S-S: pairs of identities that a neuron was selective to. S-NS: pairs of identities where a neuron was selective to one of them but not selective (NS) to the other. Error bars denote ±SEM across neurons. Asterisks indicate a significant difference between S-S vs. S-NS pairs (two-tailed paired *t*-test) or between feature vs. non-feature M-ID neurons (two-tailed two-sample *t*-test): +: P < 0.1 and ****: P < 0.0001. n.s.: not significant.

**Extended Data Fig. 10.** Region-based feature coding in non-face stimuli. **(a, b)** Human neurons. **(c-j)** Artificial neural network units. **(a, b)** Two example neurons demonstrating region-based feature coding with object stimuli. Legend conventions as in **Fig. 1. (c, d, g, h)** Response of AlexNet units to 500 face stimuli (CelebA) and 500 object stimuli (ImageNet) in arbitrary units (a.u.). Error bars denote ±SEM across stimuli. **(e, f, i, j)** Projection of the AlexNet unit activation onto the VGG-Face feature space. Legend conventions as in **Fig. 1**. **(c-f)** Two example face-selective units demonstrating region-based feature coding. **(g-j)** Two example non-face-selective units demonstrating region-based feature coding.

**Extended Data Fig. 11.** Comparison between visual similarity and conceptual association. **(a)** Conceptual association rating for each pair of faces. **(b)** Visual similarity rating for each pair of faces. Color coding indicates the *z*-scored ratings within each patient. **(c)** Separate populations of M-ID neurons encoding visual features (i.e., visual similarity) and concepts (i.e., conceptual association). **(d)** DNN feature distance for each subpopulation of M-ID neurons. **(e)** Conceptual association ratings for each subpopulation of M-ID neurons. S-S: pairs of identities that a neuron was selective to. S-NS: pairs of identities where a neuron was selective to one of them but not selective (NS) to the other. Error bars denote ±SEM across neurons. Asterisks indicate a significant difference using two-tailed two-sample *t*-test: ***: P < 0.001 and ****: P < 0.0001. **(f)** Correlation between conceptual association ratings and web-association scores. Error bars denote ±SEM across participants ($n$ = 5 for patients and $n$ = 40 for general controls). **(g)** Correlation between conceptual association ratings and web-association scores across pairs of face identities. Ratings from the general controls were averaged across participants. Each dot represents a pair of face identities (*n* = 1225) and the gray line denotes the linear fit ($r(1255) = 0.16$, $P = 4.5\times10^{-8}$). **(h)** Correlation between general controls' visual similarity ratings and DNN feature similarity (i.e., the negative of the DNN feature distance) for each DNN layer. Solid circles represent a significant correlation (permutation test: $P < 0.05$; Bonferroni correction for multiple comparisons across DNN layers) and open circles represent a non-significant correlation. Shaded area denotes ±SD across permutation runs. **(i)** Correlation between visual similarity ratings and absolute difference in social trait judgment ratings. **(j)** Correlation between conceptual association ratings and absolute difference in social trait judgment ratings. Correlation was performed across all face pairs within each patient and then averaged across patients. Error bars denote ±SEM across patients (*n* = 5). Asterisks indicate a significant difference from 0 (two-tailed paired *t*-test; Bonferroni correction for multiple comparisons). **: $P < 0.01$. **(k)** Correlation of social trait ratings between neurosurgical patients and online participants from the general population (controls; *n* = 500). Shown are average correlation coefficients, and error bars denote ±SEM across patients (*n* = 6). Asterisks indicate a significant difference from 0 (two-tailed paired *t*-test; Bonferroni correction for multiple comparisons). **: $P < 0.01$, ***: $P < 0.001$, and ****: $P < 0.0001$.

**Extended Data Fig. 12.** Web-association matrix and correlation with deep neural network (DNN) features. **(a)** Web-association values between the 50 face identities. The color bar on the right shows the strength of association values in arbitrary units. **(b)** Correlation between the web-association score and DNN features. The web-association score for each identity pair was correlated with the Euclidean distance between their DNN features in each layer. Statistical significance was estimated by permutation testing: in each of the 1000 runs, labels of the web-association score were shuffled. P-values were calculated by comparing the observed correlation coefficient to those from the null distribution derived from the permutation. Web-association score did not correlate with any DNN features.

**Extended Data Fig. 13.** Feature-based coding with well-characterized identity neurons from a publicly available dataset [6]. Here, we restricted our analysis to identities that could be associated to individual faces (removing, for example, landmarks, animals, and music bands) while removing the concepts for which we could not compute an association score (*n* = 16 neurons). **(a)** An example neuron from layer FC6 that encoded visually similar and conceptually associated identities. Response-eliciting (i.e., selective) pictures are shown, while the remaining ones are represented by gray dots. **(b)** An example neuron from layer FC7 that only showed conceptual association but not feature-based coding (Prince William and his wife Kate Middleton). **(c)** An example neuron from layer FC6 that only showed feature-based coding but not conceptual association. **(d)** Another neuron from layer FC7 responding to Jim Carrey and John Terry, which were neither conceptually associated nor visually similar. **(e, f)** Normalized feature distance for each multiple identity (M-ID) neuron. Feature distance was calculated between images using the deep neural network (DNN) full feature of faces. Feature distance was then normalized by the maximum DNN full feature distance separately for layers FC6 and FC7. **(e)** Results from layer FC6. **(f)** Results from layer FC7. **(g)** Association score for M-ID neurons. For each neuron, we compared the normalized feature distance or association score between selective-selective (S-S) pairs (i.e., the neuron elicited a response to both pictures/identities; shown in red) vs. selective-non-selective (S-NS) pairs (i.e., the neuron elicited a response to only one of the pictures/identities; shown in gray) using a one-tailed one-sample Wilcoxon signed-rank test (i.e., only neurons that showed a significantly shorter feature distance or a significantly higher association for S-S pairs were detected). ***: $P < 0.001$ and ****: $P < 0.0001$. Error bars denote ±SEM across identity pairs. Note that if a neuron encoded more than one S-S pair, we averaged the feature distance and web-association score across S-S pairs for statistics. **(h)** Summary of each type of M-ID neuron (note that layers FC6 and FC7 revealed the same feature M-ID neurons). We found separate populations of M-ID neurons encoding visual features (i.e., visual similarity) and concepts (i.e., conceptual association).

**Extended Data Fig. 14.** Feature-based coding with well-characterized identity neurons from a publicly available dataset [6]. Here, we further restricted our analysis to famous faces only and compared the encoded identities with the CelebA stimuli (*n* = 11 neurons). **(a, b)** Two example neurons that encoded visually similar identities. We downloaded 8 images from the internet for the identities from the new dataset (shown in images) and projected these images onto our original CelebA feature space (shown in dots; each dot represents a face, and each color represents a different identity). **(c)** An example neuron that only showed conceptual association but not feature-based coding. **(d)** An example neuron that only showed feature-based coding but not conceptual association. **(e, f)** Normalized feature distance for each multiple identity (M-ID) neuron. Feature distance was calculated between identities using the average deep neural network (DNN) full feature of faces from each identity. Feature distance was then normalized by the maximum DNN full feature distance separately for layers FC6 and FC7. **(e)** Results from layer FC6. **(f)** Results from layer FC7. **(g)** Web-association score for M-ID neurons. For each neuron, we compared the normalized feature distance or web-association score between selective-selective (S-S) identity pairs (i.e., the neuron was selective to both identities; data shown in red) vs. selective-non-selective (S-NS) identity pairs (i.e., the neuron was selective to only one of the identities; data shown in gray) using one-tailed one-sample Wilcoxon signed-rank test (i.e., only neurons that showed a significantly shorter feature distance or a significantly higher web-association for S-S pairs were detected). *: $P < 0.05$, **: $P < 0.01$, ***: $P < 0.001$, and ****: $P < 0.0001$. Error bars denote ±SEM across identity pairs. Note that if a neuron encoded more than one S-S pair, we averaged the feature distance and web-association score across S-S pairs for statistics. **(h)** Summary of each type of M-ID neuron (note that layers FC6 and FC7 revealed the same feature M-ID neurons). We found separate populations of M-ID neurons encoding visual features (i.e., visual similarity) and concepts (i.e., conceptual association).

**Extended Data Fig. 15.** Projection of faces onto feature space derived using uniform manifold approximation and projection (UMAP). Each color represents a different identity (names shown in the legend). Although feature maps constructed by UMAP were different compared to those constructed by t-SNE (**Extended Data Fig. 4**), the relative distance between identities was largely preserved in these feature maps.

**Extended Data Fig. 16.** More example feature neurons from deep neural network (DNN) layers Conv5_2 and Conv5_3. Legend conventions as in **Fig. 1**. Feature neurons did not need to be identity neurons and they could appear in DNN layers where faces from the same identity were less clustered.

**Extended Data Fig. 17,** Specificity of the feature space and stimuli. **(a)** Projection of the CelebA stimuli onto the feature space constructed by the AlexNet layer FC6. **(b)** Projection of the CelebA stimuli onto the feature space constructed by the AlexNet layer FC7. **(c, d)** Projection of both CelebA face stimuli and ImageNet object stimuli onto the feature space constructed by the AlexNet layer FC6. **(e)** Projection of the CelebA stimuli onto the feature space constructed by the ResNet layer Res5b. **(f)** Projection of the CelebA stimuli onto the feature space constructed by the ResNet layer Res5c. **(g)** Projection of the CelebA stimuli onto the feature space constructed by the ResNet average pooling layer. **(h)** Projection of the CelebA stimuli onto the feature space constructed by the ResNet FC layer. **(i, j)** Projection of both CelebA face stimuli and ImageNet object stimuli onto the feature space constructed by the ResNet layer Res5c. **(k, l)** Projection of both CelebA face stimuli and ImageNet object stimuli onto the feature space constructed by the ResNet average pooling layer. Note that both AlexNet and ResNet were pre-trained for object recognition using ImageNet object stimuli. **(m, n)** Projection of both CelebA face stimuli and ImageNet object stimuli onto the original VGG-Face FC6 feature space. **(o, p)** Projection of both CelebA face stimuli and ImageNet object stimuli onto the original VGG-Face FC7 feature space. Dot: CelebA face stimuli. Diamond: ImageNet object stimuli. Each color represents a different identity or object category.

**Extended Data Fig. 18.** Additional results from the FBI Twins dataset. **(a, b)** Two example neurons demonstrating region-based feature coding in the FBI face space. **(c-e)** Common feature spaces for CelebA and FBI stimuli. The red outline delineates the tuning region of each example neuron. Note that in the face feature space combining the CelebA and FBI stimuli, the clustering based on gender and skin color was retained and similar to the feature spaces using the CelebA stimuli only. **(f-h)** Envelopes that include FBI stimuli within the vicinity of CelebA stimuli. We drew an envelope of the CelebA stimuli based on their distribution (shown in cyan contours): we first calculated a density map of the CelebA stimuli by smoothing a binary map of CelebA stimulus locations and then thresholded the density map to create the mask/envelope. **(f)** DNN layer Pool5. **(g)** DNN layer FC6. **(h)** DNN layer FC7. **(i)** Population results comparing neuronal response to FBI stimuli falling in vs. out of the tuning region ($n$ = 27). Each dot represents a neuron. Error bars denote ±SEM across neurons. Asterisks indicate a significant difference between In vs. Out responses using one-tailed paired *t*-test ($P < 0.01$).

**Extended Data Fig. 19.** Neurons responding to a linear combination of facial features. **(a, b)** Percentage of neurons showing a significant linear regression with the two t-SNE features (i.e., feature dimensions used to construct feature space for each layer). **(a)** MTL neurons. **(b)** IT MUA channels. Regression was performed individually for each DNN layer. Black dashed lines show the chance number of significant neurons/channels (5% of all recorded neurons/channels). Solid bars show conditions with an above-chance number of significant neurons/channels (binomial test: $P < 0.05$; after Bonferroni correction). **(c, d)** Example MTL neurons showing a significant partial least squares (PLS) regression with deep neural network (DNN) feature maps. Each dot represents a face identity, and the gray line denotes the linear fit. **(e, f)** Example MTL neurons showing a significant linear regression with the two t-SNE features. Note that these two example neurons are from **Fig. 1a-c** and **Extended Data Fig. 5b** (i.e., neurons demonstrating region-based feature coding), thus suggesting that region-based feature coding could lead to a significant linear regression (shown by the plane), because an elevated response in one part of the feature space could drive the regression. **(g, h)** Comparison of axis-based coding models between the MTL (red) and IT (blue). Model predictability was assessed using the Pearson correlation between the predicted and actual neural response in the test dataset. Error bars denote ±SEM across neurons/channels. **(g)** PLS regression model. **(h)** Linear regression model with the two t-SNE features.

**Extended Data Fig. 20.** Control analysis for the comparison of coding in the inferotemporal (IT) cortex and the medial temporal lobe (MTL). Legend conventions as in **Fig. 5**. **(a-e)** Analysis of IT multi-unit activity (MUA) using the first presentation of stimuli. **(f-m)** Analysis of MTL MUA. **(a, b, f, g)** The proportion of MUA channels demonstrating axis-based feature coding. **(c, h)**

Dissimilarity matrix (DM). Color coding shows dissimilarity values ($1-r$). **(d, i, j)** Observed vs. permuted correlation coefficient between MTL and IT DMs. **(i)** IT DM calculated using all presentations of stimuli. **(j)** IT DM calculated using the first presentation of stimuli only. **(e, k)** Correlation between pairwise distance in the neuronal face space and pairwise distance in the DNN face space. **(l)** Depth of selectivity (DOS) index. Error bars denote ±SEM across neurons/channels. Asterisks indicate a significant difference using two-tailed unpaired *t*-test. ****: $P < 0.0001$. **(m)** Ordered average responses from the most- to the least-preferred identity. Responses were normalized by the response to the most-preferred identity. Shaded areas denote ±SEM across channels. The top bar indicates a significant difference between MTL and IT identity channels (two-tailed unpaired *t*-test, $P < 0.05$, corrected by FDR for $Q < 0.05$). MTL channels showed a steeper change from the most- to the least-preferred identity compared to IT channels.

4 He, K., Zhang, X., Ren, S. & Sun, J. in *Proceedings of the IEEE conference on computer vision and pattern recognition.* 770-778.

5 Koch, G., Zemel, R. & Salakhutdinov, R. in *Proceedings of the 32nd International Conference on Machine Learning.*

6 Rey, H. G. *et al.* Encoding of long-term associations through neural unitization in the human medial temporal lobe. *Nature Communications* **9**, 4372, doi:10.1038/s41467-018-06870-2 (2018).

7 De Falco, E., Ison, M. J., Fried, I. & Quian Quiroga, R. Long-term coding of personal and universal associations underlying the memory web in the human brain. *Nature Communications* **7**, 13408, doi:10.1038/ncomms13408 (2016).

8 Deng, J. *et al.* in *2009 IEEE conference on computer vision and pattern recognition.* 248-255 (Ieee).

9 Rutishauser, U., Mamelak, A. N. & Schuman, E. M. Single-Trial Learning of Novel Stimuli by Individual Neurons of the Human Hippocampus-Amygdala Complex. *Neuron* **49**, 805-813, doi:10.1016/j.neuron.2006.02.015 (2006).

10 Brown, M. W. & Aggleton, J. P. Recognition memory: What are the roles of the perirhinal cortex and hippocampus? *Nature Reviews Neuroscience* **2**, 51-61, doi:10.1038/35049064 (2001).

11 Quian Quiroga, R. Concept cells: the building blocks of declarative memory functions. *Nature Reviews Neuroscience* **13**, 587, doi:10.1038/nrn3251 (2012).

12 Waydo, S., Kraskov, A., Quian Quiroga, R., Fried, I. & Koch, C. Sparse Representation in the Human Medial Temporal Lobe. *The Journal of Neuroscience* **26**, 10232, doi:10.1523/JNEUROSCI.2101-06.2006 (2006).

13 Wang, J., Cao, R., Brandmeir, N. J., Li, X. & Wang, S. Face identity selectivity in the deep neural network and human brain. *bioRxiv*, 2021.2005.2012.443829, doi:10.1101/2021.05.12.443829 (2021).

14 Mormann, F. *et al.* Latency and Selectivity of Single Neurons Indicate Hierarchical Processing in the Human Medial Temporal Lobe. *The Journal of Neuroscience* **28**, 8865-8872, doi:10.1523/jneurosci.1640-08.2008 (2008).

15 Allison, T., Puce, A., Spencer, D. D. & McCarthy, G. Electrophysiological Studies of Human Face Perception. I: Potentials Generated in Occipitotemporal Cortex by Face and Non-face Stimuli. *Cerebral Cortex* **9**, 415-430, doi:10.1093/cercor/9.5.415 (1999).

16 Ghuman, A. S. *et al.* Dynamic encoding of face information in the human fusiform gyrus. *Nature Communications* **5**, 5672, doi:10.1038/ncomms6672 (2014).

17 Grossman, S. *et al.* Convergent evolution of face spaces across human face-selective neuronal groups and deep convolutional networks. *Nature Communications* **10**, 4934, doi:10.1038/s41467-019-12623-6 (2019).

18 Decramer, T. *et al.* Single-Unit Recordings Reveal the Selectivity of a Human Face Area. *The Journal of Neuroscience* **41**, 9340, doi:10.1523/JNEUROSCI.0349-21.2021 (2021).
19 Khuvis, S. *et al.* Face-Selective Units in Human Ventral Temporal Cortex Reactivate during Free Recall. *The Journal of Neuroscience* **41**, 3386, doi:10.1523/JNEUROSCI.2918-19.2020 (2021).
20 Kar, K., Kubilius, J., Schmidt, K., Issa, E. B. & DiCarlo, J. J. Evidence that recurrent circuits are critical to the ventral stream's execution of core object recognition behavior. *Nature Neuroscience* **22**, 974-983, doi:10.1038/s41593-019-0392-5 (2019).
21 Kar, K. & DiCarlo, J. J. Fast Recurrent Processing via Ventrolateral Prefrontal Cortex Is Needed by the Primate Ventral Stream for Robust Core Visual Object Recognition. *Neuron* **109**, 164-176.e165, doi:https://doi.org/10.1016/j.neuron.2020.09.035 (2021).
22 Pessoa, L. & Adolphs, R. Emotion processing and the amygdala: from a 'low road' to 'many roads' of evaluating biological significance. *Nat Rev Neurosci* **11**, 773-783, doi:http://www.nature.com/nrn/journal/v11/n11/suppinfo/nrn2920_S1.html (2010).
23 Minxha, J. *et al.* Fixations Gate Species-Specific Responses to Free Viewing of Faces in the Human and Macaque Amygdala. *Cell Reports* **18**, 878-891, doi:http://dx.doi.org/10.1016/j.celrep.2016.12.083 (2017).
24 Squire, L. R., Stark, C. E. L. & Clark, R. E. The Medial Temporal Lobe. *Annual Review of Neuroscience* **27**, 279-306, doi:10.1146/annurev.neuro.27.070203.144130 (2004).
25 Squire, L. R. The Legacy of Patient H.M. for Neuroscience. *Neuron* **61**, 6-9, doi:10.1016/j.neuron.2008.12.023 (2009).
26 Cerf, M. *et al.* On-line, voluntary control of human temporal lobe neurons. *Nature* **467**, 1104, doi:10.1038/nature09510 https://www.nature.com/articles/nature09510#supplementary-information (2010).
27 Quian Quiroga, R., Reddy, L., Kreiman, G., Koch, C. & Fried, I. Invariant visual representation by single neurons in the human brain. *Nature* **435**, 1102-1107, doi:http://www.nature.com/nature/journal/v435/n7045/suppinfo/nature03687_S1.html (2005).
28 Rey, H. G. *et al.* Single Neuron Coding of Identity in the Human Hippocampal Formation. *Current Biology*, doi:https://doi.org/10.1016/j.cub.2020.01.035 (2020).
29 Liu, Z., Luo, P., Wang, X. & Tang, X. in *Proceedings of International Conference on Computer Vision (ICCV).*
30 Oosterhof, N. N. & Todorov, A. The functional basis of face evaluation. *Proceedings of the National Academy of Sciences* **105**, 11087-11092, doi:10.1073/pnas.0805664105 (2008).
31 Lin, C., Keles, U. & Adolphs, R. Four dimensions characterize attributions from faces using a representative set of English trait words. *Nature Communications* **12**, 5168, doi:10.1038/s41467-021-25500-y (2021).
32 Brainard, D. H. The Psychophysics Toolbox. *Spat Vis* **10**, 433-436 (1997).
33 Parkhi, O. M., Vedaldi, A. & Zisserman, A. Deep face recognition. (2015).
34 Hinton, G. E. & Roweis, S. T. Stochastic neighbor embedding. *Advances in neural information processing systems*, 857-864 (2003).
35 van der Maaten, L. & Hinton, G. Visualizing Data using t-SNE. *Journal of Machine Learning Research* **9**, 2579-2605 (2008).
36 Rutishauser, U., Ross, I. B., Mamelak, A. N. & Schuman, E. M. Human memory strength is predicted by theta-frequency phase-locking of single neurons. *Nature* **464**, 903-907,

doi:http://www.nature.com/nature/journal/v464/n7290/suppinfo/nature08860_S1.html (2010).
37 Rutishauser, U., Schuman, E. M. & Mamelak, A. N. Online detection and sorting of extracellularly recorded action potentials in human medial temporal lobe recordings, in vivo. *Journal of Neuroscience Methods* **154**, 204-224, doi:10.1016/j.jneumeth.2005.12.033 (2006).
38 Rutishauser, U. *et al.* Single-Unit Responses Selective for Whole Faces in the Human Amygdala. *Current biology : CB* **21**, 1654-1660 (2011).
39 Rutishauser, U. *et al.* Single-Neuron Correlates of Atypical Face Processing in Autism. *Neuron* **80**, 887-899 (2013).
40 Wang, S. *et al.* Neurons in the human amygdala selective for perceived emotion. *Proceedings of the National Academy of Sciences* **111**, E3110-E3119, doi:10.1073/pnas.1323342111 (2014).
41 Wang, S. *et al.* The human amygdala parametrically encodes the intensity of specific facial emotions and their categorical ambiguity. *Nature Communications* **8**, 14821, doi:10.1038/ncomms14821 https://www.nature.com/articles/ncomms14821#supplementary-information (2017).
42 Wang, S., Mamelak, A. N., Adolphs, R. & Rutishauser, U. Encoding of Target Detection during Visual Search by Single Neurons in the Human Brain. *Current Biology* **28**, 2058-2069.e2054, doi:https://doi.org/10.1016/j.cub.2018.04.092 (2018).
43 Benjamini, Y. & Hochberg, Y. Controlling the False Discovery Rate: A Practical and Powerful Approach to Multiple Testing. *Journal of the Royal Statistical Society. Series B (Methodological)* **57**, 289-300 (1995).
44 Rainer, G., Asaad, W. F. & Miller, E. K. Selective representation of relevant information by neurons in the primate prefrontal cortex. *Nature* **393**, 577-579 (1998).
45 Meyers, E. The Neural Decoding Toolbox. *Frontiers in Neuroinformatics* **7**, 1-12, doi:10.3389/fninf.2013.00008 (2013).
46 Rutishauser, U. *et al.* Representation of retrieval confidence by single neurons in the human medial temporal lobe. *Nat Neurosci* **18**, 1041-1050 (2015).
47 Wang, S., Mamelak, A. N., Adolphs, R. & Rutishauser, U. Abstract goal representation in visual search by neurons in the human pre-supplementary motor area. *Brain* **142**, 3530-3549, doi:10.1093/brain/awz279 (2019).
48 Yamins, D. L. K. *et al.* Performance-optimized hierarchical models predict neural responses in higher visual cortex. *Proceedings of the National Academy of Sciences* **111**, 8619, doi:10.1073/pnas.1403112111 (2014).
49 Ponce, C. R. *et al.* Evolving Images for Visual Neurons Using a Deep Generative Network Reveals Coding Principles and Neuronal Preferences. *Cell* **177**, 999-1009.e1010, doi:10.1016/j.cell.2019.04.005 (2019).
50 Chang, L. & Tsao, D. Y. The Code for Facial Identity in the Primate Brain. *Cell* **169**, 1013-1028.e1014, doi:10.1016/j.cell.2017.05.011 (2017).
51 Kriegeskorte, N., Mur, M. & Bandettini, P. Representational similarity analysis - connecting the branches of systems neuroscience. *Frontiers in Systems Neuroscience* **2**, doi:10.3389/neuro.06.004.2008 (2008).
52 Stolier, R. M. & Freeman, J. B. Neural pattern similarity reveals the inherent intersection of social categories. *Nat Neurosci* **19**, 795-797, doi:10.1038/nn.4296

http://www.nature.com/neuro/journal/v19/n6/abs/nn.4296.html#supplementary-information (2016).
53 Harris, K. D., Hirase, H., Leinekugel, X., Henze, D. A. & Buzsáki, G. Temporal Interaction between Single Spikes and Complex Spike Bursts in Hippocampal Pyramidal Cells. *Neuron* **32**, 141-149, doi:https://doi.org/10.1016/S0896-6273(01)00447-0 (2001).
54 Schmitzer-Torbert, N., Jackson, J., Henze, D., Harris, K. & Redish, A. D. Quantitative measures of cluster quality for use in extracellular recordings. *Neuroscience* **131**, 1-11, doi:https://doi.org/10.1016/j.neuroscience.2004.09.066 (2005).
55 Bach, S. *et al.* On Pixel-Wise Explanations for Non-Linear Classifier Decisions by Layer-Wise Relevance Propagation. *PLOS ONE* **10**, e0130140, doi:10.1371/journal.pone.0130140 (2015).

**Figure S1**

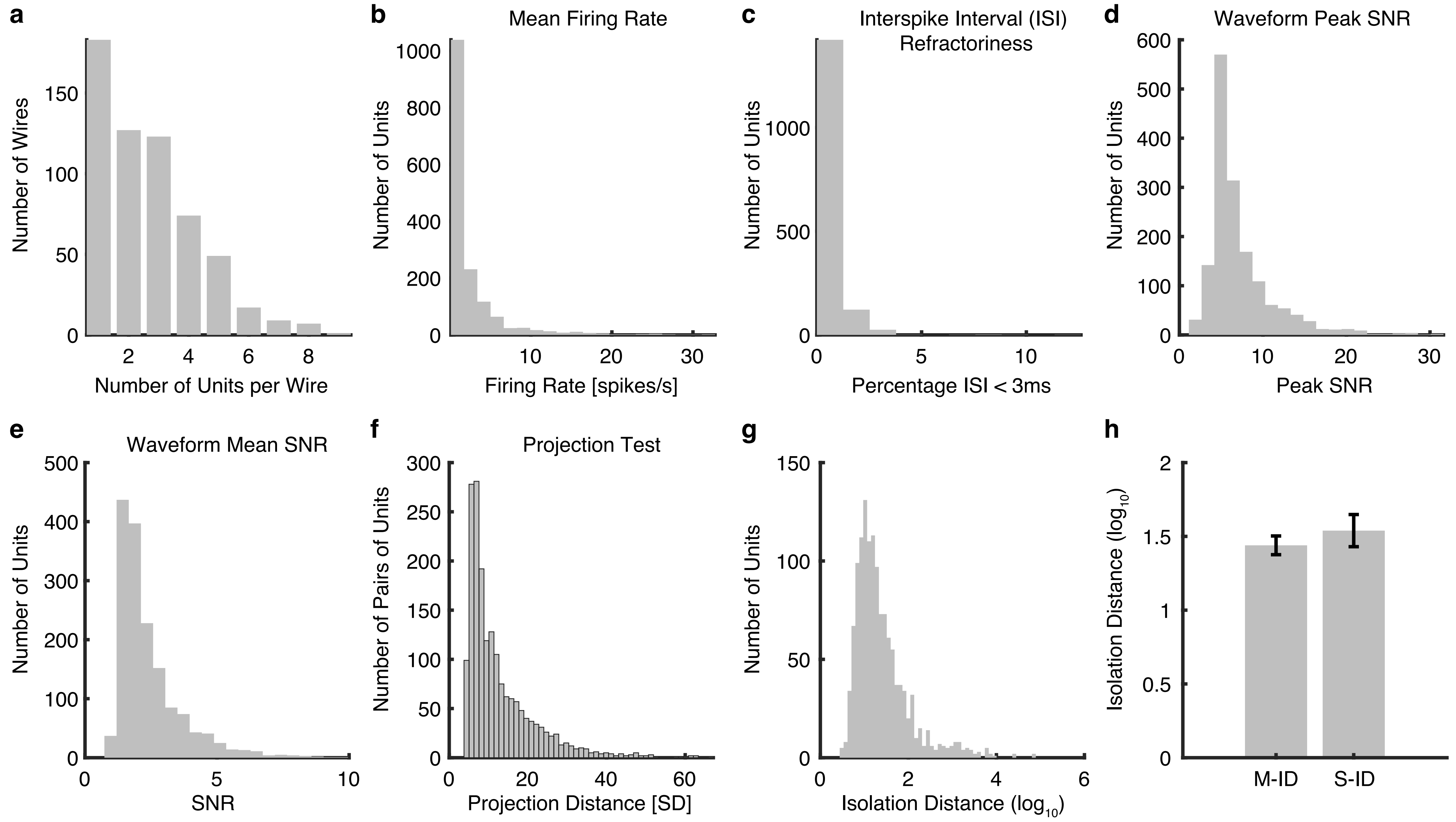

a
Number of Wires
Number of Units per Wire
b
Mean Firing Rate
Number of Units
Firing Rate [spikes/s]
c
Interspike Interval (ISI) Refractoriness
Number of Units
Percentage ISI < 3ms
d
Waveform Peak SNR
Number of Units
Peak SNR
e
Waveform Mean SNR
Number of Units
SNR
f
Projection Test
Number of Pairs of Units
Projection Distance [SD]
g
Number of Units
Isolation Distance ($log_{10}$)
h
Isolation Distance ($log_{10}$)
M-ID
S-ID

Figure S2

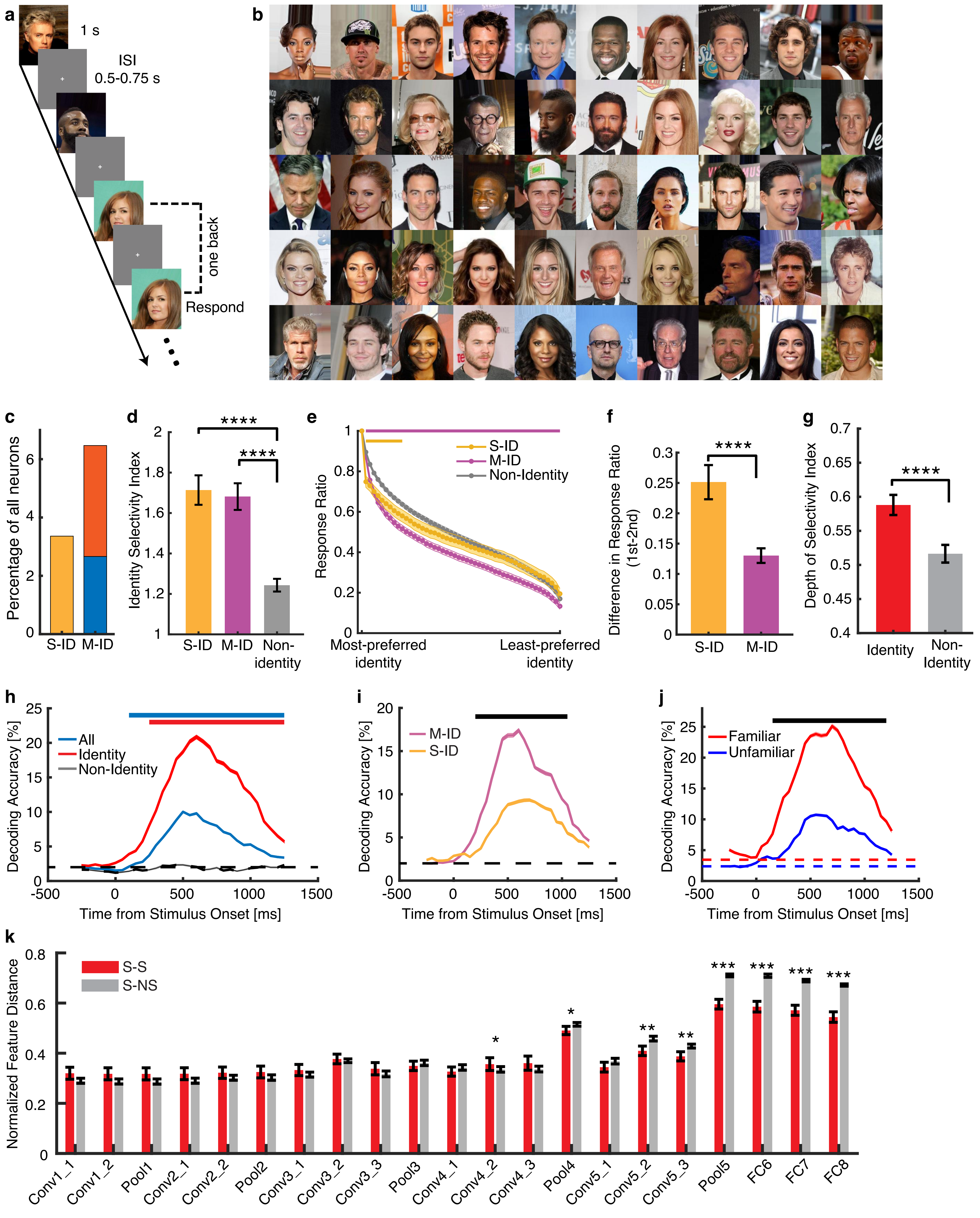
a
1 s
ISI
0.5-0.75 s
one back
Respond
b
c
Percentage of all neurons
S-ID
M-ID
d
****
****
Identity Selectivity Index
S-ID
M-ID
Non-identity
e
S-ID
M-ID
Non-Identity
Response Ratio
Most-preferred identity
Least-preferred identity
f
****
Difference in Response Ratio (1st-2nd)
S-ID
M-ID
g
****
Depth of Selectivity Index
Identity
Non-Identity
h
All
Identity
Non-Identity
Decoding Accuracy [%]
Time from Stimulus Onset [ms]
i
M-ID
S-ID
Decoding Accuracy [%]
Time from Stimulus Onset [ms]
j
Familiar
Unfamiliar
Decoding Accuracy [%]
Time from Stimulus Onset [ms]
k
S-S
S-NS
Normalized Feature Distance
Conv1_1
Conv1_2
Pool1
Conv2_1
Conv2_2
Pool2
Conv3_1
Conv3_2
Conv3_3
Pool3
Conv4_1
Conv4_2
Conv4_3
Pool4
Conv5_1
Conv5_2
Conv5_3
Pool5
FC6
FC7
FC8

Figure S3

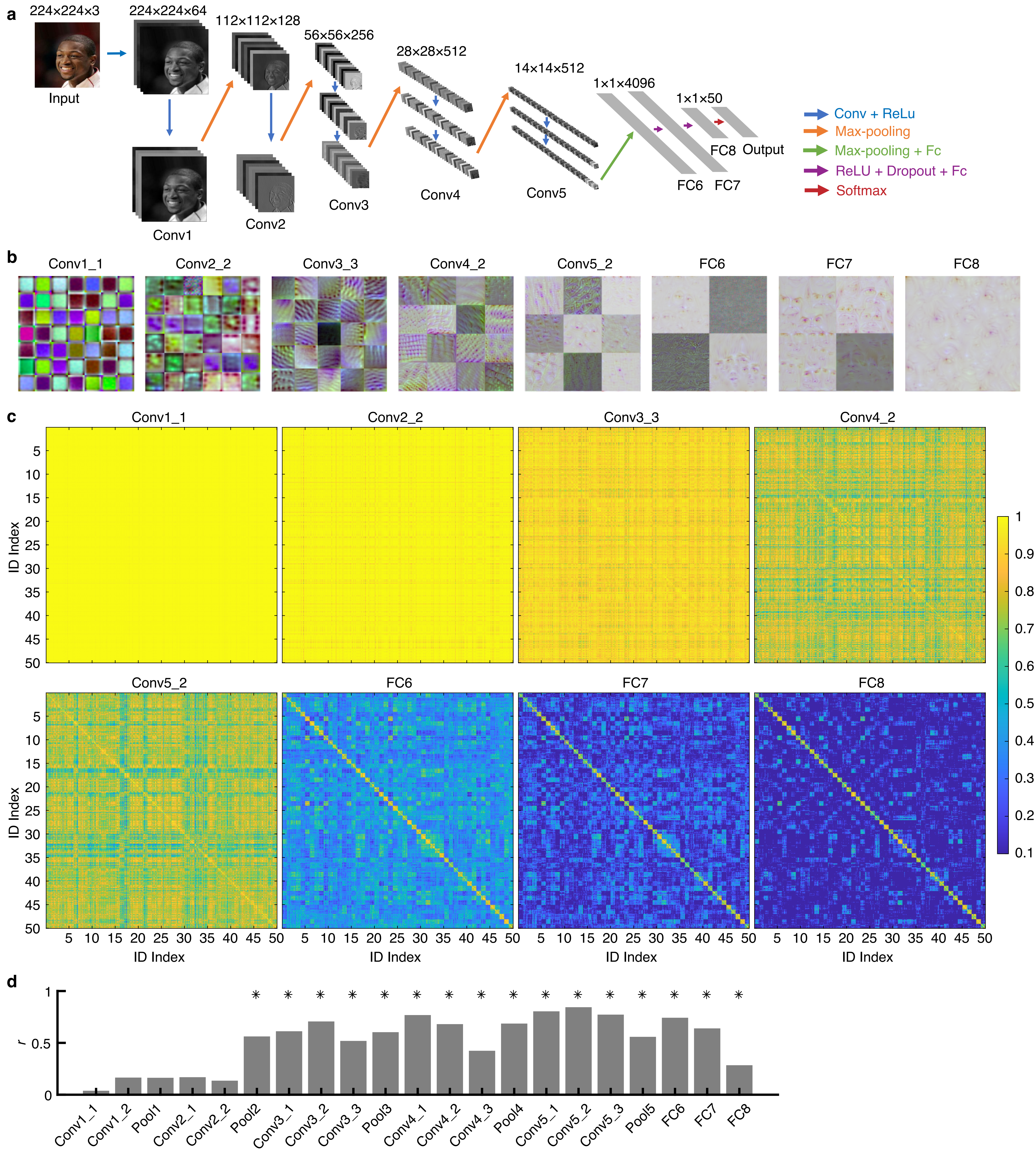
a
224×224×3
Input
224×224×64
Conv1
112×112×128
Conv2
56×56×256
Conv3
28×28×512
Conv4
14×14×512
Conv5
1×1×4096
FC6
FC7
1×1×50
FC8
Output
Conv + ReLu
Max-pooling
Max-pooling + Fc
ReLU + Dropout + Fc
Softmax
b
Conv1_1
Conv2_2
Conv3_3
Conv4_2
Conv5_2
FC6
FC7
FC8
c
Conv1_1
Conv2_2
Conv3_3
Conv4_2
Conv5_2
FC6
FC7
FC8
ID Index
ID Index
5 10 15 20 25 30 35 40 45 50
1 0.9 0.8 0.7 0.6 0.5 0.4 0.3 0.2 0.1
d
r
0 0.5 1
Conv1_1 Conv1_2 Pool1 Conv2_1 Conv2_2 Pool2 Conv3_1 Conv3_2 Conv3_3 Pool3 Conv4_1 Conv4_2 Conv4_3 Pool4 Conv5_1 Conv5_2 Conv5_3 Pool5 FC6 FC7 FC8

**Figure S4**

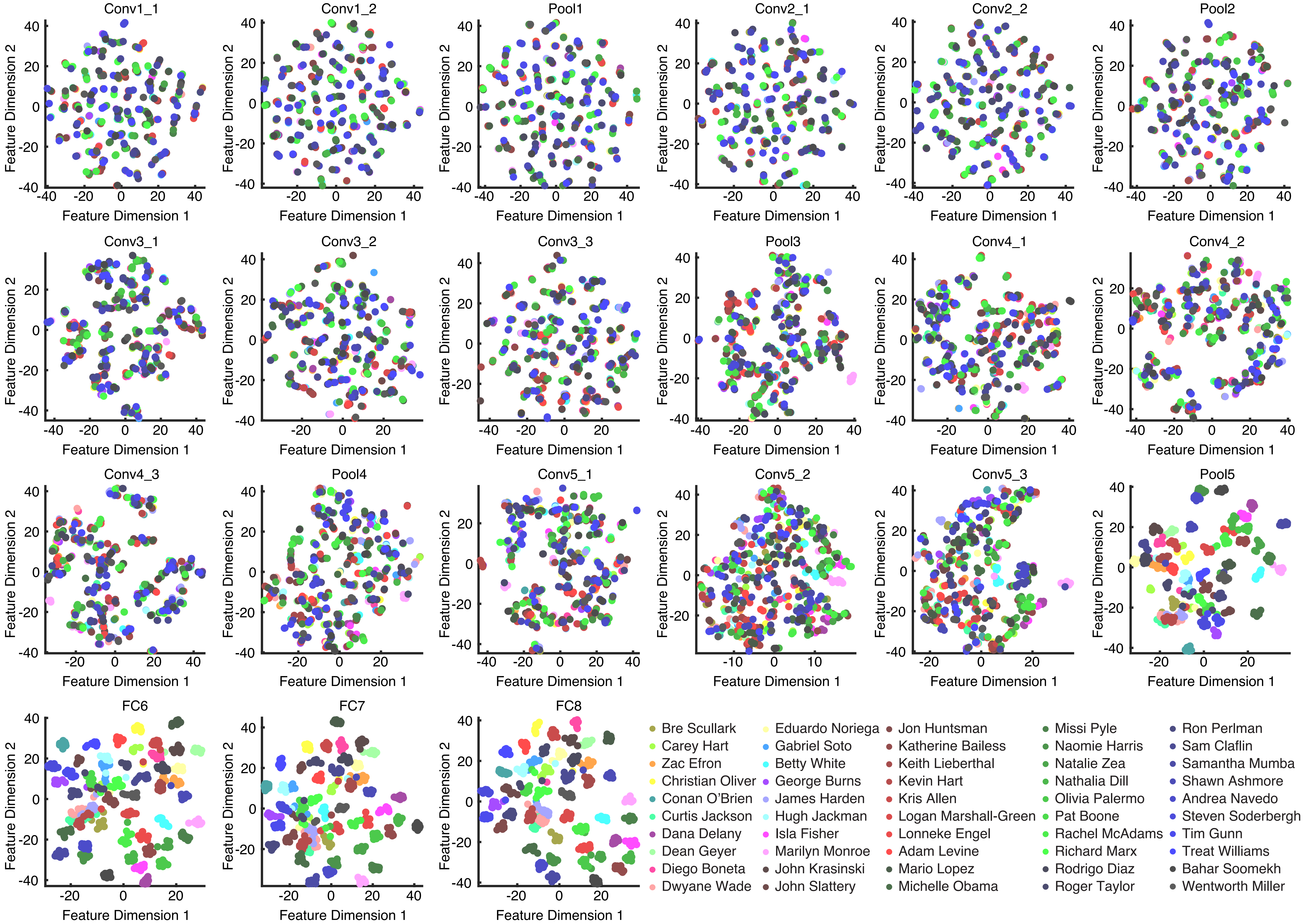

Conv1_1
Conv1_2
Pool1
Conv2_1
Conv2_2
Pool2
Conv3_1
Conv3_2
Conv3_3
Pool3
Conv4_1
Conv4_2
Conv4_3
Pool4
Conv5_1
Conv5_2
Conv5_3
Pool5
FC6
FC7
FC8
Feature Dimension 1
Feature Dimension 2
Bre Scullark
Carey Hart
Zac Efron
Christian Oliver
Conan O'Brien
Curtis Jackson
Dana Delany
Dean Geyer
Diego Boneta
Dwyane Wade
Eduardo Noriega
Gabriel Soto
Betty White
George Burns
James Harden
Hugh Jackman
Isla Fisher
Marilyn Monroe
John Krasinski
John Slattery
Jon Huntsman
Katherine Bailess
Keith Lieberthal
Kevin Hart
Kris Allen
Logan Marshall-Green
Lonneke Engel
Adam Levine
Mario Lopez
Michelle Obama
Missi Pyle
Naomie Harris
Natalie Zea
Nathalia Dill
Olivia Palermo
Pat Boone
Rachel McAdams
Richard Marx
Rodrigo Diaz
Roger Taylor
Ron Perlman
Sam Claflin
Samantha Mumba
Shawn Ashmore
Andrea Navedo
Steven Soderbergh
Tim Gunn
Treat Williams
Bahar Soomekh
Wentworth Miller

**Figure S5**

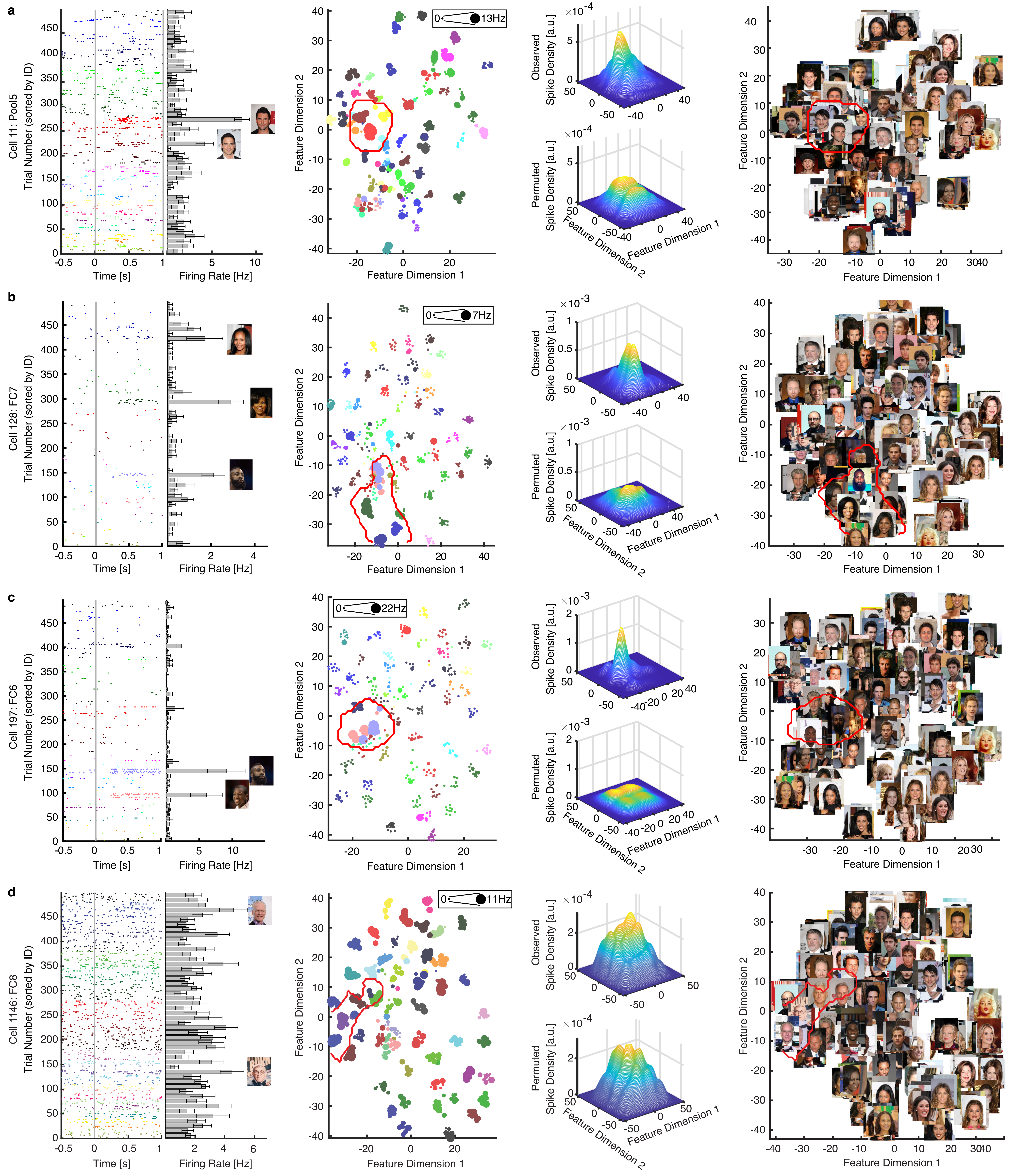
a
Cell 11: Pool5
Trial Number (sorted by ID)
Time [s]
Firing Rate [Hz]
Feature Dimension 1
Feature Dimension 2
0
13Hz
Observed
Spike Density [a.u.]
$\times 10^{-4}$
Permuted
Spike Density [a.u.]
b
Cell 128: FC7
Trial Number (sorted by ID)
Time [s]
Firing Rate [Hz]
7Hz
$\times 10^{-3}$
c
Cell 197: FC6
Trial Number (sorted by ID)
Time [s]
Firing Rate [Hz]
22Hz
$\times 10^{-3}$
d
Cell 1146: FC8
Trial Number (sorted by ID)
Time [s]
Firing Rate [Hz]
11Hz
$\times 10^{-4}$

**Figure S6**

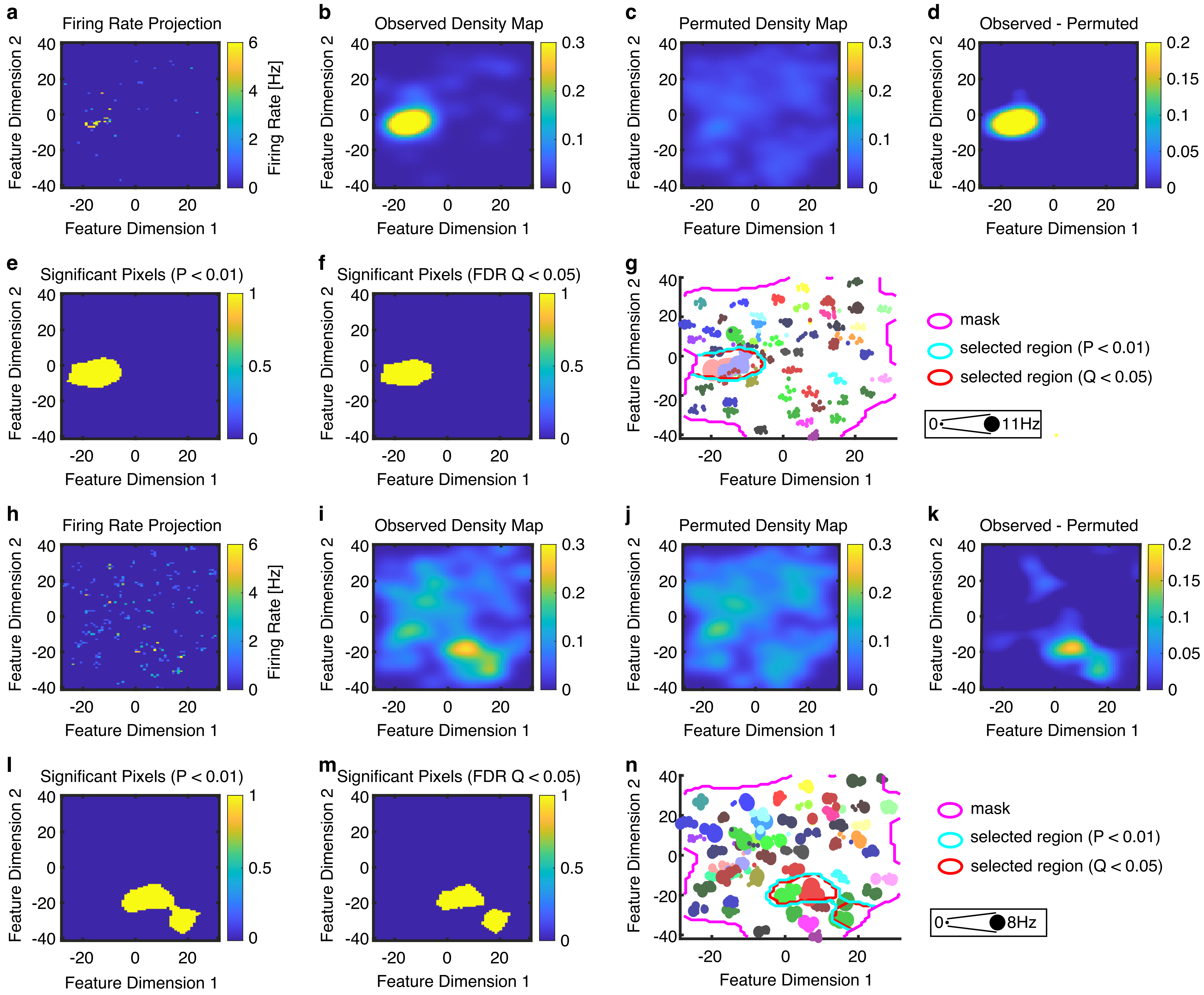
a
Firing Rate Projection
Firing Rate [Hz]
b
Observed Density Map
c
Permuted Density Map
d
Observed - Permuted
e
Significant Pixels (P < 0.01)
f
Significant Pixels (FDR Q < 0.05)
g
mask
selected region (P < 0.01)
selected region (Q < 0.05)
0
11Hz
h
Firing Rate Projection
Firing Rate [Hz]
i
Observed Density Map
j
Permuted Density Map
k
Observed - Permuted
l
Significant Pixels (P < 0.01)
m
Significant Pixels (FDR Q < 0.05)
n
mask
selected region (P < 0.01)
selected region (Q < 0.05)
0
8Hz
Feature Dimension 1
Feature Dimension 2

**Figure S7**

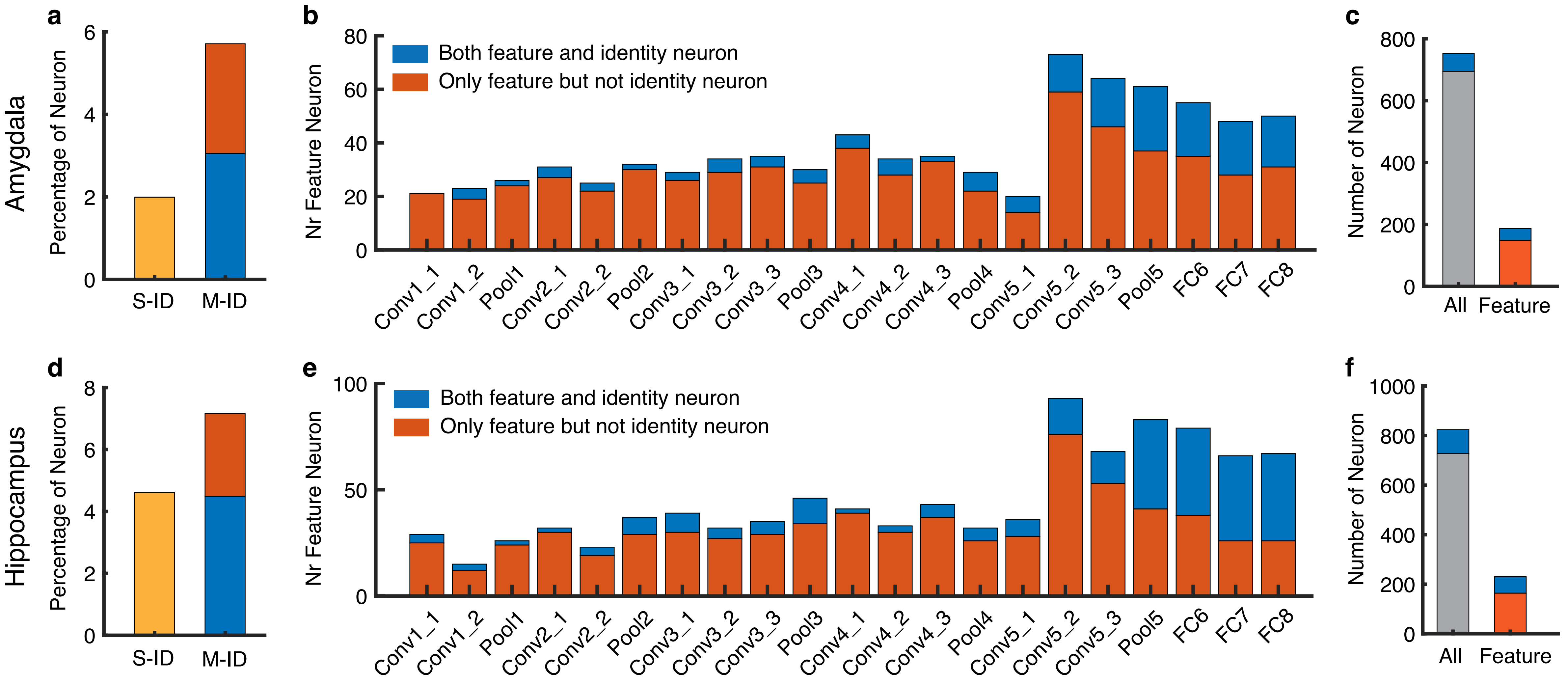
a
b
c
d
e
f
Amygdala
Hippocampus
Percentage of Neuron
S-ID
M-ID
Nr Feature Neuron
Both feature and identity neuron
Only feature but not identity neuron
Conv1_1
Conv1_2
Pool1
Conv2_1
Conv2_2
Pool2
Conv3_1
Conv3_2
Conv3_3
Pool3
Conv4_1
Conv4_2
Conv4_3
Pool4
Conv5_1
Conv5_2
Conv5_3
Pool5
FC6
FC7
FC8
Number of Neuron
All
Feature

**Figure S8**

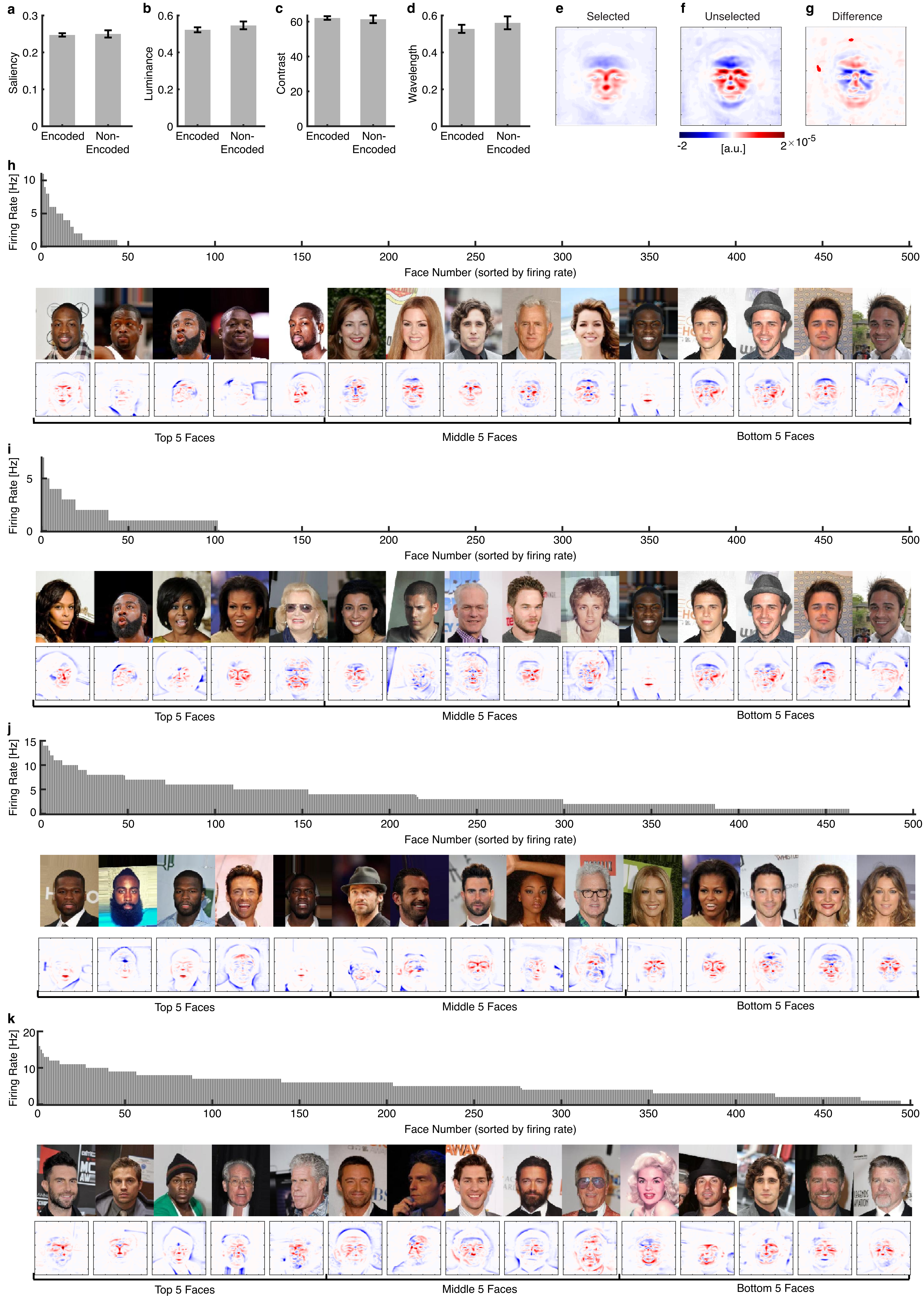

a
Saliency
Encoded
Non-Encoded
b
Luminance
c
Contrast
d
Wavelength
e
Selected
f
Unselected
g
Difference
[a.u.]
h
Firing Rate [Hz]
Face Number (sorted by firing rate)
Top 5 Faces
Middle 5 Faces
Bottom 5 Faces
i
j
k

#### Figure S9

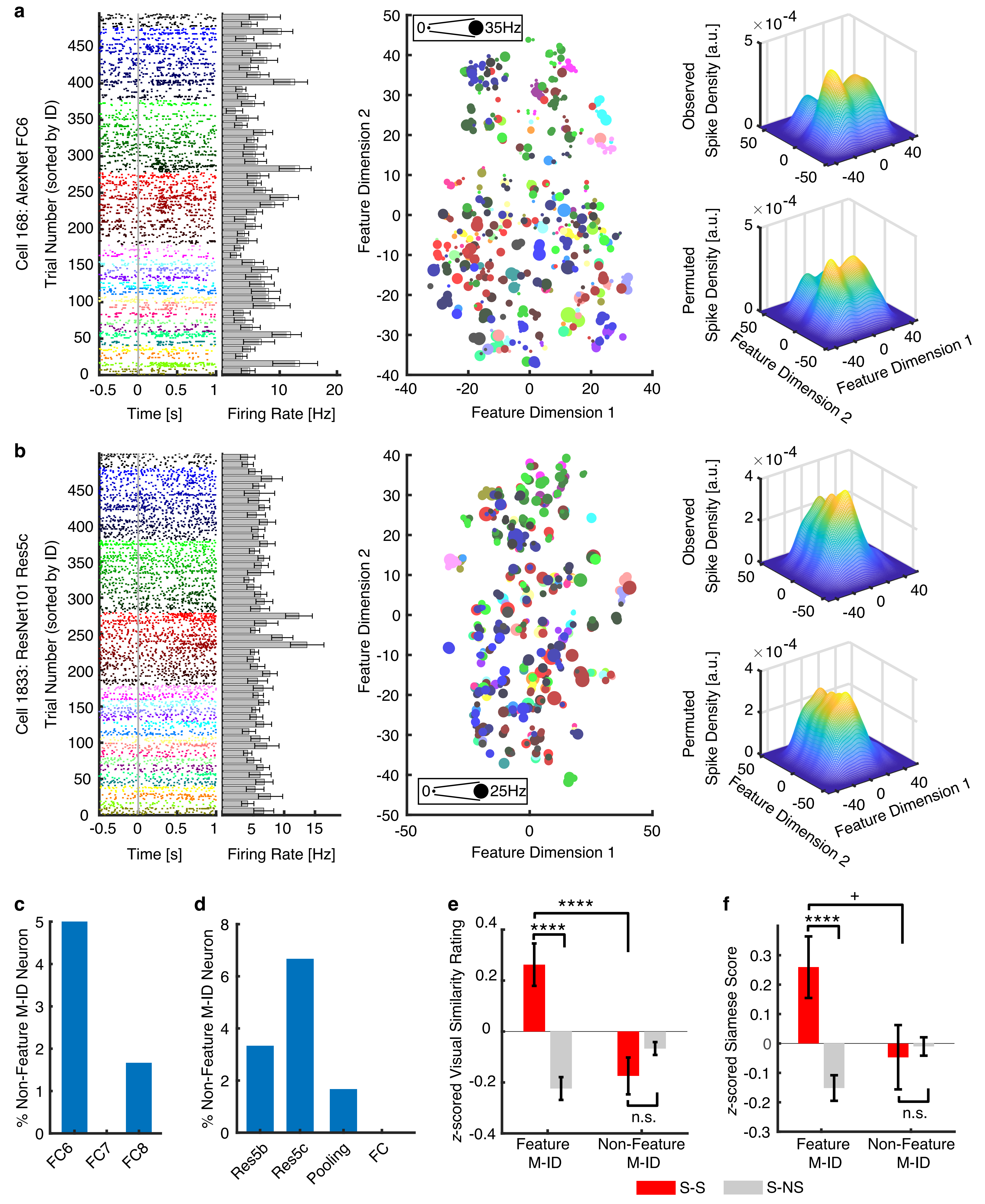
a
Cell 168: AlexNet FC6
Trial Number (sorted by ID)
Time [s]
Firing Rate [Hz]
0
35Hz
Feature Dimension 1
Feature Dimension 2
Observed
Permuted
Spike Density [a.u.]
×10^-4
b
Cell 1833: ResNet101 Res5c
25Hz
c
% Non-Feature M-ID Neuron
FC6
FC7
FC8
d
Res5b
Res5c
Pooling
FC
e
z-scored Visual Similarity Rating
****
n.s.
Feature M-ID
Non-Feature M-ID
f
z-scored Siamese Score
+
S-S
S-NS

**Figure S10**

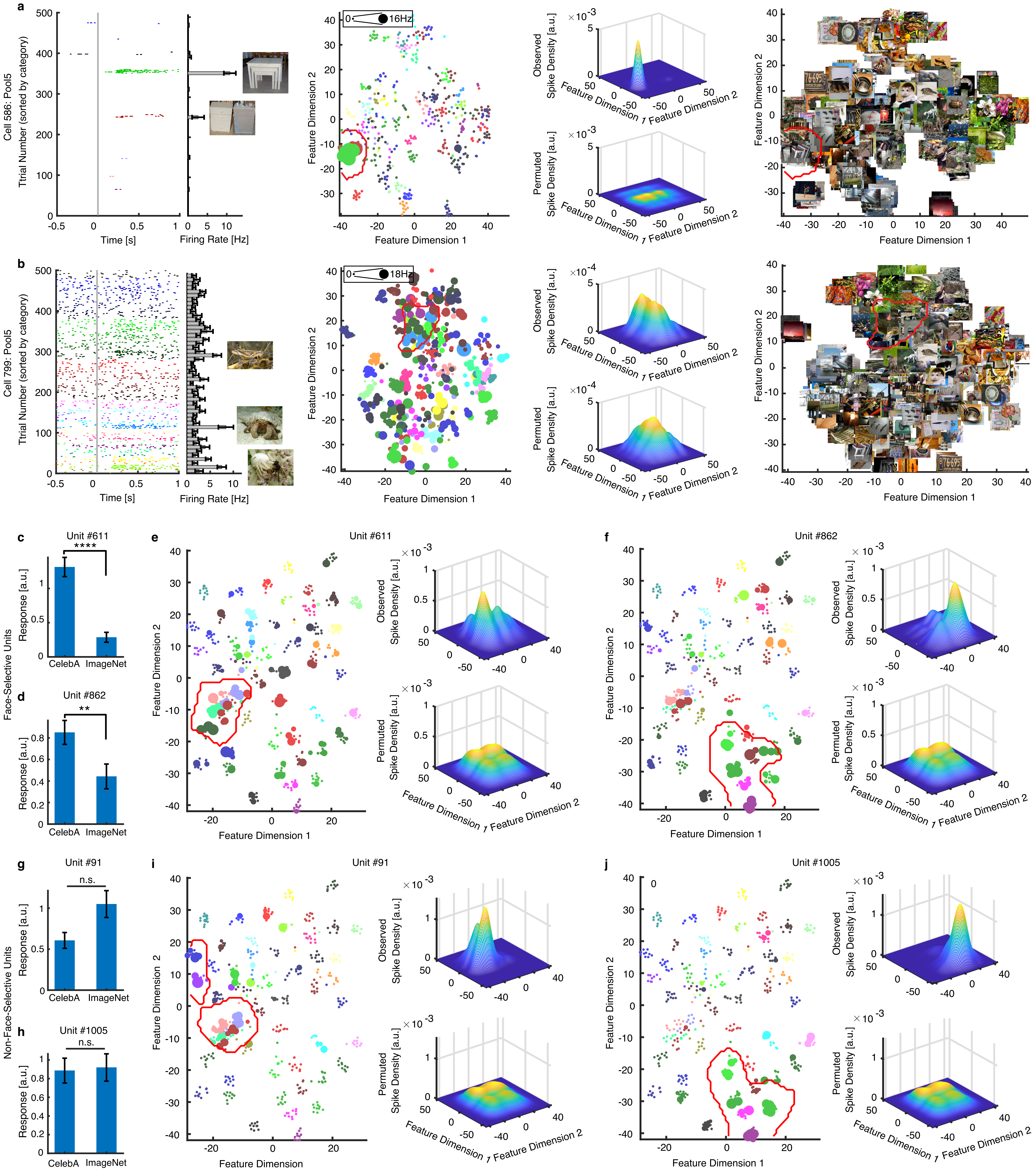
a
Cell 586: Pool5
Ttrial Number (sorted by category)
Time [s]
Firing Rate [Hz]
16Hz
Feature Dimension 1
Feature Dimension 2
Observed Spike Density [a.u.]
Permuted Spike Density [a.u.]
b
Cell 799: Pool5
18Hz
c
Unit #611
****
CelebA
ImageNet
Response [a.u.]
Face-Selective Units
d
Unit #862
**
e
f
g
Unit #91
n.s.
Non-Face-Selective Units
h
Unit #1005
i
j

**Figure S11**

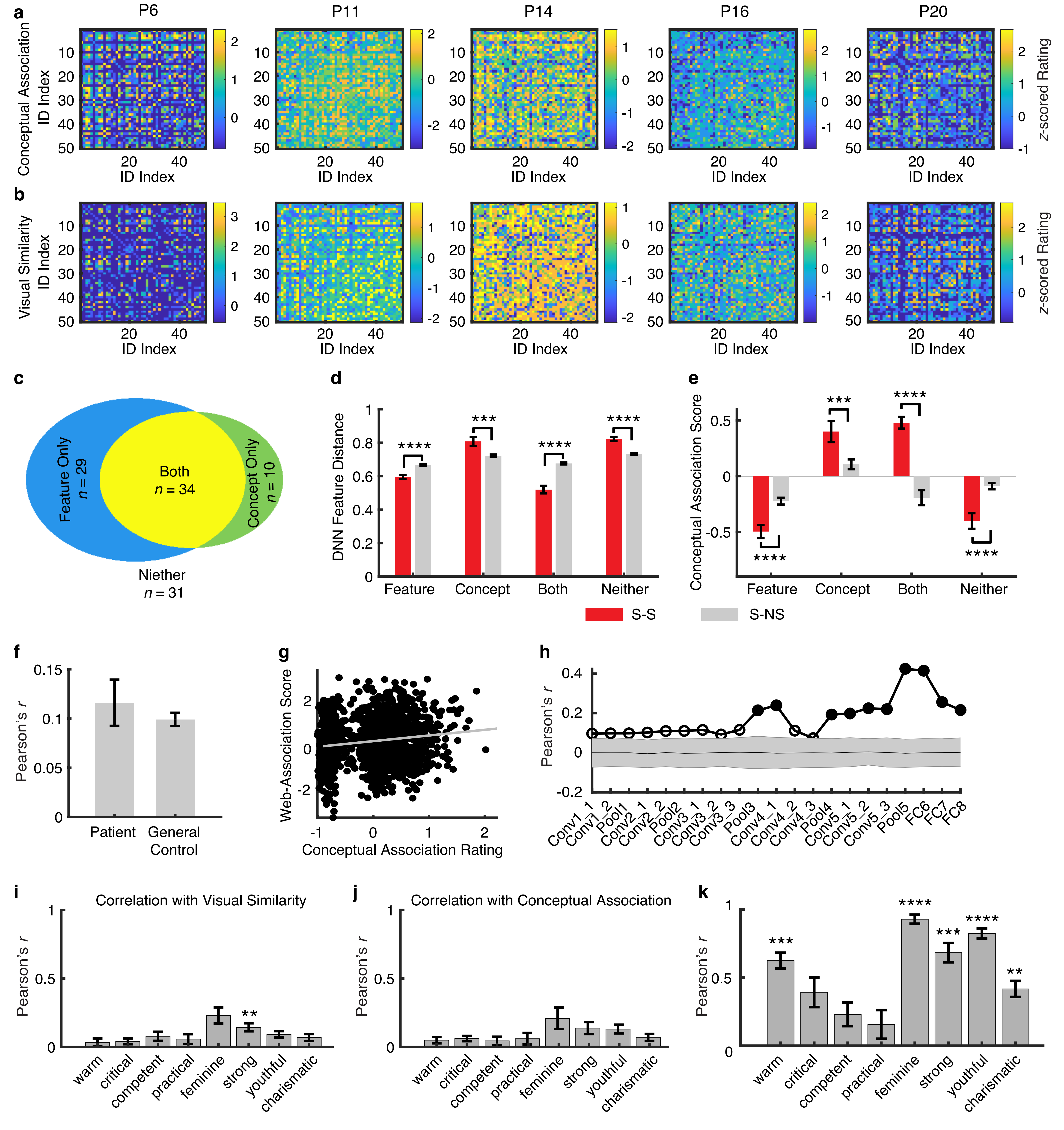
a
b
c
d
e
f
g
h
i
j
k
P6
P11
P14
P16
P20
Conceptual Association ID Index
Visual Similarity ID Index
ID Index
z-scored Rating
Feature Only n = 29
Both n = 34
Concept Only n = 10
Niether n = 31
DNN Feature Distance
Conceptual Association Score
Feature
Concept
Both
Neither
S-S
S-NS
Pearson's r
Patient
General Control
Web-Association Score
Conceptual Association Rating
Conv1_1
Conv1_2
Pool1
Conv2_1
Conv2_2
Pool2
Conv3_1
Conv3_2
Conv3_3
Pool3
Conv4_1
Conv4_2
Conv4_3
Pool4
Conv5_1
Conv5_2
Conv5_3
Pool5
FC6
FC7
FC8
Correlation with Visual Similarity
Correlation with Conceptual Association
warm
critical
competent
practical
feminine
strong
youthful
charismatic

**Figure S12**

**a**

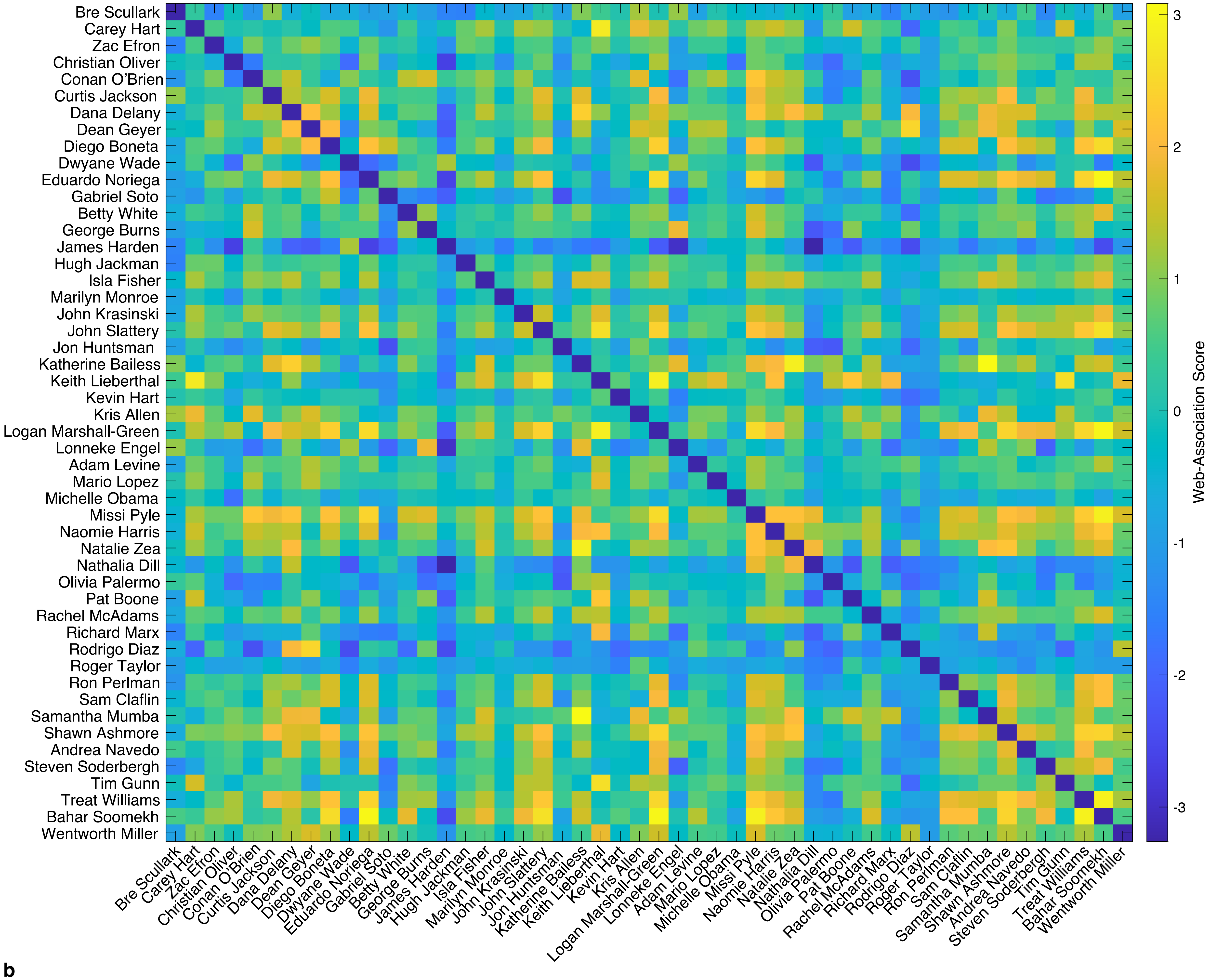


**b**

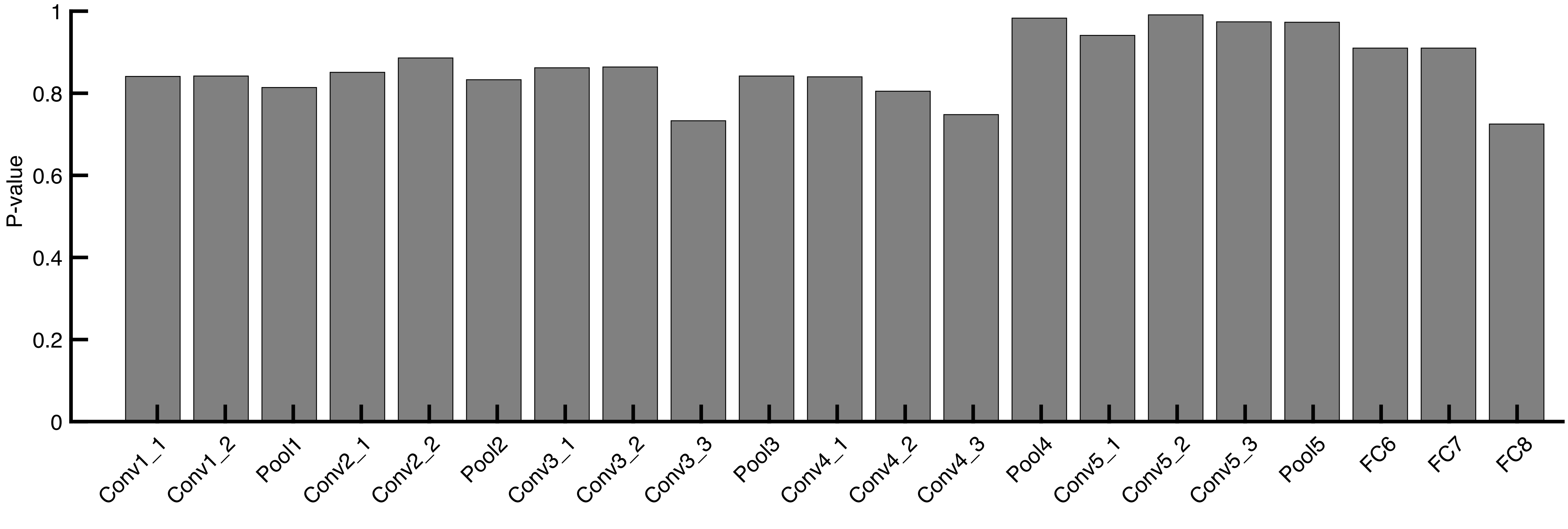

**Figure S13**

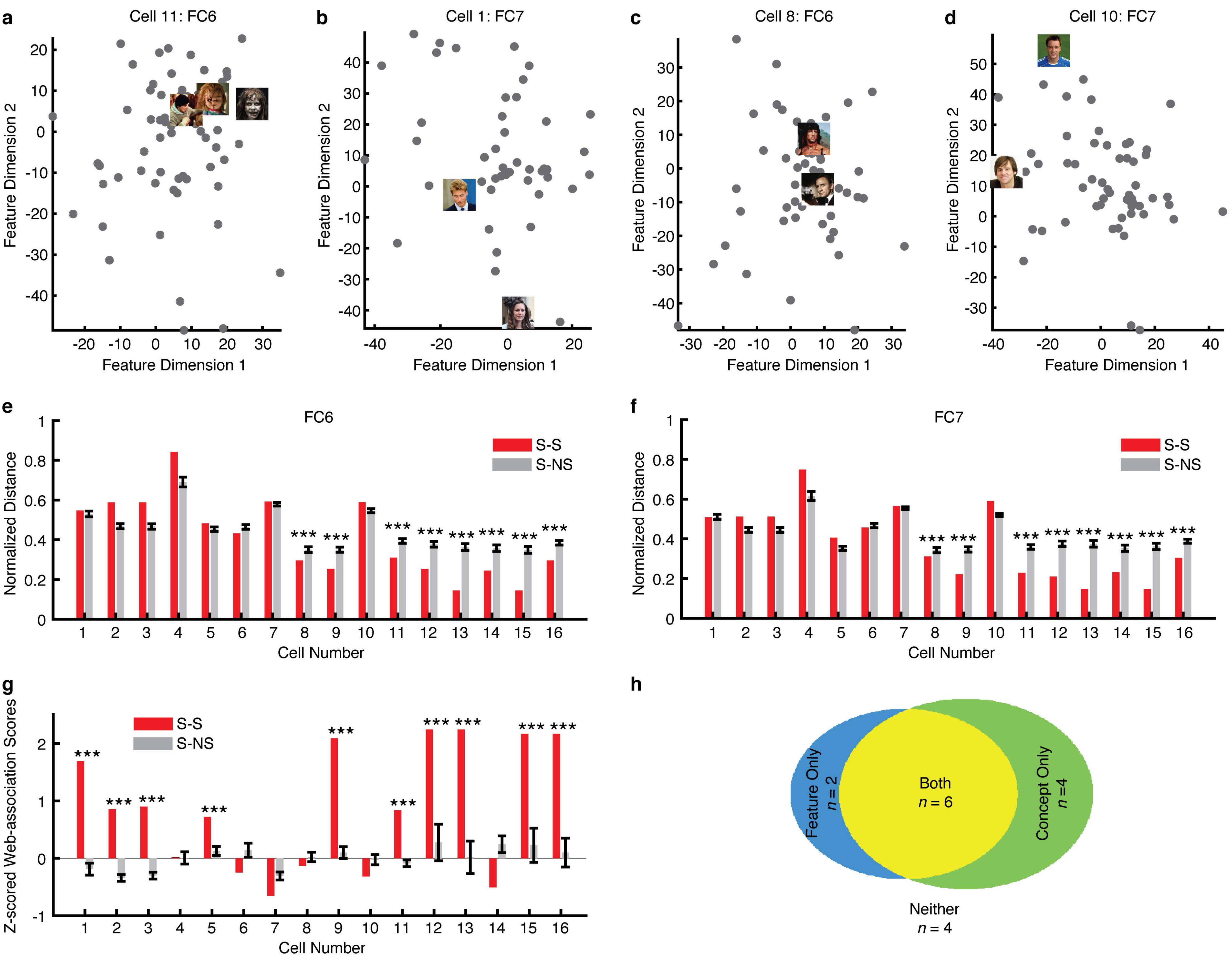
a
Cell 11: FC6
b
Cell 1: FC7
c
Cell 8: FC6
d
Cell 10: FC7
Feature Dimension 1
Feature Dimension 2
e
FC6
f
FC7
S-S
S-NS
Normalized Distance
Cell Number
***
g
Z-scored Web-association Scores
h
Feature Only
n = 2
Both
n = 6
Concept Only
n =4
Neither
n = 4

**Figure S14**

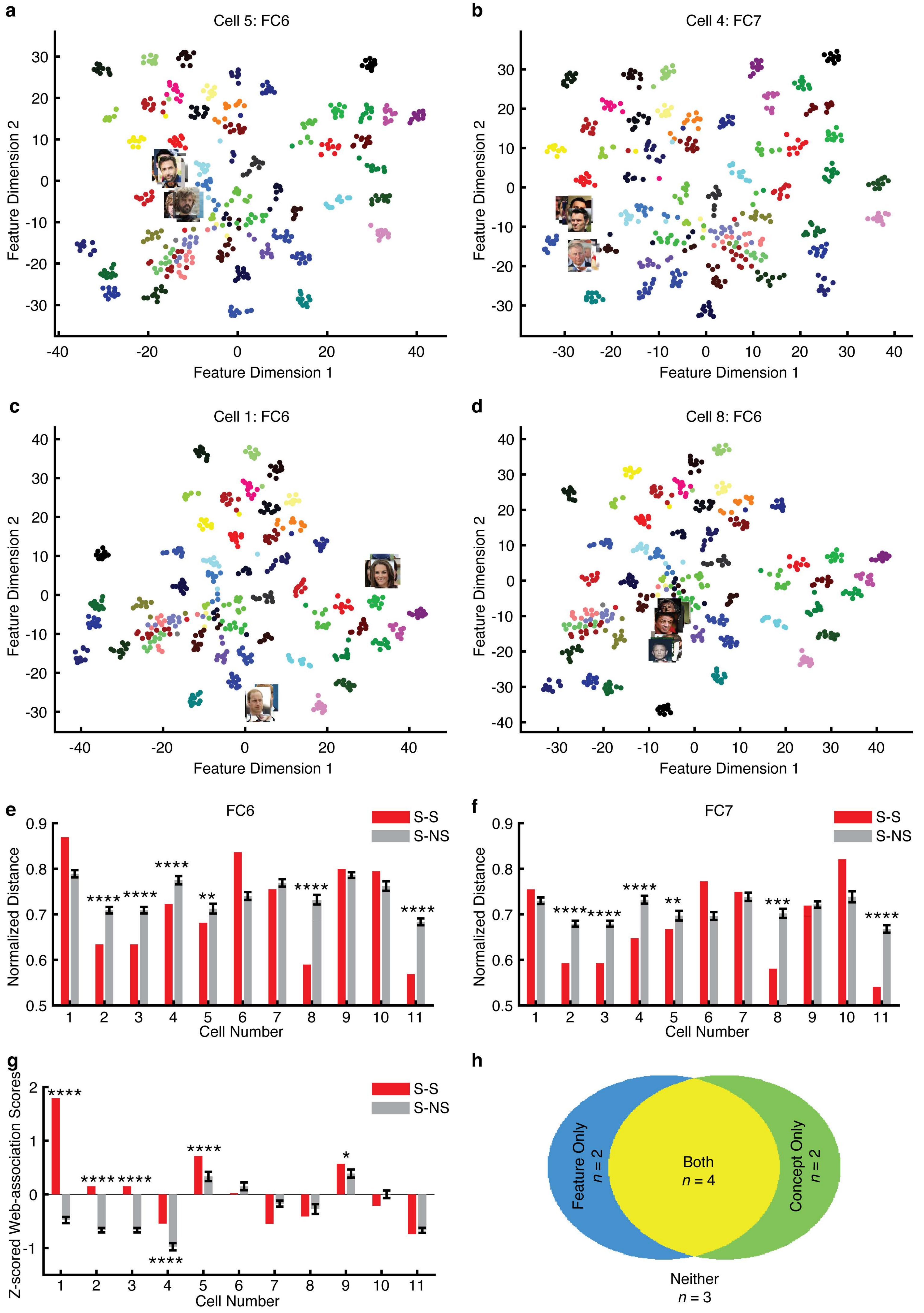
a
Cell 5: FC6
b
Cell 4: FC7
c
Cell 1: FC6
d
Cell 8: FC6
Feature Dimension 1
Feature Dimension 2
e
FC6
f
FC7
S-S
S-NS
Normalized Distance
Cell Number
g
Z-scored Web-association Scores
h
Feature Only
n = 2
Both
n = 4
Concept Only
n = 2
Neither
n = 3

### Figure S15

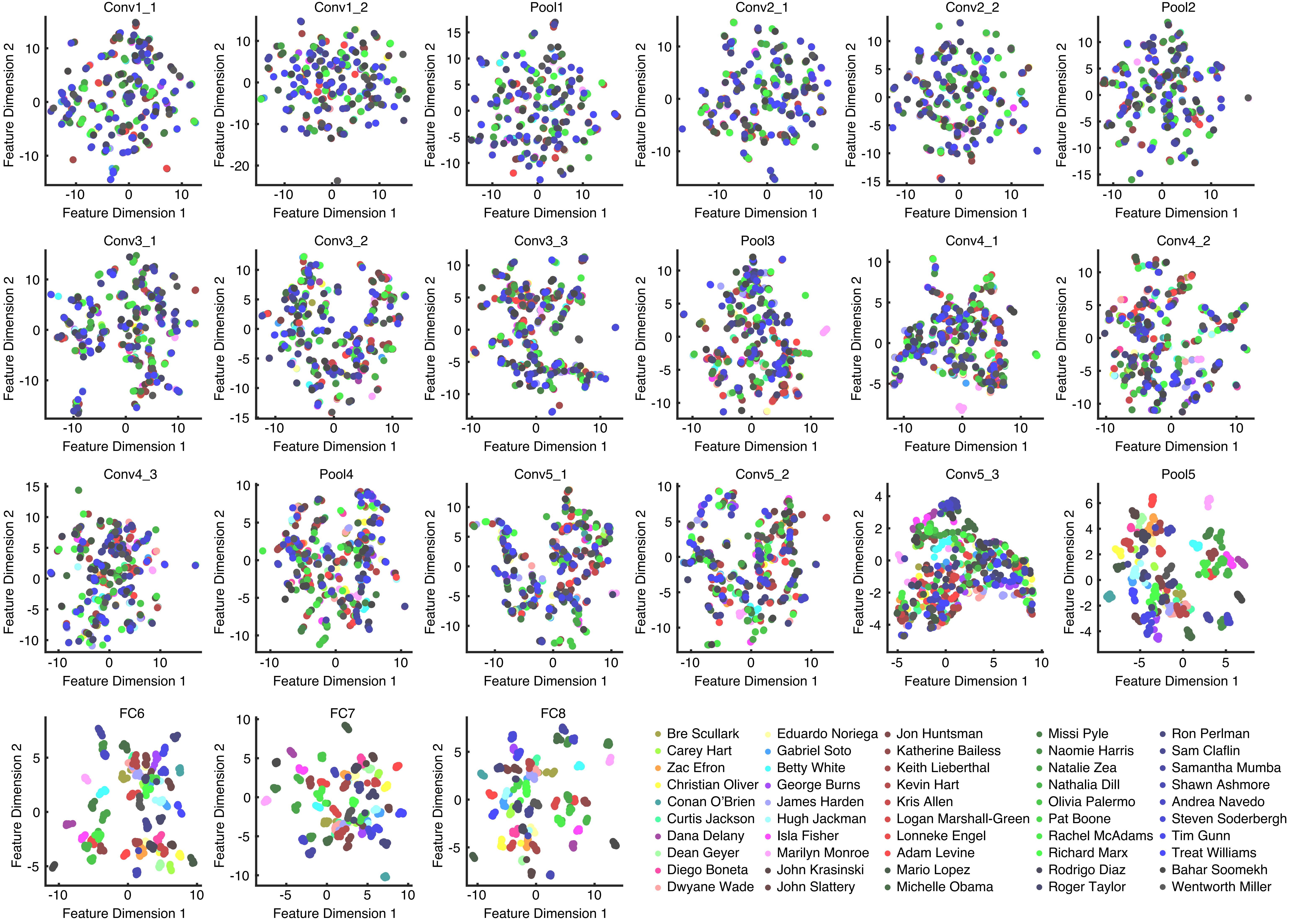
Conv1_1
Conv1_2
Pool1
Conv2_1
Conv2_2
Pool2
Conv3_1
Conv3_2
Conv3_3
Pool3
Conv4_1
Conv4_2
Conv4_3
Pool4
Conv5_1
Conv5_2
Conv5_3
Pool5
FC6
FC7
FC8
Feature Dimension 1
Feature Dimension 2
Bre Scullark
Carey Hart
Zac Efron
Christian Oliver
Conan O'Brien
Curtis Jackson
Dana Delany
Dean Geyer
Diego Boneta
Dwyane Wade
Eduardo Noriega
Gabriel Soto
Betty White
George Burns
James Harden
Hugh Jackman
Isla Fisher
Marilyn Monroe
John Krasinski
John Slattery
Jon Huntsman
Katherine Bailess
Keith Lieberthal
Kevin Hart
Kris Allen
Logan Marshall-Green
Lonneke Engel
Adam Levine
Mario Lopez
Michelle Obama
Missi Pyle
Naomie Harris
Natalie Zea
Nathalia Dill
Olivia Palermo
Pat Boone
Rachel McAdams
Richard Marx
Rodrigo Diaz
Roger Taylor
Ron Perlman
Sam Claflin
Samantha Mumba
Shawn Ashmore
Andrea Navedo
Steven Soderbergh
Tim Gunn
Treat Williams
Bahar Soomekh
Wentworth Miller

Figure S16

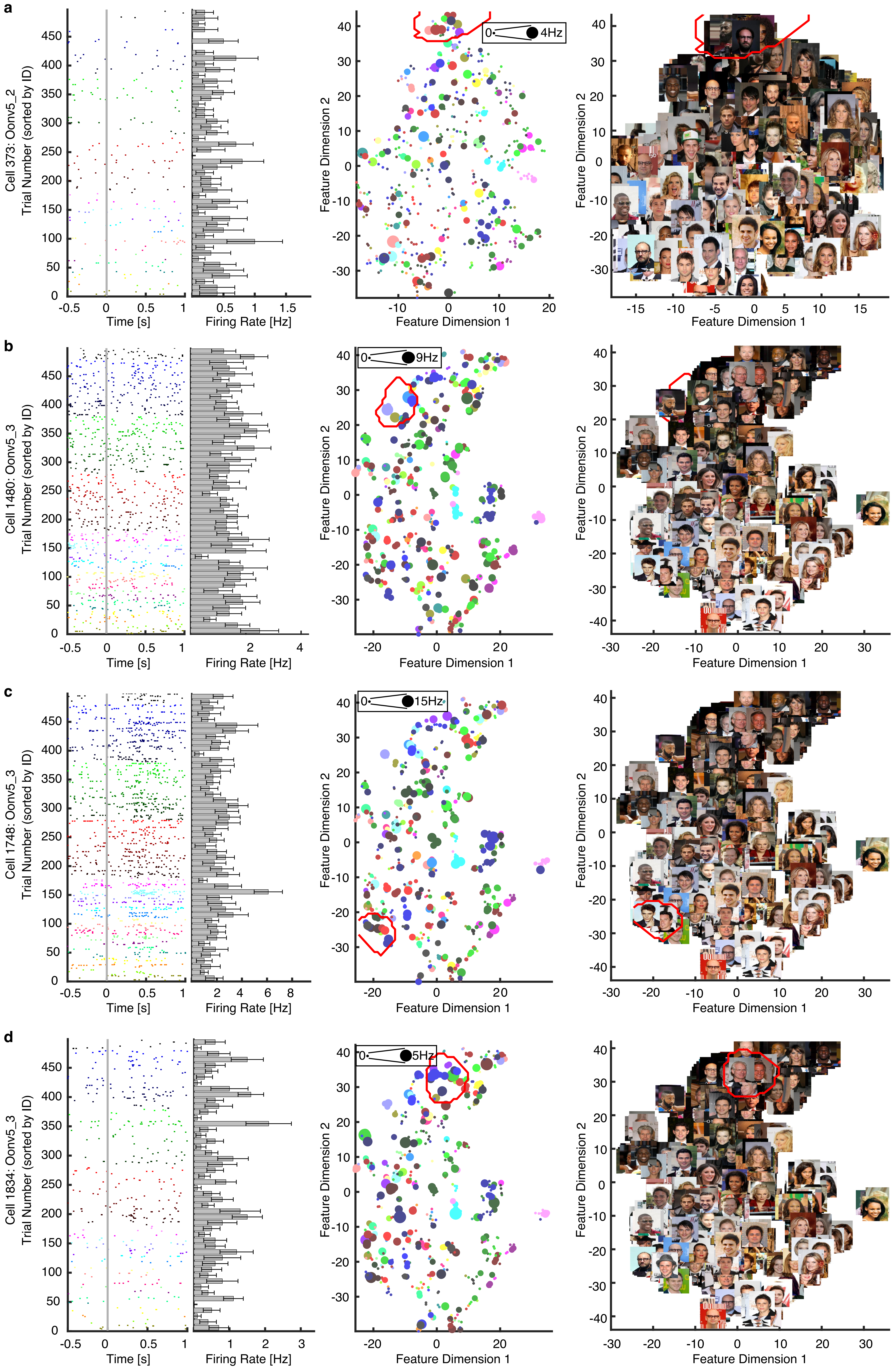
a
b
c
d
Cell 373: Oonv5_2
Cell 1480: Oonv5_3
Cell 1748: Oonv5_3
Cell 1834: Oonv5_3
Trial Number (sorted by ID)
Time [s]
Firing Rate [Hz]
Feature Dimension 1
Feature Dimension 2
0
4Hz
9Hz
15Hz
5Hz

**Figure S17**

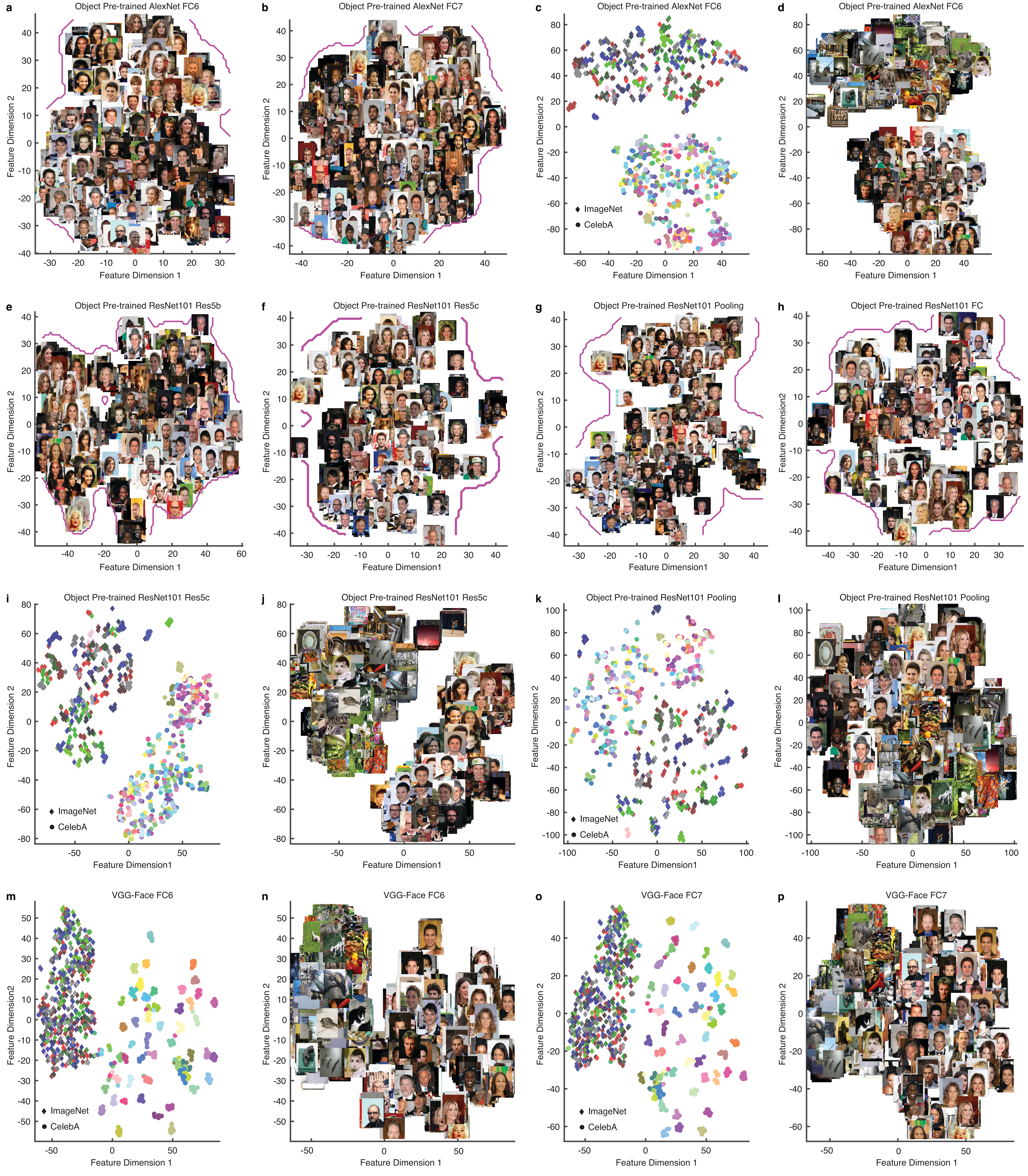
a
Object Pre-trained AlexNet FC6
b
Object Pre-trained AlexNet FC7
c
Object Pre-trained AlexNet FC6
d
Object Pre-trained AlexNet FC6
e
Object Pre-trained ResNet101 Res5b
f
Object Pre-trained ResNet101 Res5c
g
Object Pre-trained ResNet101 Pooling
h
Object Pre-trained ResNet101 FC
i
Object Pre-trained ResNet101 Res5c
j
Object Pre-trained ResNet101 Res5c
k
Object Pre-trained ResNet101 Pooling
l
Object Pre-trained ResNet101 Pooling
m
VGG-Face FC6
n
VGG-Face FC6
o
VGG-Face FC7
p
VGG-Face FC7
Feature Dimension 1
Feature Dimension 2
Feature Dimension1
Feature Dimension2
ImageNet
CelebA

**Figure S18**

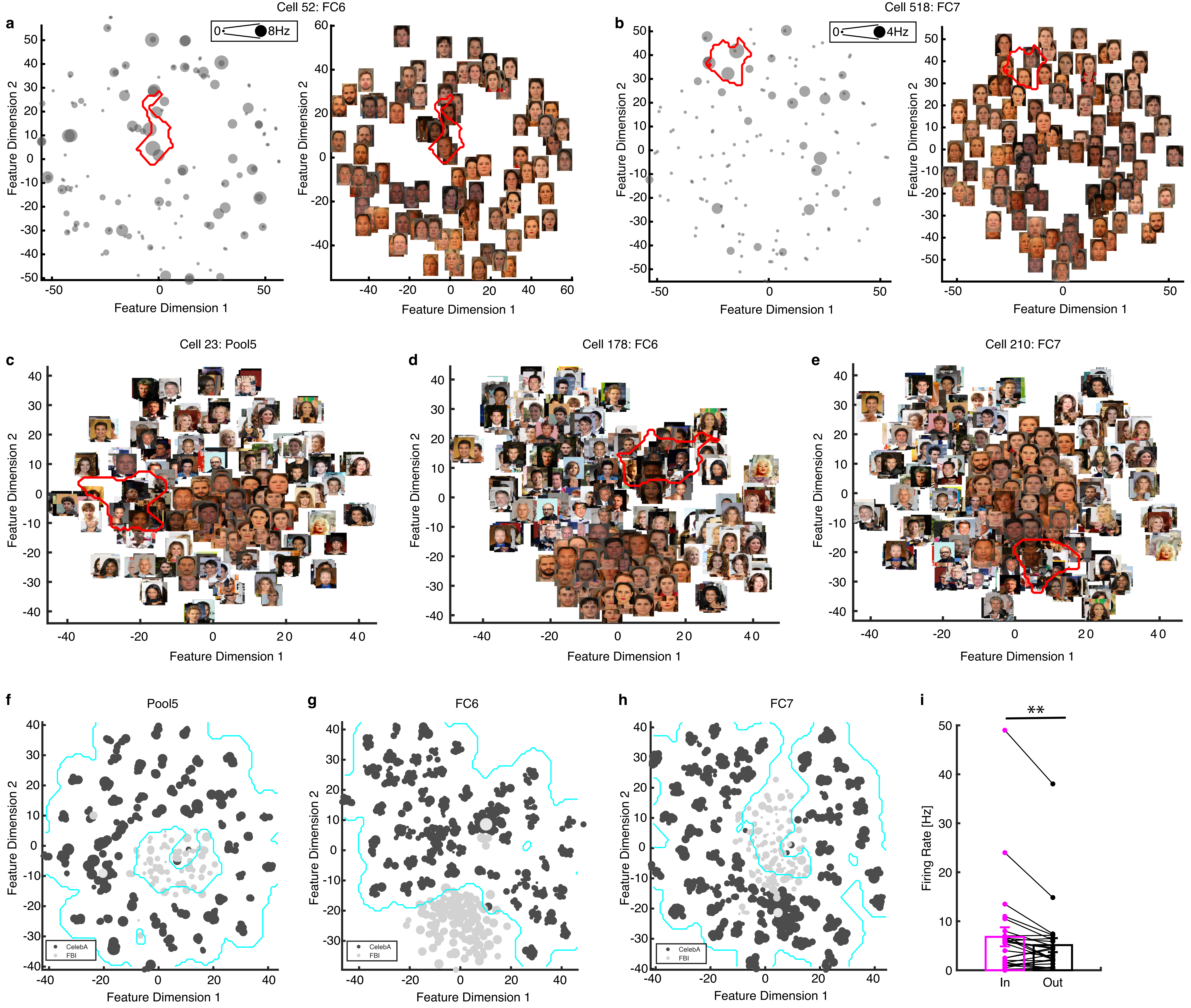
Cell 52: FC6
a
0
8Hz
Feature Dimension 1
Feature Dimension 2
Cell 518: FC7
b
0
4Hz
Cell 23: Pool5
c
Cell 178: FC6
d
Cell 210: FC7
e
f
Pool5
g
FC6
h
FC7
CelebA
FBI
i
**
Firing Rate [Hz]
In
Out

#### Figure S19

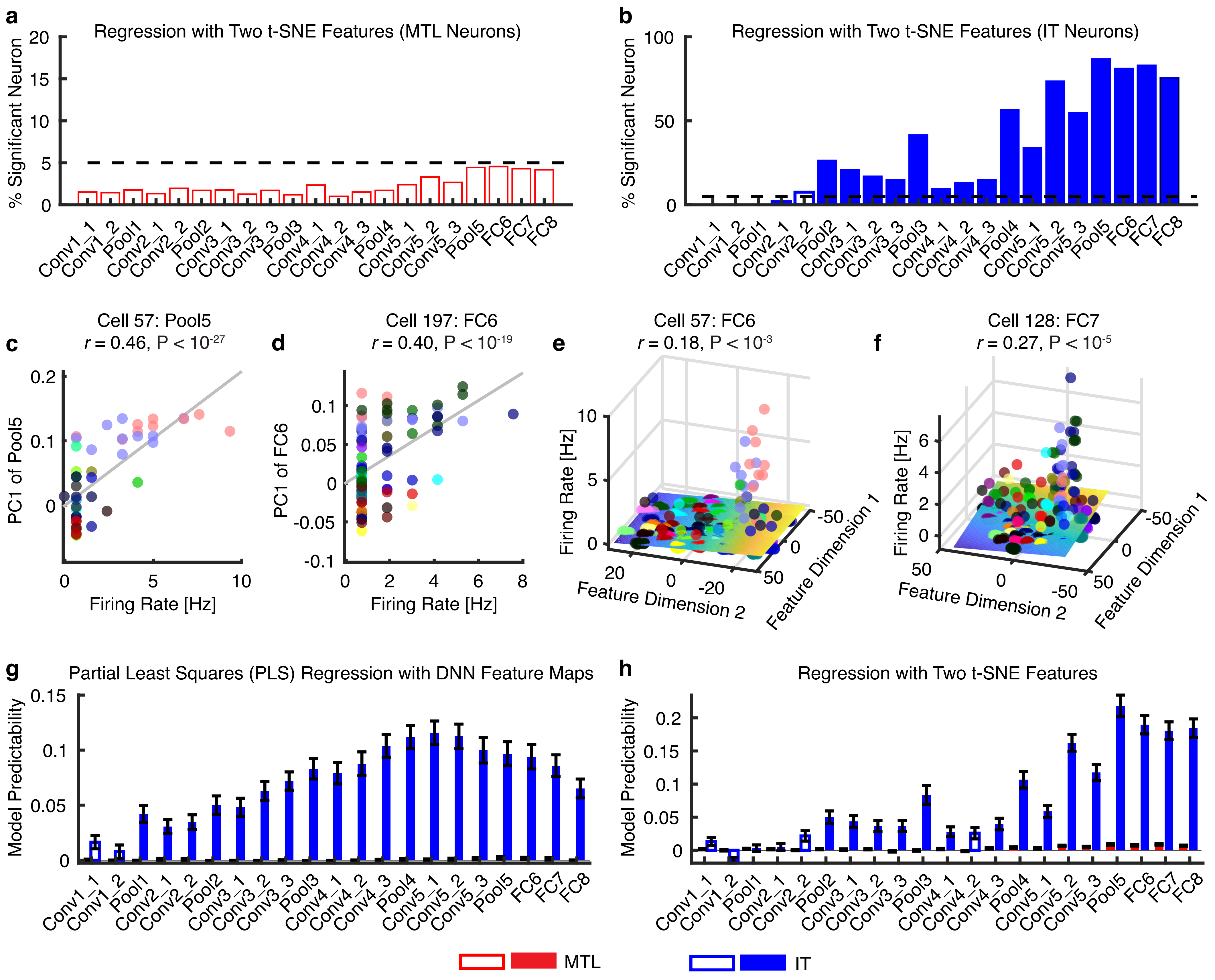

a
Regression with Two t-SNE Features (MTL Neurons)
% Significant Neuron
b
Regression with Two t-SNE Features (IT Neurons)
% Significant Neuron
Conv1_1 Conv1_2 Pool1 Conv2_1 Conv2_2 Pool2 Conv3_1 Conv3_2 Conv3_3 Pool3 Conv4_1 Conv4_2 Conv4_3 Pool4 Conv5_1 Conv5_2 Conv5_3 Pool5 FC6 FC7 FC8
c
Cell 57: Pool5
r = 0.46, P < 10-27
PC1 of Pool5
Firing Rate [Hz]
d
Cell 197: FC6
r = 0.40, P < 10-19
PC1 of FC6
Firing Rate [Hz]
e
Cell 57: FC6
r = 0.18, P < 10-3
Firing Rate [Hz]
Feature Dimension 2
Feature Dimension 1
f
Cell 128: FC7
r = 0.27, P < 10-5
Firing Rate [Hz]
Feature Dimension 2
Feature Dimension 1
g
Partial Least Squares (PLS) Regression with DNN Feature Maps
Model Predictability
h
Regression with Two t-SNE Features
Model Predictability
MTL
IT

**Figure S20**

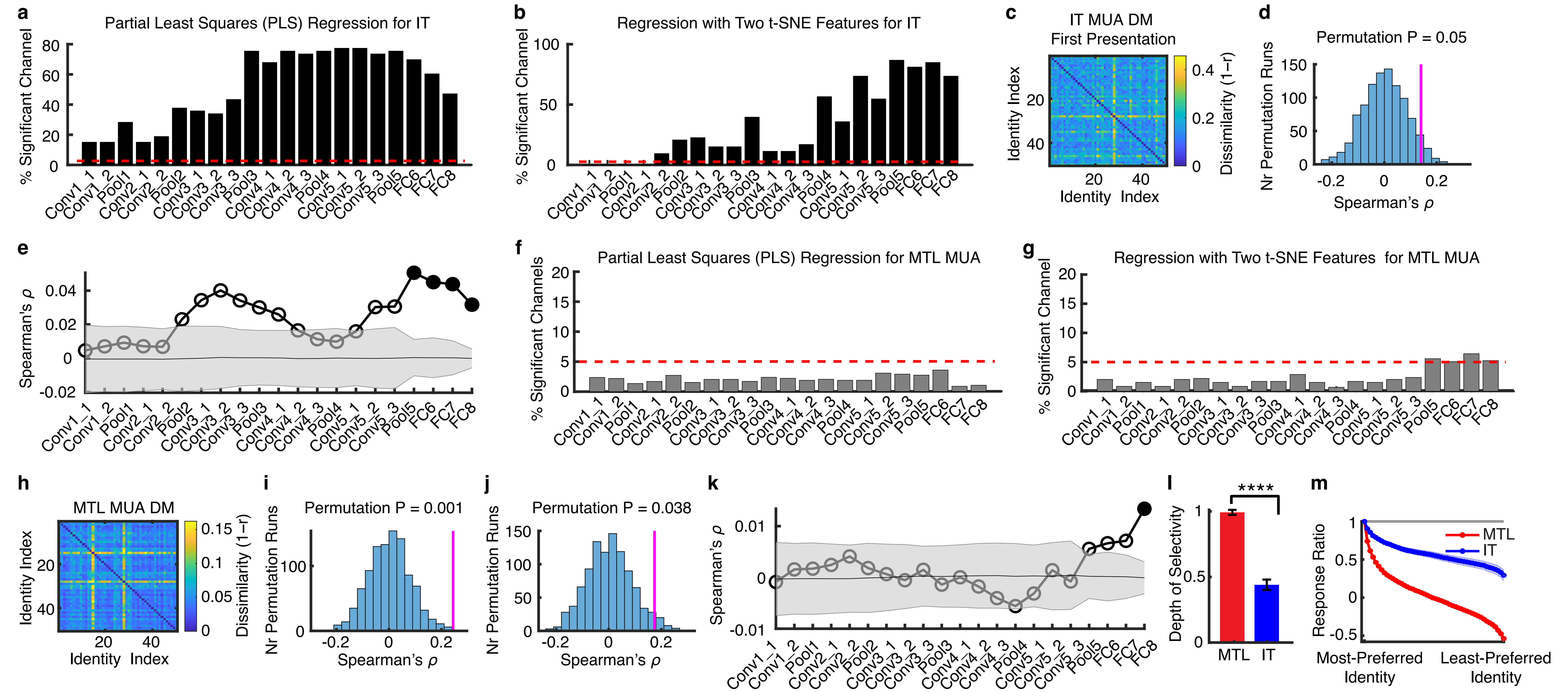
a
Partial Least Squares (PLS) Regression for IT
% Significant Channel
b
Regression with Two t-SNE Features for IT
% Significant Channel
c
IT MUA DM
First Presentation
Identity Index
Dissimilarity (1−r)
d
Permutation P = 0.05
Nr Permutation Runs
Spearman's ρ
e
Spearman's ρ
f
Partial Least Squares (PLS) Regression for MTL MUA
% Significant Channels
g
Regression with Two t-SNE Features for MTL MUA
% Significant Channel
Conv1_1
Conv1_2
Pool1
Conv2_1
Conv2_2
Pool2
Conv3_1
Conv3_2
Conv3_3
Pool3
Conv4_1
Conv4_2
Conv4_3
Pool4
Conv5_1
Conv5_2
Conv5_3
Pool5
FC6
FC7
FC8
h
MTL MUA DM
Identity Index
Dissimilarity (1−r)
i
Permutation P = 0.001
Nr Permutation Runs
Spearman's ρ
j
Permutation P = 0.038
Nr Permutation Runs
Spearman's ρ
k
Spearman's ρ
l
Depth of Selectivity
****
MTL
IT
m
Response Ratio
MTL
IT
Most-Preferred Identity
Least-Preferred Identity
